# Robust and Versatile Arrayed Libraries for Human Genome-Wide CRISPR Activation, Deletion and Silencing

**DOI:** 10.1101/2022.05.25.493370

**Authors:** Jiang-An Yin, Lukas Frick, Manuel C. Scheidmann, Tingting Liu, Chiara Trevisan, Ashutosh Dhingra, Anna Spinelli, Yancheng Wu, Longping Yao, Dalila Laura Vena, Britta Knapp, Elena De Cecco, Kathi Ging, Andrea Armani, Edward Oakeley, Florian Nigsch, Joel Jenzer, Jasmin Haegele, Michal Pikusa, Joachim Täger, Salvador Rodriguez-Nieto, Jingjing Guo, Vangelis Bouris, Rafaela Ribeiro, Federico Baroni, Manmeet Sakshi Bedi, Scott Berry, Marco Losa, Simone Hornemann, Martin Kampmann, Lucas Pelkmans, Dominic Hoepfner, Peter Heutink, Adriano Aguzzi

## Abstract

Arrayed CRISPR libraries extend the scope of gene-perturbation screens but require large numbers of efficacious sgRNA-expressing vectors. Using a newly invented liquid-phase plasmid cloning methodology, we constructed genome-wide arrayed libraries for human gene ablation (19,936 plasmids), activation, and epigenetic silencing (22,442 plasmids). At least 76% of each plasmid preparation encoded an intact array of 4 non-overlapping sgRNAs designed to tolerate most human DNA polymorphisms. We achieved perturbation efficacies of 75-99%, 76-92% and up to 10,000x in deletion, silencing and activation experiments, respectively. Upon conversion into massively parallel lentiviral vectors, an arrayed activation screen of 1,634 human transcription factors yielded 11 novel regulators of the cellular prion protein PrP^C^. Furthermore, a screen using a pooled version of the ablation library identified 5 novel modifiers of autophagy that went undetected with either of two 1sgRNA libraries. The CRISPR libraries described here represent a powerful resource for the targeted perturbation of human protein-coding genes.

## Introduction

According to Karl Popper, fundamentally new discoveries cannot be rooted in prior knowledge (Popper, 1992). A powerful strategy to circumvent this limitation is to perform experiments that do not rely on priors. Unbiased genetic screens, whose development Popper did not live to see, fulfill this requirement. In the last decades, RNA interference or mutagen-mediated screenings have greatly improved our understanding of biology and human health and have transformed drug development for diseases (Mohr et al., 2014; Moresco et al., 2013). CRISPR-mediated techniques have enormously expanded the toolkits of genetic screening, and now allow for gene ablation (CRISPRko), activation (CRISPRa), interference (CRISPRi) and epigenetic silencing (CRISPRoff) (Kampmann, 2020; Liu et al., 2022). CRISPR-based screenings yield both overlapping and distinct hits compared to RNA-interference-based screenings, and CRISPR-mediated gene perturbations are more specific than RNA interference (Evers et al., 2016; Gilbert et al., 2014; Morgens et al., 2016; Smith et al., 2017). Thus, genome-wide CRISPR-based gene perturbation libraries are of essential importance to identify and understand the underlying biology of genes involved in fundamental biological processes and pathological conditions leading to diseases.

Genome-wide CRISPR-based pooled libraries including CRISPRko, CRISPRa and CRISPRi (Hart et al., 2017; Horlbeck et al., 2016; Konermann et al., 2015; Sanson et al., 2018; Shalem et al., 2014) have been deployed in screens for disparate cellular phenotypes (Kampmann, 2020). However, pooled screens cannot be applied to cell-non-autonomous phenotypes, in which genetically mutant cells cause other cells to exhibit a mutant phenotype, and are therefore unsuitable for the study of cell-cell (e.g. glia-neuron) interactions (Araque and Navarrete, 2010). Furthermore, pooled libraries perform suboptimally in genome-wide high-content optical screens and in secretome screens (Feldman et al., 2019; Kampmann, 2020; Kanfer et al., 2021; Yan et al., 2021).

Arrayed sgRNA libraries, which target genes one-by-one in distinct wells, may uncover novel biology through the combination of multiparametric readouts. However, there are only few commercial (HorizonDiscovery; Synthego; ThermoFisher) and academic (Erard et al., 2017; Metzakopian et al., 2017; Schmidt et al., 2015) arrayed libraries and their effectiveness is poorly documented. Moreover, current arrayed libraries suffer from several limitations. Firstly, synthetic crRNA libraries are limited to easily transfectable cells and do not allow for selection of transfected cells. Secondly, plasmid-based libraries featuring one sgRNA per vector exhibit low and heterogenous gene perturbation efficiency (Chakrabarti et al., 2019). Thirdly, single sgRNAs driven by a single promoter may not be effective in the cell types of interest. Fourthly, the sgRNA design algorithms employed by most existing libraries are based only on the hg38 or earlier versions of human reference genomes (Hanna and Doench, 2020; Sanson et al., 2018). However, the genome of cells used for gene-perturbation screens, such as patient-derived human induced pluripotent stem cells (hiPSCs) (Lessard et al., 2017) may diverge from the reference genome, leading to impaired sgRNA function.

For the above reasons, we decided to construct a new generation of highly active, robust and versatile CRISPR arrayed libraries with relatively small size to overcome shared limitations of existing libraries, thereby enabling the investigation of biological space that cannot yet be addressed. The use of multiple sgRNAs targeting each gene improves the potency of gene perturbation applications (Chavez et al., 2016; Erard et al., 2017; McCarty et al., 2020). It was suggested that four sgRNAs/gene achieves saturation gene perturbation effect (Sanson et al., 2018), but the efficiency of 4sgRNA onto gene perturbation has not been tested yet. Moreover, traditional cloning methods (McCarty et al., 2020) relying on gel purification, colony picking and Sanger sequencing are unsuitable for the high-throughput generation of arrayed 4sgRNA plasmids. We have developed the APPEAL method (Automated-liquid-Phase-Plasmid-assEmbly-And-cLoning) which assembles four sgRNAs targeting the same gene, driven by four different promoters, into a single vector cost-effectively. We developed a custom sgRNA design algorithm to select highly specific non-overlapping sgRNAs optimized to tolerate most common polymorphisms. With these methods, we constructed a deletion and a combined activation/silencing library (termed T.spiezzo and T.gonfio) consisting of 19,936 and 22,442 plasmids and targeting 19,820 and 19,839 human protein-coding genes, respectively. Using several paradigms, we show that these libraries can be usefully deployed for largescale gene activation, silencing and ablation screens. We identified several novel transcription factors regulating the expression of the prion protein PrP^C^ and modifiers of autophagy that could not be detected with existing libraries.

## Results

### The APPEAL Cloning Method

Traditional cloning procedures yield heterogeneous recombination products, of which a variable fraction reflects the desired output. This limitation requires the isolation and verification of clonal bacterial colonies streaked on semisolid culture media, which is not easily automatable. This reduces the throughput necessary for the simultaneous generation of large numbers of plasmids. Here we describe the APPEAL procedure (Automated-liquid-Phase-Plasmid-assEmbly-And-cLoning) which allows for one-pot plasmid assembly, transformation and bacterial selection. Thanks to a dual sequential antibiotic selection in the precursor vector (ampicillin) and the final plasmid (trimethoprim), APPEAL only allows for the growth of bacteria harboring the desired plasmids.

We used APPEAL to assemble four sgRNAs, each followed by a distinct variant of tracrRNA and driven by a different ubiquitously active Type-III RNA polymerase promoter (human U6, mouse U6, human H1, and human 7SK) into a single vector (**Fig. 1A** and **S1**) (Adamson et al., 2016; Kabadi et al., 2014; Sanson et al., 2018). The 4sgRNA vector includes puromycin and TagBFP cassettes flanked by lentiviral long terminal repeats and PiggyBac (PB) transposon elements enabling multiple routes of selection and transduction (Metzakopian et al., 2017) (**Fig. 1A**). The four sgRNAs were individually synthesized as 59-meric oligonucleotide primers comprising the 20-nucleotide protospacer sequence and a constant region including amplification primer annealing sites (**Fig. S1A**). In three distinct polymerase chain reactions (PCR), the primers were mixed with the corresponding constant-fragment templates to produce three individual amplicons. These amplicons and the digested empty vector (pYJA5) contain directionally distinct overlapping ends (approximately 20 nucleotides) enabling Gibson assembly (Gibson et al., 2009) (**Fig. 1A** and **S1**).

**Fig. 1.**
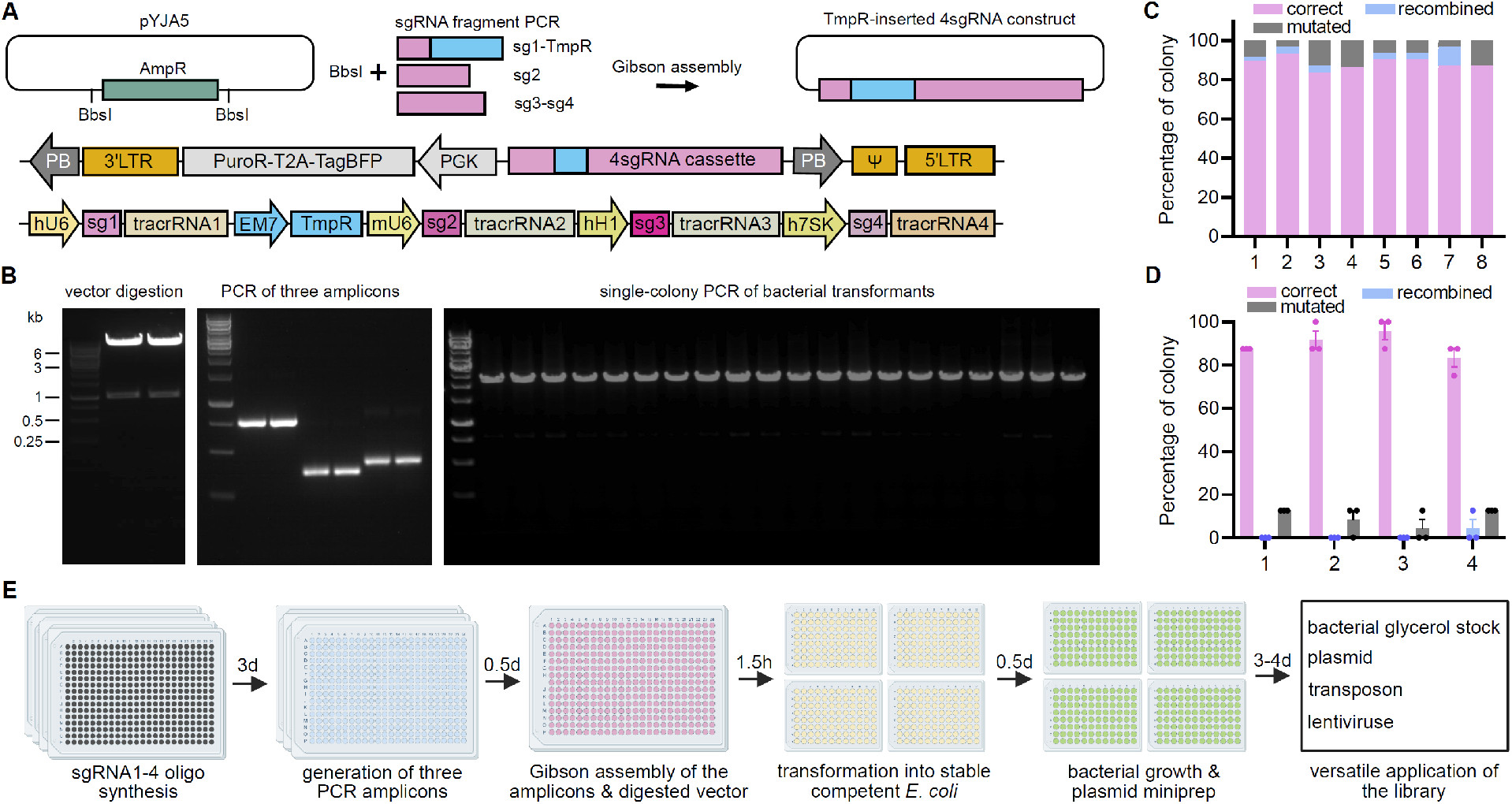
APPEAL cloning. **A**, The ampicillin resistance gene (AmpR) was removed from the vector pYJA5. sgRNA1-4 and the trimethoprim resistance gene (TmpR) was fused with three distinct PCR amplicons. All elements were Gibson- assembled to form the 4sgRNA-pYJA5 plasmid, and transformants were selected with trimethoprim. The detailed structure of 4sgRNA-pYJA5 full plasmid and 4sgRNA cassette were depicted. LTR, long terminal repeat; Ψ, packaging signal sequence; PB, piggyBac transposon element; PuroR, puromycin resistance element; hU6, mU6, hH1 and h7SK are ubiquitously expressed RNA polymerase-III promoters; sg, sgRNA. **B**, Representative pYJA5 restriction fragments, 3-fragment PCRs, and single-colony PCR of APPEAL cloning products after transforming into *E. coli* and trimethoprim selection. *Bbs*I digestion of pYJA5 yielded ∼1–kilobase (kb) band of the AmpR element and ∼7.6-kb band of the line- arized vector (left). After PCR with corresponding sgRNA primers, the three amplicons showed the expected size of 761, 360 and 422 bp on agarose gels, respectively (middle). Single-colony PCR with primers flanking the 4sgRNA cassette of APPEAL cloning products in transformed bacteria plate consistently yielded the expected size (2.2 kb, right). **C**, Percentage of correct, recombined, and mutated 4sgRNA plasmids in 8 independent APPEAL experiments with distinct 4sgRNA sequences (≥22 colonies were tested in each experiment). **D**, Percentage of correct, recombined, and mutated 4sgRNA plasmids in four APPEAL experiments. Each dot represented an independent biological replica consisting of eight colonies (n=24; Mean ± S.E.M.). **E**, Timeline of APPEAL cloning in high-throughput format (h: hours; d: days).

In the precursor vector pYJA5, the β-lactamase gene (AmpR) providing ampicillin resistance was flanked by two *Bbs*I restriction sites and was removed to minimize the size of final 4sgRNA plasmids for the subsequent Gibson assembly steps. A trimethoprim-resistant dihydrofolate reductase gene (TmpR) was incorporated into the first amplicon between sgRNA1 and sgRNA2, enabling a selection switch from ampicillin to trimethoprim **(Fig. 1A** and **S1**) (Adamson et al., 2016; McCarty et al., 2020). To test the accuracy of APPEAL, we cloned the sgRNA-containing amplicons into pYJA5 using Gibson assembly (**Fig. 1B**) and performed PCR on single colonies with primers flanking the 4sgRNA insert. All PCR products from the tested colonies showed the expected size of 2.2 kilobases, suggesting the correct assembly of all three fragments and backbone (**Fig. 1B**). We then sequenced single colonies from eight independent cloning procedures (≥22 colonies/procedure). All colonies showed the desired antibiotic selection switch, and each procedure resulted in 83%-93% of colonies showing correct 4sgRNA sequences (**Fig. 1C**).

Repetitive DNA sequences may lead to inappropriate recombination. To minimize this effect, each of the 4 sgRNA is driven by a different Pol-III promoter and followed by a distinct tracer RNA. In the eight cloning trials described above, 0–10% of tested colonies harbored recombined plasmids (**Fig. 1C**). Furthermore, cloned plasmids may harbor point mutations. The frequency of mutated plasmids in each of the cloning trials was 3–14% (**Fig. 1C**). Such mutations may result from errors in oligonucleotide synthesis and the tolerance of DNA mismatches by the Taq DNA ligase during Gibson assembly (Gibson et al., 2009; Lohman et al., 2016). Four of the eight cloning trials were repeated three times; the percentages of colonies with correct, recombined, or mutated plasmids were similar to the previous trials (**Fig. 1D**). Hence, APPEAL improves the selection of desired end products and generates high-quality plasmids without requiring the isolation of single bacterial colonies. For highthroughput cloning, APPEAL steps were performed in 384-well plates. Reaction products were transferred to deep-96-well plates for transformation and amplification in recombination-deficient chemically competent *E. coli*. Magnetic bead-based plasmid minipreps were performed in the same microplates using custom-made equipment, enabling the construction of >42,000 individual plasmids (∼2,000 plasmids/week with two full-time equivalents) with a yield of ∼25 µg/plasmid (**Fig. 1E**).

### Superiority and Robustness of 4sgRNAs in Gene Activation and Ablation

As a proof-of-concept, we tested the efficiency of APPEAL-cloned 4sgRNA plasmids for gene activation and ablation in HEK293 (human embryotic kidney) cells. For CRISPRa, we co-transfected the CRISPR activator dCas9-VPR with sgRNA-expressing plasmids targeting the genes *ASCL1, NEUROD1* and *CXCR4* which have low, moderate and high baseline expression respectively(Chavez et al., 2016). Compared to individual sgRNAs cloned into a pYJA5-derived vector modified for one sgRNA insertion, 4sgRNA vectors massively increased target gene activation (**Fig. 2A**), consistent with a synergistic gene activation efficiency by three sgRNAs (Chavez et al., 2016). We then tested genes (*HBG1*, *KLF4*, *POU5F1*, *ZFP42*, *IL1R2*, *MYOD1*, *TINCR*, and the long non-coding RNAs *LIN28A*, *LINC00925*, *LINC00514*, and *LINC00028*) that proved difficult to activate before the invention of the synergistic activation mediator (SAM) (Konermann et al., 2013; Mali et al., 2013; Perez-Pinera et al., 2013). All tested genes were successfully activated by 4sgRNA plasmids (**Fig. 2B**). Furthermore, we used the cell-surface proteins CD2, CD4, and CD200 in HEK293 cells to test if the 4sgRNA design might reduce the considerable cell-to-cell heterogeneity afflicting gene-activating single sgRNAs (Chong et al., 2018). Single sgRNAs induced variable, mostly low gene activation, whereas 4sgRNA transduction into HEK293 cells stably expressing doxycycline-inducible dCas9- VPR led to robust cell-surface expression of CD2, CD4 and CD200 with improved separation of activated cells from the non-targeting controls, as reflected by superior Z’ factors (**Fig. 2C** and **S2A**). We assessed the CRISPRko efficacy by live-cell antibody staining of the cell-surface molecules CD47, IFNGR1 and MCAM (Bausch-Fluck et al., 2015). For each gene, 12 single sgRNAs from widely used resources (Hart et al., 2017; Sanjana et al., 2014; Sanson et al., 2018) were tested.

**Fig. 2.**
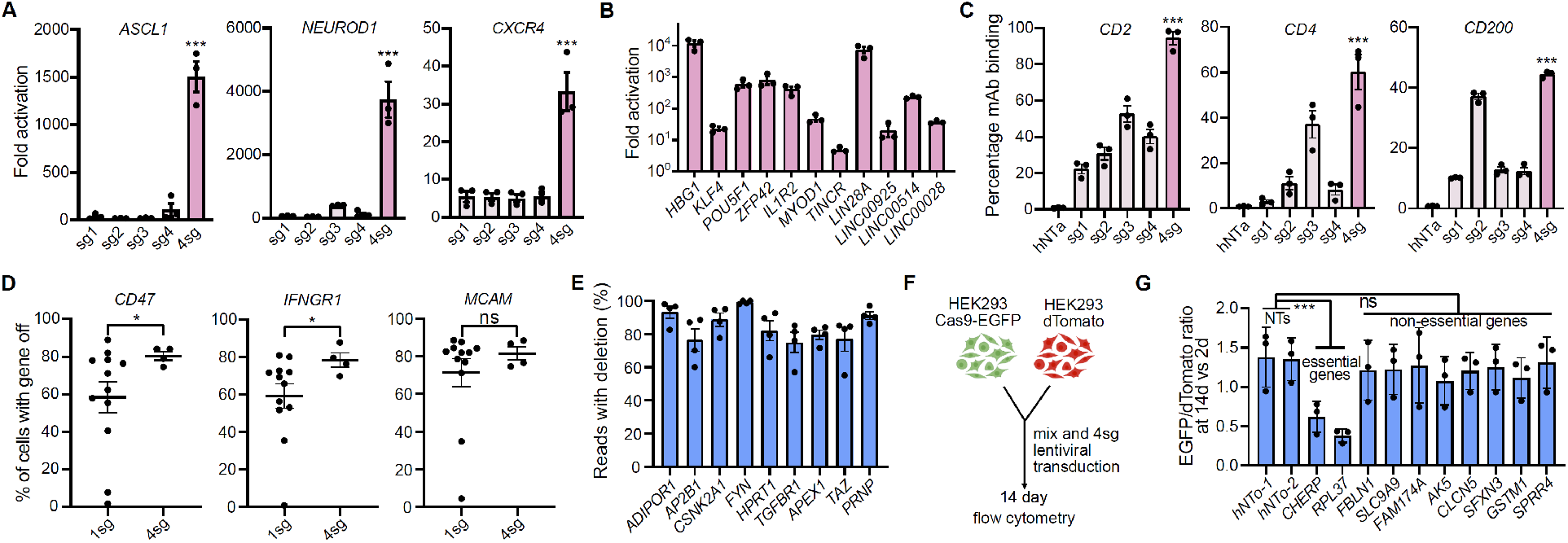
Gene activation and ablation with 4sgRNA. **A**, Gene activation (qRT-PCR) in HEK293 cells 3 days post transfection with dCas9-VPR and single (sg1-4) or four sgRNA (4sg) plasmids. Dots (here and henceforth except **D**): independent experiments (Mean ± S.E.M.).. **B**, Gene activation by 4sgRNA plasmids co-transfected with dCas9-VPR into HEK293 cells (3 days post transfection). **C**, Surface expression of CD2, CD4, and CD200 in HEK293 cells after gene activation with single or four sgRNAs, hNTa, non-targeting control plasmid; Cell surface proteins were stained with fluorescent-conjugated antibodies and analyzed via flow cytometry. **D**, Gene disruption efficiency by single sgRNAs vs 4sgRNAs in HEK293 cells via co-transfection with Cas9 plasmid. 12 single sgRNAs (1sg) from the Brunello, GeCKOv2 and TKOv3 libraries were tested; 4sgRNA (4sg) plasmids were assembled with 4 random or the 4 worst- performed single sgRNAs among the 12 sgRNAs tested; Each dot represents the mean of three independent biological repeats (Fig. S2B); hNTo, non-targeting control plasmid; Assay were performed similarly as in **C**. **E**, Robust ablation of genes inadequately disrupted by single sgRNAs(de Groot et al., 2018). Single sgRNAs were assembled into 4sgRNA plasmids and co-transfected with Cas9 plasmid into HEK293 cells (as in **D**). Gene disruption was quantified by single-molecule real-time long-read sequencing of the genomic region covering all sgRNAs target sites. **F**, EGFP and Cas9-expressing HEK293 cells (HEK293-Cas9-EGFP) were mixed with dTomato-expressing HEK293 (HEK293- dTomato) cells (∼1:1 ratio) and transduced with 4sgRNA lentiviruses. **G**, EGFP/dTomato ratio in a HEK293-Cas9- EGFP and HEK293-dTomato co-culture two weeks after transduction of 4sgRNA lentiviruses. 4sgRNA targeting essential genes, non-essential genes, or nontargeting controls (NTs) were tested. EGFP/dTomato ratio at end point (day 14) was normalized to the ratio at day 2 post transduction. In **A** and **C**, p values were determined by Dunnett’s test, whereas in **D** and **G**, two-tailed Student’s t-test was applied. Here and henceforth: *: p<0.05; **: p<0.01; ***: p<0.001; ns: not significant.

Single sgRNAs showed variable ablation efficiencies (5%-85% for *CD47*, 1%-76% for *IFNGR1*, and 6%-85% for *MCAM*). In contrast, the respective 4sgRNA plasmids showed ablation efficiencies of >70% (**Fig. 2D** and **S2B)**. To further test the efficiency and robustness of the 4sgRNA approach for gene ablation, we examined 9 genes (*ADIPOR1*, *AP2B1*, *CSNK2A1*, *FYN*, *HPRT1*, *TGFBR1*, *APEX1*, *TAZ*, and *PRNP*) for which single sgRNAs have low or moderate ablation efficiency (de Groot et al., 2018). Single molecule real-time (SMRT) long-read sequencing of PCR amplicons from genome-edited cells showed nucleotide deletions in 75%-99% of the sequencing reads (**Fig. 2E**). Agarose-gel electrophoresis showed conspicuous deletions of the genomic region between the sgRNA cutting sites for all nine genes (**Fig. S2C**).

Gene ablation with 4sgRNA plasmids may induce multiple DNA double-strand breaks, resulting in toxicity. We tested this by transducing equal numbers of dTomato-expressing wild-type and EGFP- Cas9-expressing HEK293 cells (**Fig. 2F**) with nontargeting 4sgRNAs (hNTo-1 and hNTo-2) or 4sgRNAs against essential (*CHERP* and *RPL37*) or non-essential (*FBLN1, SLC9A9, FAM174A, AK5, CLCN5, SFXN3, GSTM1* and *SPRR4*) genes (**Fig. 2G**). After two weeks in culture, transduction with 4sgRNAs targeting essential genes led to the profound depletion of EGFP. However, no EGFP depletion was observed in cells transduced with nontargeting 4sgRNAs or with 4sgRNA plasmids targeting non-essential genes (**Fig. 2G**). We conclude that 4sgRNA knockout plasmids do not impose detectable toxicity onto genome-edited cells.

Together, these results demonstrate the high efficacy and robustness of our 4sgRNA/gene strategy for both gene activation and ablation.

### Lentiviral Packaging and Delivery of the 4sgRNA Vectors

We established a robust lentiviral production protocol in 384-well plates in HEK293T cells (**Fig. S2D**). With this method we can convert 2000-4000 4sgRNA plasmids per week into lentiviruses with a median titer of ∼ 5x10^6^ lentiviral transducing units per milliliter (TU/ml) in raw culture-medium supernatants (**Fig. S2D** and **S2E**). These viruses efficiently delivered sgRNA to the human lymphocyte-related cell lines THP-1 and ARH-77, the human neuroblastoma cell line GIMEN, the human glioblastoma cell line U251-MG, and patient-derived iPSCs, as indicated by the fraction of tagBFP^+^ cells measured by flow cytometry (**Fig. S2F**). We then examined the efficiency of gene activation in iPSC-derived neurons (iNeurons, which stably express dCas9-VPR) using lentivirus-mediated delivery of the 4sgRNA vector. Gene activation was generally efficient, and its extent depended on the basal expression levels of the target genes (**Fig. S2G**).

### Updated Algorithms for Generic, Specific and Synergistic sgRNA Selection

To enable gain-of-function and loss-of-function arrayed CRISPR screens, we decided to generate CRISPRa (termed as T.gonfio, meaning swelling up) and CRISPRko (termed as T.spiezzo, meaning sweeping away) arrayed libraries for human protein-coding genes using APPEAL (**Fig. 1E**). We used the Calabrese and hCRISPRa-v2 sgRNA sequences for T.gonfio, and the Brunello and TKOv3 sequences for T.spiezzo (Hart et al., 2017; Horlbeck et al., 2016; Sanson et al., 2018) as a baseline to generate our arrayed libraries with a novel algorithm to select the optimal combination of four sgRNAs (**Fig. 3A**).

**Fig. 3.**
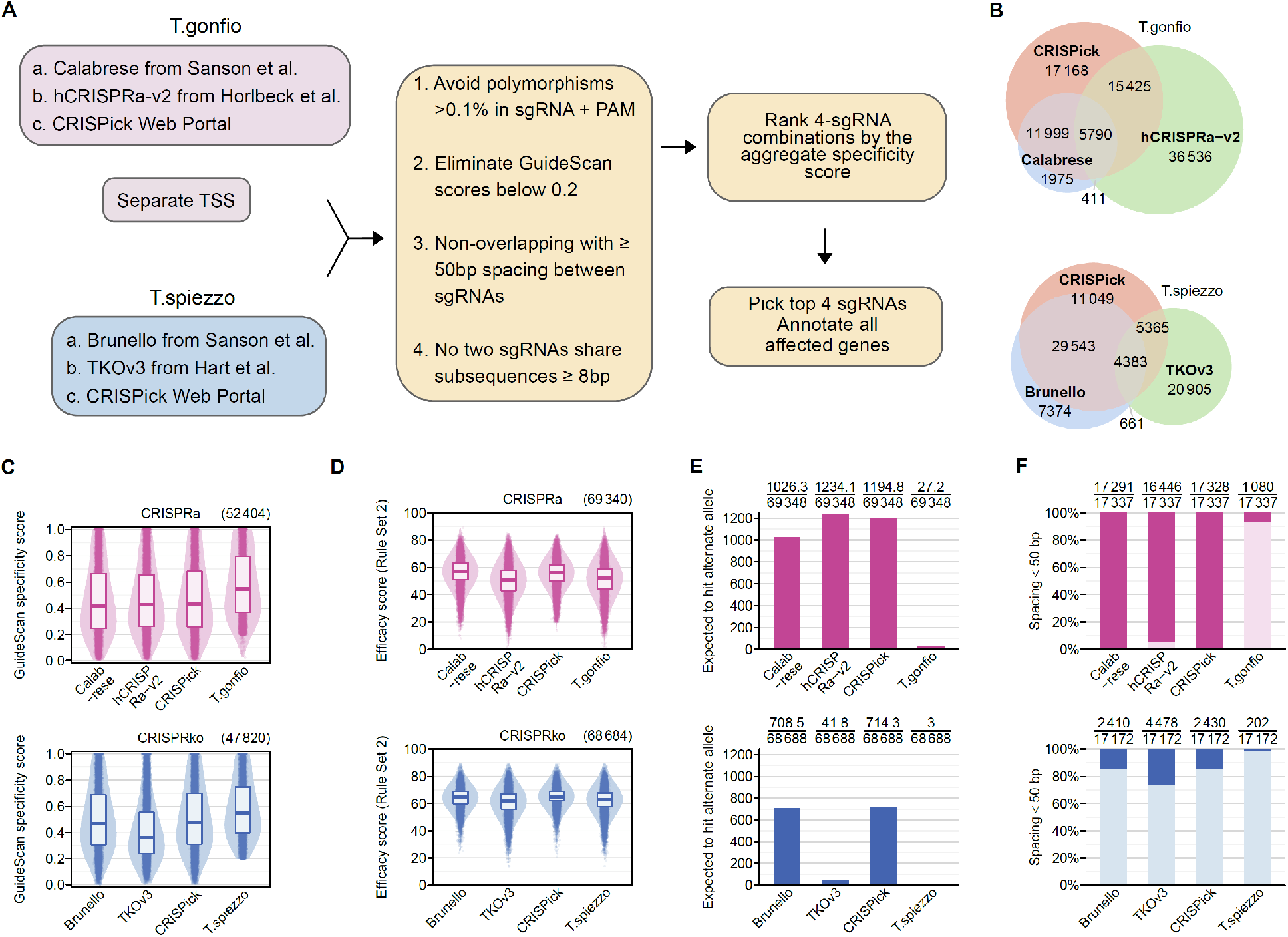
sgRNA selection and features of the T.spiezzo and T.gonfio libraries. **A**, A large pool of potential sgRNAs was explored by combining existing CRISPR libraries and the output of the CRISPick platform. Each sgRNA was annotated with data on genetic polymorphisms affecting the target region and with GuideScan specificity scores. For T.gonfio, individual 4sgRNA plasmids were designed to target each alternative TSS (transcription start site). The best combination of four non-overlapping sgRNAs was chosen (Table S1). **B**, Numbers of sgRNAs of the T.spiezzo and T.gonfio libraries originated from existing libraries and resources. **C**, Cross-library comparison of GuideScan specificity scores for the individual sgRNAs. The top 4 sgRNAs from each library were included, based on their original ranking, for genes that are present in all source libraries. **D**, Cross-library comparison of predicted sgRNA efficacy scores. **E**, Comparison of the number of sgRNAs expected to target alternate alleles of genetically polymorphic regions, based on prevalence in humans. Only polymorphisms with a frequency >0.1% are considered. **F**, Percentage of 4sgRNA combinations where ≥1 pair of sgRNAs is spaced fewer than 50 bp apart, potentially leading to interference.

Common DNA polymorphisms in human genomes, such as single-nucleotide polymorphisms (SNPs), affect 0.1% of the genome and may reduce CRISPR efficacy (Lessard et al., 2017). Except for the TKOv3 library, most CRISPR libraries did not consider DNA polymorphisms when selecting sgRNA sequences, hampering their usage on primary patient-derived cells. We obtained a dataset from over 10’000 complete human genomes (http://db.systemsbiology.net/kaviar/) providing the coordinates of genetic polymorphisms aligned to the hg38 reference genome. Guide RNAs were flagged as unsuitable if the genomic coordinates of the 20-nucleotide protospacer sequence or the two guanine nucleotides of the protospacer adjacent motif (PAM) were affected by a polymorphism with a frequency higher than 0.1%.

The GuideScan algorithm (http://www.guidescan.com) can predict off-target effects of sgRNAs with high accuracy, showing a strong correlation with the unbiased genome-wide off-target assay GUIDESeq (Tsai et al., 2015; Tycko et al., 2019). Guide RNAs with GuideScan scores exceeding 0.2 are generally considered specific (Tycko et al., 2019), and we imposed this constraint for sgRNA selection in our libraries.

Previous libraries chose top-ranking sgRNA sequences based on high on-target efficacy scores and low predicted off-target effects, yet this could result in the selection of overlapping sgRNAs whose target positions differed only by a few nucleotides. This was particularly common in CRISPRa libraries, due to the limited target window for sgRNAs upstream of the transcription start site (TSS). We found that four non-overlapping sgRNAs (spaced at least 50 nucleotides apart) resulted in significantly higher gene activation than 4sgRNA combinations that did not meet this criterion, suggesting that spatially unconstrained binding of sgRNA-dCas9-VPR complexes is strongly synergistic (**Fig. S3A**). Secondly, since we generated our libraries using the Gibson assembly method, if two or more sgRNAs share identical subsequences of 8 nucleotides or more, the prevalence of correct plasmids decreased because of recombination between identical sequences among the four sgRNAs (**Fig. S3B**-**C**).

The efficacy of CRISPRa-mediated gene activation relies on sgRNAs targeting a narrow window of 400 base pairs (bp) upstream of TSS of a gene (Gilbert et al., 2014; Sanson et al., 2018). Many genes have more than one TSS that may exhibit different activity in different cell models (Consortium et al., 2014; Sanson et al., 2018). Therefore, the T.gonfio library targets each major TSS with an individual 4sgRNA plasmid (except for genes whose TSSs were spaced ˂1000 bp apart). Because some genes or TSSs did not have four sgRNAs that fulfilled these requirements, we supplemented the above-mentioned libraries with sgRNAs from the CRISPick web portal (https://portals.broadinstitute.org/gppx/crispick/public), which designs sgRNAs with the same algorithm that was used for the Calabrese and Brunello libraries for CRISPRa and CRISPRko, respectively. Finally, after filtering with the above four constraints, all possible combinations of four sgRNAs targeting a gene/TSS were ranked by their aggregate specificity score, enabling the selection of sgRNA sequences with minimized potential off-target effects (**Fig. 3A**).

### Features of the T.spiezzo and T.gonfio libraries

The T.gonfio and the T.spiezzo libraries include 22,442 plasmids and 19,936 plasmids covering 19,839 and 19,820 human protein-coding genes, respectively. Each library contains 116 non-targeting control plasmids and is organized into thematic sublibraries (**Table 1**). Transcription factor, secretome and G protein-coupled receptor (GPCR) sublibraries were defined according to current gene catalogs (Lambert et al., 2018; Uhlen et al., 2019). Other sublibraries were based on categories defined by the pooled library hCRISPRa-v2 (Horlbeck et al., 2016). The sgRNAs selected with our updated algorithm originated mostly from previously published libraries (**Fig. 3B**). T.gonfio covers 17,528 genes targeted at a single TSS and 2,311 genes targeted at ≥2 TSSs (**Fig. S3D**). Among the 19,820 genes targeted by the T.spiezzo library, the expected deletions in the human genome ranges between 10 and >10^5^ bases (**Fig. S3D**). By excluding sgRNAs with GuideScan scores <0.2, we enriched for specificity without sacrificing the predicted efficacy **(Fig. 3C**-**D**). Both T.spiezzo and T.gonfio were designed to improve targeting by avoiding genetically polymorphic regions (**Fig. 3E** and **S3E**). Our sgRNA selection algorithm maximized the proportion of sgRNAs spaced ≥50 nucleotides apart (**Fig. 3F**), thus increasing targeting efficacy. Furthermore, we avoided 4sgRNA combinations that shared sub-sequences of ≥8 base pairs (**Fig. S3F**), thus ensuring minimal chances for sgRNA recombination.

We avoided sgRNAs with multiple perfect genomic matches wherever possible. However, when targeting families of closely related, paralogous genes, there were often no specific sgRNAs to choose from. For simplicity, we created a separate 4sgRNA plasmid for each protein-coding gene that possessed its own unique Entrez gene identifier. In T.spiezzo, such unspecific sgRNAs were mostly excluded, whereas in T.gonfio the proportion of sgRNAs with off-site targets (0.8%) was comparable to the reference pooled libraries because the window around the TSS constrained the choice of target sites (**Fig. S3G**).

CRISPR activation of unintended genes may also occur if two genes locate on opposite strands of the genome and share a bidirectional promoter region. Such effects are unavoidable; indeed, when considering a window of 1 kb surrounding the TSS, around 20% of CRISPRa sgRNAs affected additional genes in all examined libraries including T.gonfio (**Fig. S3H**). All sgRNAs that affect any genes other than the intended gene have been annotated (**Fig. S3I** and **Supplemental Dataset 1**).

### Sequencing the T.spiezzo and T.gonfio Libraries

To assess the plasmid quality of our libraries, we amplified the 4sgRNA-expression cassettes in each well with barcoded primers and subjected pools of amplicons (2.2-kilobase) to single-molecule real-time (SMRT) sequencing (**Fig. S4A**). To estimate technical errors, 74 single-colony-derived, fully-sequenced plasmids bearing distinct 4sgRNA sequences were included in each sequencing round. The median read count (at CCS7 quality) per plasmid was 86 with ≥10 reads for 98.7% and ≥1 read for 99.9% of plasmids, respectively (**Fig. S4B**).

Mutations, deletions and recombinations may occur in plasmids constructed by Gibson assembly; because APPEAL does not rely on colony-picking, these alterations may affect a fraction of the plasmid pool in each well. This heterogeneity was precisely quantified by single-molecule long-read sequencing of all four promoter, protospacer and tracrRNA sequences. sgRNAs were considered correct if the 20-nucleotide sgRNA and tracrRNA sequences were present and error-free. When considering the median across all wells in the libraries, the percentage of reads with at least one, two, three or four correct sgRNAs was 98%, 94%, 92% and 78% for the T.gonfio library, and 98%, 92%, 89% and 76% for the T.spiezzo library, respectively (**Fig. 4A**). At the 5^th^ percentile (i.e., worse than 95% of wells in the library), the fraction of reads with ≥3 correct sgRNAs was 77% for the T.gonfio library, and 71% for the T.spiezzo library (**Fig. 4B**). 99.7% of wells passed the minimal quality standard of >50% reads with at least one correct sgRNA (76 wells failed to meet this standard, including 38 wells with zero CCS7 reads). We thus observed acceptable error rates for the vast majority of wells in both libraries.

**Fig. 4.**
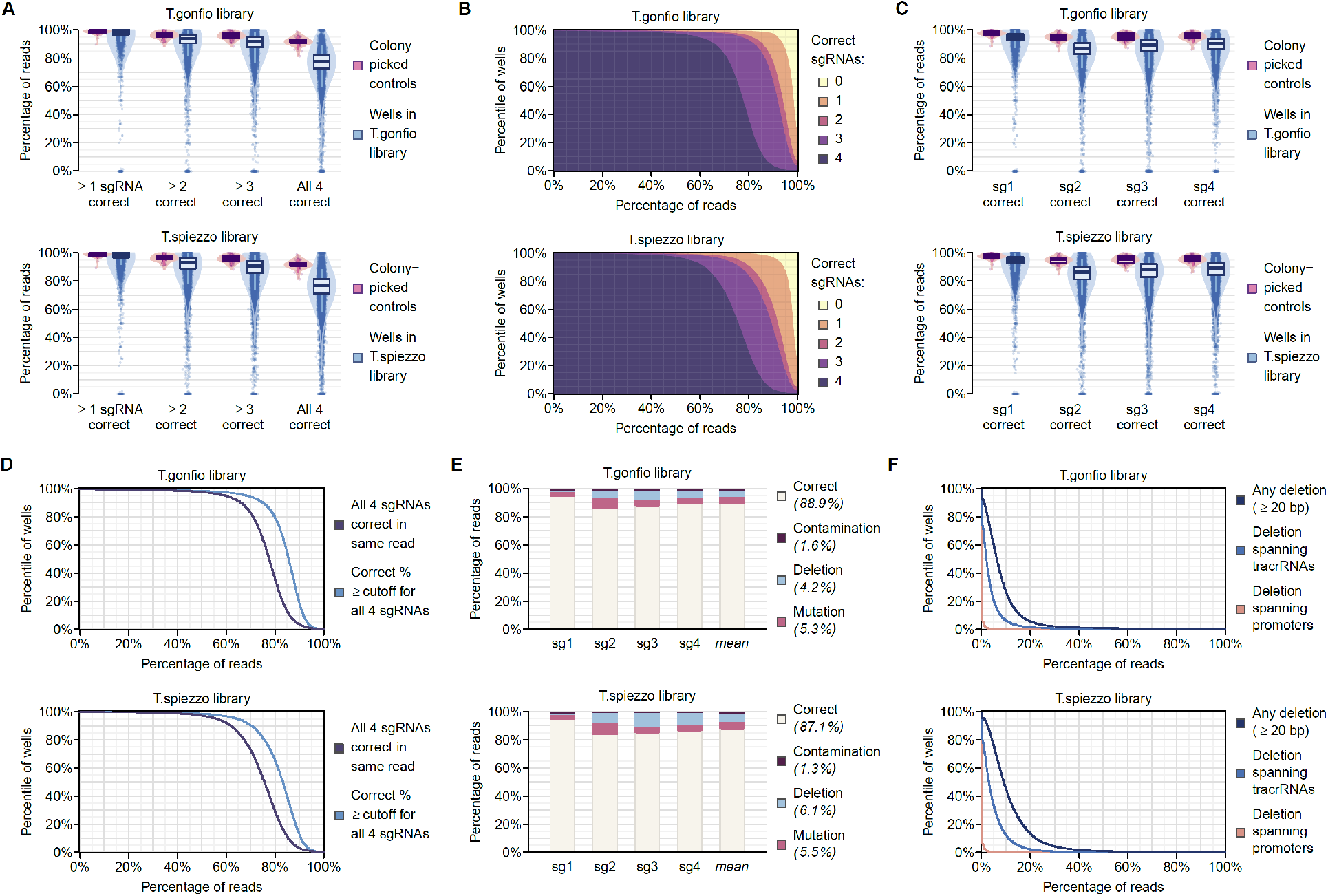
Genome-wide sequencing of the T.spiezzo and T.gonfio libraries. **A**, Reads with 0, 1, 2, 3, or 4 correct sgRNAs for each well in the T.spiezzo and T.gonfio libraries (quantitative SMRT long-read sequencing). To assess technical errors, we added barcoded amplicons of 74 single-colony-derived, sequence-validated 4sgRNA plasmids as internal NGS controls. The box plot denotes the median and interquartile range; the whiskers indicate the 5th and 95th percentile. **B**, Cumulative distribution of the percentage of reads with 0-4 correct sgRNAs in each well of the T.spiezzo and T.gonfio libraries. **C**, Percentage of correct sgRNA-1, sgRNA-2, sgRNA-3, and sgRNA-4 cassettes among the plasmid pools in each well of the T.spiezzo and T.gonfio libraries. **D**, Percentage of reads with four entirely correct sgRNAs in the same vector (black) and minimum percentage threshold passed by each of the four sgRNAs individually (blue) considering the entire pool of plasmids in each well. **E**, Mean percentage of mutations, deletions, and cross-well contaminations in the T.spiezzo and T.gonfio libraries. **F**, Cumulative distribution of plasmids with recombination in each well of the T.spiezzo and T.gonfio libraries.

When considering the four sgRNAs individually, the median percentage of correct reads was ≥85% for all four sgRNAs in both the T.gonfio and T.spiezzo libraries (**Fig. 4C**). While 65% of wells in the T.gonfio library had ≥75% reads with four entirely correct sgRNAs (within the same read), when each sgRNA is considered separately, 90% of wells were ≥75% correct for each of the four individual sgRNAs (**Fig. 4D**). Hence mutations may be compensated for by other clones in the same well. In a more stringent analysis where sgRNAs were considered correct only if the preceding promoter sequence was ≥95% correct, these percentages remained similar (**Fig. S4C**-**D**).

Incorrect sgRNAs were classified as contaminated (matching sgRNAs from other wells), deleted, or mutated (**Fig. 4E**). Contaminations were rare (1.6% and 1.3% contaminating sgRNAs in T.gonfio and T.spiezzo, respectively). Large deletions (≥50% of sgRNA and tracrRNA) affected 4.2% and 6.1% sgRNAs in T.gonfio and T.spiezzo, respectively. 4.1% of reads were affected by deletions spanning two tracrRNAs, whereas 0.1% of reads contained deletions spanning two promoters (**Fig. 4F**). The mean percentage of plasmids with deletions affecting ≥1 sgRNA was 8.1% in T.gonfio and 11.4% in T.spiezzo, whereas mutations affected 5.3% and 5.5% of sgRNAs in T.gonfio and T.spiezzo, respectively. These estimates include PCR- and sequencing-derived errors and may overestimate the error rate. Crucially, off-target activities were acquired in only ˂0.1% of mutated sgRNAs (**Fig. S4E- F**). Perfectly correct sequences were observed in 88.9% and 87.1% of sgRNAs for T.gonfio and T.spiezzo, respectively. We conclude that APPEAL cloning resulted in the generation of these libraries with low overall error rates.

### Benchmarking of 4sgRNA Ablation Plasmids in Cells and Organoids

Next, we sought to benchmark our library plasmids against available CRISPR reagents (lentiviral packaged sgRNAs and synthetic guide RNAs) in the immortalized human colon cancer cell line HCT116, in iPSCs, and in kidney organoids using several delivery methods (transduction, transfection, and electroporation) (**Fig. S5A**). We focused on gene ablation and chose Epithelial Cell Adhesion Molecule (*EPCAM*), cell-surface glycoprotein *CD44,* and phosphatidylinositol glycan anchor biosynthesis class A (*PIGA*) as targets, based on their detectability with live-cell immunostaining and flow cytometry quantification (**Fig. S5B**). First, we transduced Cas9 expressing HCT116 (HCT116- Cas9) or doxycycline-induced Cas9 expressing iPSC (iPSC-iCas9) cells with either the lentivirally packaged 4sgRNA vector or with a pool of four individually packaged sgRNAs (ThermoFisher) at a multiplicity of infection (MOI) of 5. Transduction of both reagents resulted in a significant reduction of EPCAM and CD44 expression in a time-dependent fashion. However, the T.spiezzo lentiviruses achieved higher ablation efficiency in both cell models at 4-8 days post-transduction (**Fig. S5C**). Next, we transfected HCT116-Cas9 cells (iPSCs were not transfected due to their poor transfectability) with our 4sgRNA plasmids or a pool of four individual synthetic sgRNAs (Integrated DNA Technologies). The synthetic sgRNAs showed more rapid ablation than the 4sgRNA plasmids at day 4 post-transfection, but both reagents resulted in a similar reduction of EPCAM and CD44 detection at day 8 (**Fig. S5D**). Next, we electroporated HCT116-Cas9 and iPSC-iCas9 cells with either the 4sgRNA vectors or a pool of four individual synthetic sgRNAs (Integrated DNA Technologies). In HCT116- Cas9 cells, both reagents worked, but the four synthetic sgRNAs approach resulted in a faster and more efficient reduction of EPCAM and CD44 than the 4sgRNA vectors (**Fig. S5D**). In contrast, in iPSC-iCas9 cells, electroporation of synthetic sgRNAs showed <50% knockout efficacy whereas the 4sgRNA vector resulted in fast and highly efficient editing (**Fig. S5D**).

To further test whether our 4sgRNA vector approach can efficiently edit target genes in complex cellular models, we used an inducible Cas9 iPSC line (Ungricht et al., 2022) to generate nephron progenitor cells and further differentiated them into kidney organoids following established protocols (Morizane and Bonventre, 2017). The *PIGA* gene, which is essential for the synthesis of glycosylphosphatidylinositol inositol (GPI) anchors, was targeted, and its editing efficiency was assessed by staining with a non-toxic fluorescently labelled aerolysin (FLAER assay) (Metzakopian et al., 2017). Progenitor cells were transduced with the lentivirally packaged 4sgRNA vector or with a pool of four individually packaged sgRNAs (ThermoFisher) at increasing volumes of viral supernatant, and after 48 days, the organoids were dissociated into single cells and subsequently stained with FLAER. Lentiviruses carrying the 4sgRNA vector already showed a high knockout efficiency at low lentiviral volumes, whereas four individually packaged sgRNAs required a higher volume to achieve a similar knockout efficiency, even at equal viral titers (**Fig. S5C**). These results indicate equal or superior gene perturbation with T.spiezzo than with existing CRISPR reagents in notoriously fastidious cell models.

### An Arrayed CRISPR Activation Screen for Transcription Factors Controlling PrP^C^ Expression

The cellular prion protein PrP^C^, encoded by the *PRNP* gene, is essential for the development of prion diseases (Bueler et al., 1993; Scheckel and Aguzzi, 2018). Previous genome-wide microRNA and siRNA screens have uncovered a complex pattern of regulated expression of PrP^C^ (Heinzer et al., 2021; Pease et al., 2019). However, all previously published screens failed to identify any transcription factors (TFs) regulating PrP^C^ expression. We therefore measured PrP^C^ expression in a focused arrayed activation screen by time-resolved fluorescence resonance energy transfer (TR-FRET) using a pair of antibodies binding distinct domains of PrP^C^ (**Fig. 5A**).

**Fig. 5.**
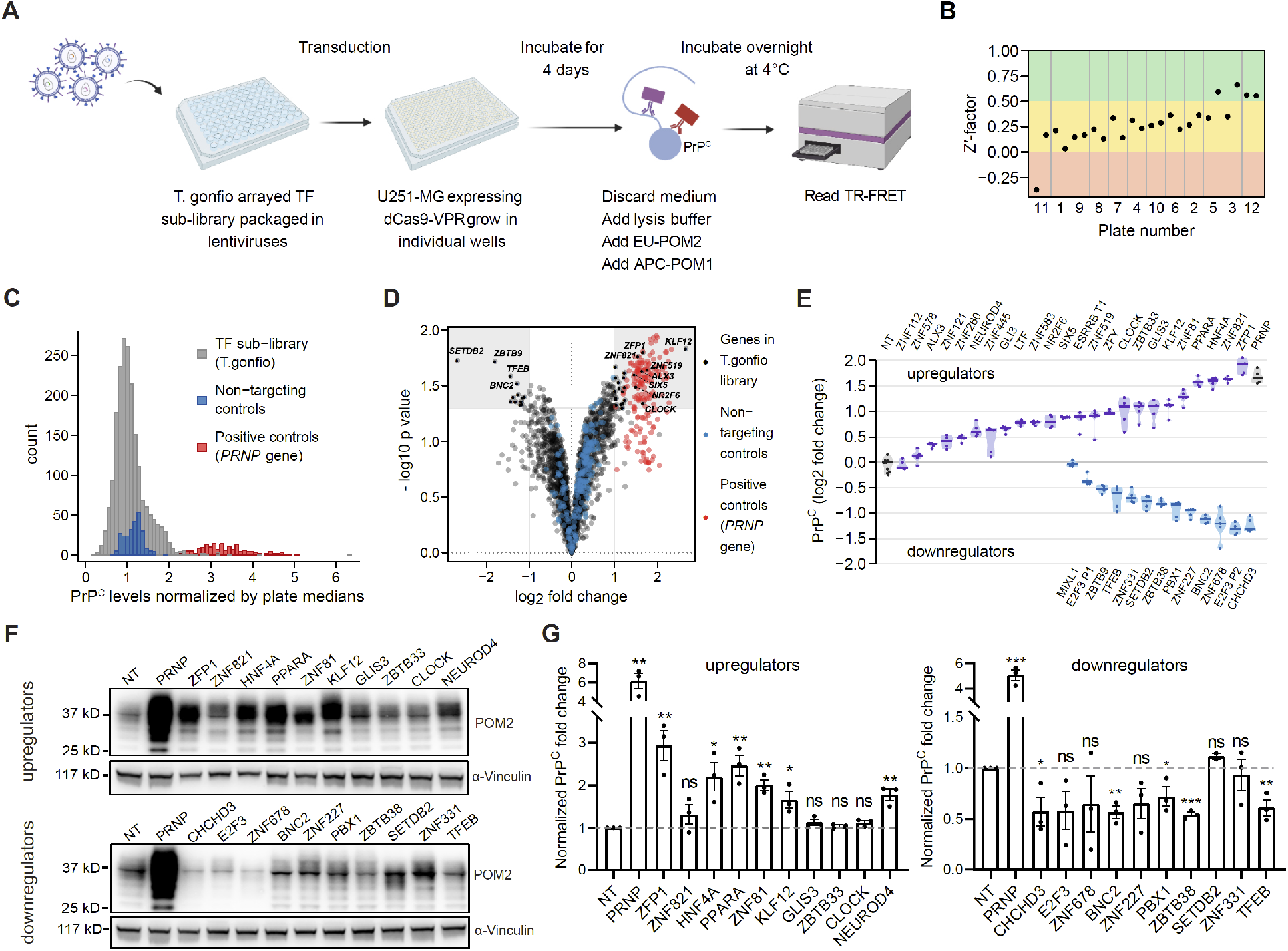
Transcription factors (TFs) regulating PrPC expression identified by arrayed activation screen with T.gonfio. **A**, Schematic of the primary PrPC TF sublibrary screen. **B**, Z’ factor of each plate from the primary screen. **C**, Distribution of positive controls (4sgRNA targeting *PRNP*), negative controls (4sgRNA non-targeting control) and samples. **D**, Volcano plot displaying –log10 p values and log2 fold change values across the T.gonfio TF sublibrary. The 36 candidate genes and *PRNP* are depicted in black and red, respectively. **E**, Repetition of the 36 candidate genes in their regulation of PrPC levels (as in **A**); NT, nontargeting control. P1 or P2 indicates the 4sgRNA plasmid targeting TSS1 or TSS2 transcriptional start site of a defined gene, respectively. **F**, An example of western blotting analysis of PrPC expression after individually activation of the top 10 most pronounced candidate genes from the upregulators or downregulators confirmed in **E**; POM2, primary antibody against PrPC, α-Vinculin, primary antibody against Vinculin (internal control). **G**, Quantification of PrPC levels after individually activation of the top 10 most pronounced candidate genes from the upregulators or downregulators as described in **F**. Dots: independent experiments (N = 3 biological repeats, Mean ± S.E.M.). p values were determined by Student’s t-test.

A T.gonfio sublibrary encompassing all human TFs (n= 1,634) was packaged into lentiviral vectors and transduced into U-251MG human glioblastoma cells stably expressing the CRISPR activator dCas9-VPR (MOI = 3). Experiments were performed as triplicates in 36 plates of 384-well microplate, each including 14 wells with non-targeting (NT) and 14 *PRNP* targeting controls. Cells were lysed 4 days post-transduction; one replicate plate was used to determine cell viability, and two replicates were used to assess PrP^C^ levels (**Fig. 5A**). Nineteen and four plates had a Z′ factor of 0–0.5 and >0.5, respectively, whereas one plate had a Z′<0 (Zhang, 2011) (**Fig. 5B**). Distinct levels of PrP^C^ were detected between non-targeting and *PRNP-*targeting controls (**Fig. 5C**). The Pearson correlation coefficient (R^2^) between duplicates was 0.77 (**Fig. S5E**). Hit calling was based on an absolute log_2_ fold change of ≥1 and a p-value of ≤ 0.05. Together, 24 and 12 genes were found to upregulate or downregulate PrP^C^ expression, respectively (**Fig. 5D**).

We repeated the TR-FRET assay with these 36 candidate hits, and found that 19 up- and 10 downregulators (log2 fold change ≥ 0.5) could be reproduced (**Fig. 5E**). We then validated the most pronounced 10 up- and 10 down-regulators by an orthogonal method (via Western blotting for PrP^C^), and confirmed their effect in six and five instances, respectively (**Fig. 5F** and **5G**).

Together, these results confirm the feasibility and power of our libraries for studying phenotypes of interest in an arrayed format.

### Novel Modifiers of Autophagy Uniquely Identified by the Pooled T.spiezzo Library

4sgRNA plasmids outperformed all single sgRNAs from which they were assembled, suggesting that they could be used efficaciously in a pooled format. Importantly, the complexity of T.gonfio and T.spiezzo is much lower than that of pooled libraries, which usually contain 5-10 different sgRNA vectors targeting the same gene, which may result in leaner, more sensitive and less expensive screens.

We pooled all 19,820 T.spiezzo 4sgRNA plasmids and performed a genome-wide CRISPR screen aimed at identifying modulators of autophagy by measuring the autophagy marker SQSTM1 protein (DeJesus et al., 2016; Larsen et al., 2010). For comparison, we repeated the same screen with the Brunello (Sanson et al., 2018) and Cellecta (DeJesus et al., 2016) libraries (**Fig. 6A**). Each library was packaged into lentiviruses and transduced (MOI = 0.4) into H4 cells stably expressing Cas9 and a green fluorescent protein (GFP)-tagged SQSTM1. Cells were selected with puromycin for 3 days and maintained in non-selective medium for 7 days. Then, GFP^high^ or GFP^low^ (upper or lower quartile of GFP fluorescence, respectively) cells were separated and collected by fluorescence activated cell sorting (FACS) (**Fig. 6A** and **S6A**). Genomic DNA was isolated and the abundance of sgRNAs was determined by Illumina paired-end next-generation sequencing (NGS).

**Fig. 6.**
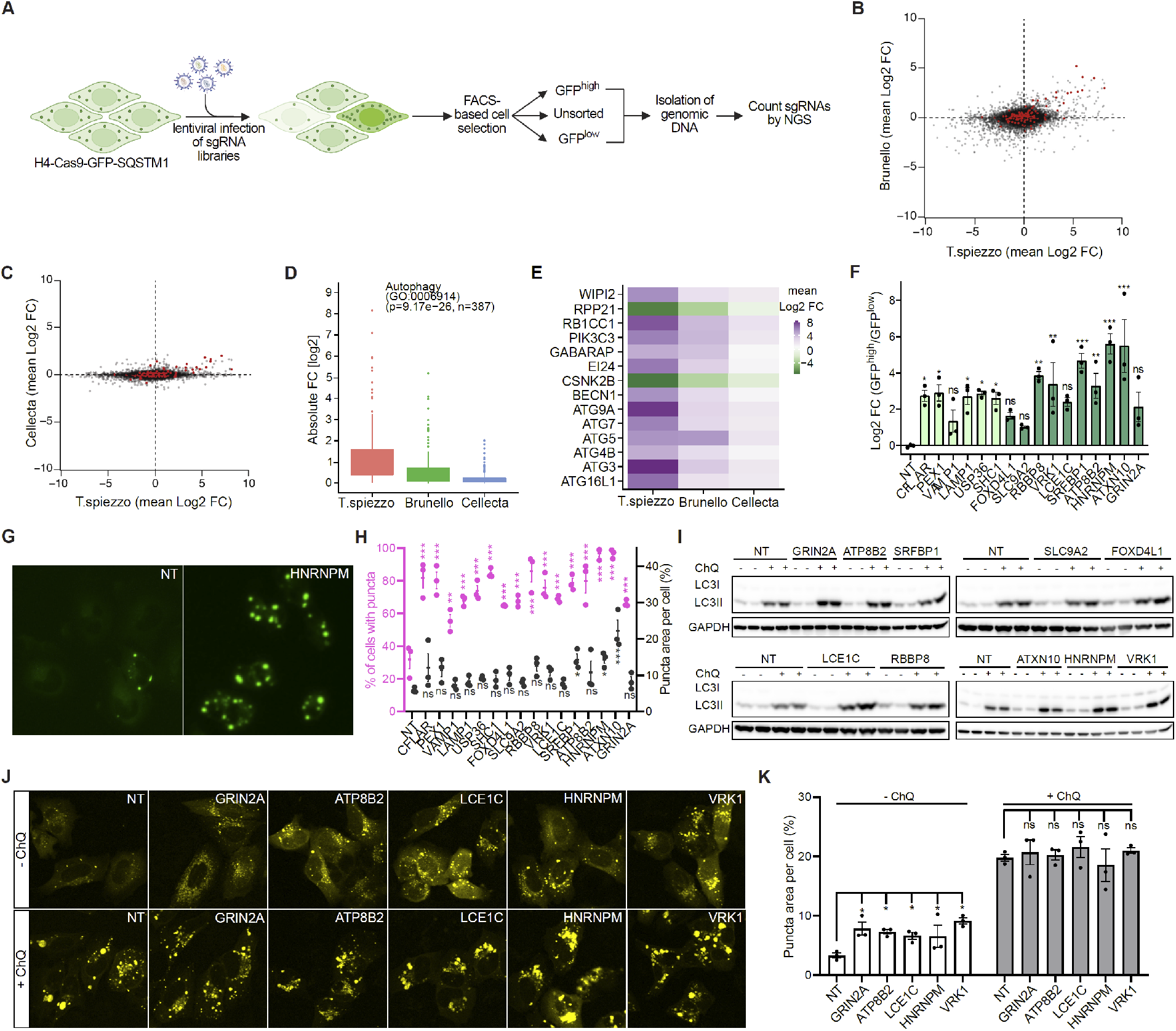
Novel autophagy modifiers identified by pooled T.spiezzo screen. **A,** H4-Cas9-GFP-SQSTM1 cells were transduced with genome-wide pooled lentiviral sgRNA ablation libraries (T.spiezzo, Brunello, and Cellecta) and selected with puromycin. Cells in the uppermost and lowermost fluorescence quartile were collected by FACS and sgRNAs were quantified by sequencing genomic DNA. **B** and **C,** Average log2 fold change (FC) in GFP^high^ and GFP^low^ samples from T.spiezzo versus Brunello **(B)** and T.spiezzo versus Cellecta **(C)**. Autophagy-relevant genes (autophagy, GO:0006914) are highlighted in red. **D,** Autophagy genes enriched in GFP^high^ cells from the T.spiezzo, Brunello and Cellecta screens. **E**, Heatmap showing the highest and lowest mean log2 FC (GFP^high^ versus GFP^low^) of genes identified among the top 100 genes in in the T.spiezzo, Brunello and Cellecta screens. **F**, Quantification of log2 FC of cell count in GFP^high^ vs GFP^low^ populations transduced with T.spiezzo 4sgRNA lentivirus against nontargeting (NT) control or each of the 16 genes selected for validation. The boundaries for GFP^high^ and GFP^low^ cell populations were set according to the NT condition. Dots (here and henceforth): independent experiments, Mean ± S.E.M.. **G**, An example of GFP-SQSTM1 puncta in H4-Cas9-GFP-SQSTM1 cells transduced with T.spiezzo 4sgRNA lentivirus against NT or HNRNPM (see Fig. S6O for all genes tested). **H**, Percentage of cells with GFP-SQSTM1 puncta (purple) and puncta area (black) in H4-Cas9-GFP-SQSTM1 cells transduced with T.spiezzo 4sgRNA lentivirus against 16 genes selected for validation. **I**, Immunoblotting of LC3-II from H4-Cas9 cells transduced with T.spiezzo 4sgRNA lentivirus against 10 possible autophagy modulators in the absence (-) or presence (+) of chloroquine (ChQ, 100 µM, 6 hrs). GAPDH: loading control. Two biological repeats were assessed for each condition. **J**, Representative micrographs of YFP-LC3 in H4-Cas9-cells transduced with T.spiezzo 4sgRNA lentivirus against NT or each of the 5 genes selected for further validation. H4-Cas9 cells were cultured and treated as described in **A** and transduced with YFP-LC3 lentiviruses 48– 60 hours before examining YFP-LC3 puncta. **K**, Quantification of puncta area of YFP-LC3 of cells and conditions described in **J**. N = 3 biological repeats, mean ± S.E.M.. p values were determined by Dunnett’s test.

When packaging pooled libraries into lentiviral particles, reverse transcriptase-mediated template switching may generate chimeric products with unpredictable sequences (Hill et al., 2018; Jetzt et al., 2000; Nikolaitchik et al., 2013). By sequencing sgRNA2 and sgRNA3 sequences from individual 4sgRNA plasmids in pools of transduced cells, we found that the percentage of intersected reads (correct alignment and linkage of sgRNA2 and sgRNA3) from the T.spiezzo pool was around 70% (**Fig. S6B** and **Table S2-4**), indicative of moderate lentiviral template switching. For all further analysis with pooled libraries, we only used reads that aligned correctly and where both sgRNAs mapped to the same plasmid.

By comparing sgRNA representational differences between GFP^high^ and GFP^low^ samples treated with T.spiezzo, Brunello or Cellecta, we identified many well-characterized modulators of autophagy including *ATG5*, *ATG7*, *BECN1*, *WIPI2* and *PIK3C3* (red dots in **Fig. 6B**-**C, Table S2-4**). Gene ontology (GO) enrichment analysis of the 200 most enriched genes in GFP^high^ samples showed that T.spiezzo identified more autophagy-related pathways than the other libraries (**Fig. S6C-E**). Importantly, the enrichment of validated autophagy modulators was significantly higher for T.spiezzo (**Fig. 6D** and **Fig. S6F-M**). To directly compare representational changes between all three screens, we selected the 100 genes with the highest and lowest fold change for each screen and intersected the lists to produce a set of shared genes. The T.spiezzo library identified all shared genes with a higher fold change compared to Brunello and Cellecta (**Fig. 6E**).

In addition, T.spiezzo (but neither Brunello nor Cellecta) identified certain genes that had not been highlighted as autophagy-relevant by GO analysis (grey dots in Fig. **6B**-**C**). To validate these potential novel modifiers of autophagy, we focused on genes with log2 fold changes ≥ 5 in the T.spiezzo screen and ≤1 in the Brunello and Cellecta screens (**Table S5**). Among the top list, we found genes (e.g. *CFLAR, PEX1, VAMP1, LAMP1, USP36* and *SHC1*) that had been reported to modulate autophagy (Cheng et al., 2018; Geisler et al., 2019; He and He, 2013; Judy et al., 2022; Mao et al., 2019; Taillebourg et al., 2012; Zheng et al., 2013), supporting the idea that some of these genes may be genuine autophagy regulators.

We then attempted to validate the regulation of autophagy by ten hit genes (*FOXD4L1, SLC9A2, RBBP8, VRK1, LCE1C, SRFBP1, ATP8B2, HNRNPM, ATXN10,* and *GRIN2A*) with no reported association with autophagy, and the six previously reported genes mentioned above. We first repeated the measurement of GFP-SQSTM1 signal intensity in GFP^high^ and GFP^low^ cell populations after individually ablating each of these 16 genes. We observed a prominent shift of GFP-SQSTM1 intensity towards GFP^high^ for all 16 genes tested (**Fig. 6F** and **S6N**). In parallel, we assessed the cytosolic distribution of the puncta of GFP-SQSTM1 by microscopy. Ablation of each gene significantly increased the percentage of cells exhibiting puncta of GFP-SQSTM1 over that of nontargeting controls, and ablation of SRFBP1, HNRNPM or ATXN10 further enlarged the size of the puncta (**Fig. 6G**-**H** and **S6O**).

The convergence between the primary screen and the validation experiments encouraged us to confirm the potential novel autophagy regulators with orthogonal methods. We thus analyzed the turnover of a second autophagy marker, LC3-II, by western blotting in the presence or absence of 100 µM chloroquine (ChQ), which blocks the autophagy flux by preventing autolysosome maturation (Mauthe et al., 2018). Compared to vehicle treatment, ChQ (6 hours) treatment conspicuously increased the levels of LC3-II in all samples. We found that ablation of *GRIN2A, ATP8B2, LCE1C, HNRNPM,* or *VRK1* further increased LC3-II levels in ChQ-treated cells consistently compared to ChQ-treated non-targeting control cells, whereas the other genes showed no or inconsistent effects (**Fig. 6I**). We then ablated these 5 genes individually and examined their effect on the level and distribution of the YFP-LC3 autophagy reporter (Flavin et al., 2017). All five genes, when ablated, significantly increased the puncta area of YFP-LC3 compared to nontargeting controls (**Fig. 6J**) but did not further modulate the area of YFP-LC3 puncta after ChQ treatment (**Fig. 6K**). We infer that the pooled T.spiezzo screen identified *GRIN2A*, *ATP8B2*, *LCE1C*, *HNRNPM*, and *VRK1* as novel bona fide modulators of the autophagic flux.

### T.gonfio Library for Targeted Epigenetic Silencing (CRISPRoff)

CRISPR-mediated targeted epigenetic silencing (Amabile et al., 2016; Nunez et al., 2021) is a loss- of-function perturbation method used as an alternative to knockout for interrogating gene functions, especially for cell models that are sensitive to DNA breakage such as iPSCs (Ihry et al., 2018). The CRISPRoff epigenome memory editor is robust and efficient for targeted gene silencing which persists even in iPSCs-differentiated neurons (Nunez et al., 2021). The sgRNA targeting window of CRISPRoff relative to the transcription start site (TSS) is quite broad and we found that 96.8% of T.gonfio sgRNAs coincided with it, with the remaining 3.2% of sgRNAs falling within <100 bp of it (**Fig. 7A**). This encouraged us to examine the effectiveness of using T.gonfio plasmids for CRIS- PRoff.

**Fig. 7.**
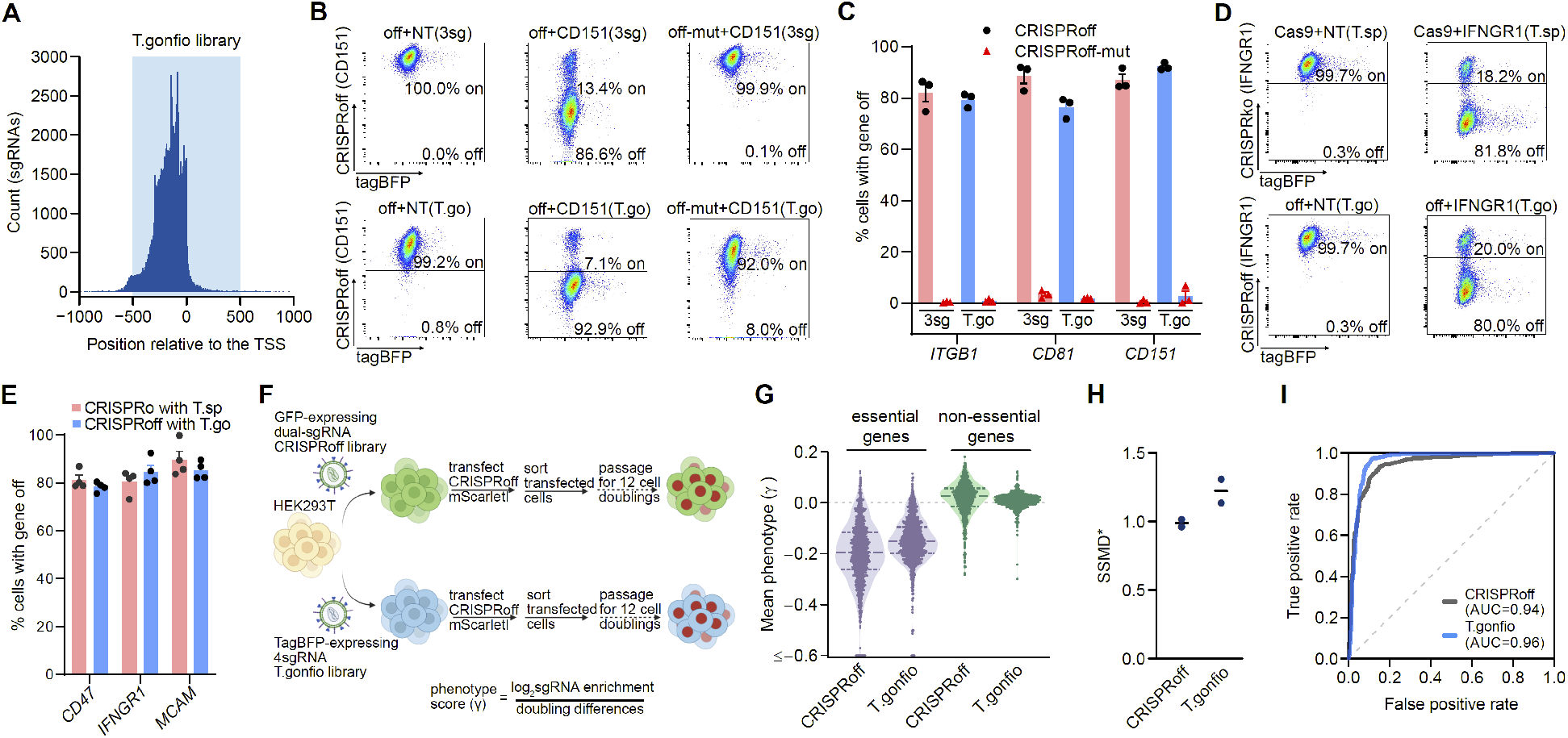
Targeted epigenetic silencing (CRISPRoff) with the T.gonfio library. **A**, Alignment of the T.gonfio sgRNAs sequences to the CRISPRoff targeting window. **B**, An example of flow cytometry measurement of CD151 in HEK293T cells after exposure (10 days) to pools of three separated sgRNAs (3-sgRNA, used in the CRISPRoff study (Nunez et al., 2021)) or the respective T.gonfio 4sgRNA. Off represented the CRISPRoff plasmid (Addgene #167981); off-mut represented the CRISPRoff mutant carrying a catalytically inactive version of the DNA methyltransferase. **C**, Quantification of percentage of cells with ITGB1, CD81 and CD151 silencing 10 days post CRISPRoff or CRISPRoff mutant with pools of three single sgRNAs (3-sg) or 4sgRNA from the T.gonfio library in HEK293T cells. Each dot represents an independent biological repeat of the assay. Data were presented in Mean ± S.E.M.. **D**, An example of flow cytometry measurement of percentage of cells with IFNGR1 silencing 10 days post CRISPR knockout with 4sgRNA plasmid from the T.spiezzo library or CRISPRoff with 4sgRNA from the T.gonfio library in HEK293T cells. **E**, Quantification of percentage of cells with CD47, IFNGR1 and MCAM silencing 10 days post CRISPR knockout with 4sgRNA plasmids from the T.spiezzo library or CRISPRoff with 4sgRNA plasmids from the T.gonfio library in HEK293T cells. Each dot represents a biological repeat of the assay. Data were presented in Mean ± S.E.M.. **F**, A schematic of pairwise pooled genome-wide CRISPRoff screens to determine the effectiveness of T.gonfio for gene silencing at the genome scale in HEK293T cells. **G**, A violin plot of the phenotype scores (γ) for essential and non-essential genes investigated in (Nunez et al., 2021). N=2 biological repeats. **H**, SSMD value measuring the separation between essential and non-essential genes for each repeat of the screens performed with the CRISPRoff or T.gonfio pool libraries. **I**, Plots of true and false positive rates of essential genes from screens with the CRISPRoff or the T.gonfio library. AUC, area under the curve.

4sgRNA plasmids targeting the TSSs of the cell-surface proteins ITGB1, CD81 and CD1 were cotransfected into HEK293T cells along with the plasmid for CRISPRoff or a CRISPRoff mutant encoding a catalytically inactive version of the DNA methyltransferase (Nunez et al., 2021). A pool of three single sgRNA plasmids used in the CRISPRoff study (Nunez et al., 2021) was used as reference for each gene. The 4sgRNA plasmids achieved a comparable gene silencing efficiency as the pool of three CRISPRoff individual sgRNA plasmids for all target genes, whereas the mutant dCas9- DNA methyltransferase complex (CRISPRoff-mut) induced minimal or no reduction of gene expression (**Fig. 7B**-**C**). We then tested three additional cell-surface proteins, CD47, IFNGR1 and MCAM, in the knockout efficiency assay (see **Fig. 2D**), to further confirm the usefulness of T.gonfio for CRIS- PRoff. In addition, we were curious to compare the gene silencing efficacy of CRISPRoff with T.gonfio plasmids with that induced by gene knockout via the corresponding T.spiezzo plasmids. The T.spiezzo plasmids with Cas9 induced 80–90% gene ablation, and the T.gonfio plasmids with CRIS- PRoff induced a similar extent of gene silencing (**Fig. 7D**-**E**).

The successful targeted gene silencing of the six cell-surface protein-coding genes tested above encouraged us to examine the generalized suitability of T.gonfio-mediated epigenetic silencing. We adapted the same genome-wide dropout screen from the CRISPRoff study (Nunez et al., 2021) (see also **Fig. 7F**) and benchmarked T.gonfio against the refined pooled dual-sgRNA CRISPRoff library, which incorporated a more expansive set of screening data to prioritize guides (Nunez et al., 2021; Replogle et al., 2022). The TagBFP expressing 4sgRNA T.gonfio (pool of 22,442 plasmids) and the dual-sgRNA CRISPRoff (GFP-expressing) libraries were packaged into lentiviral particles and transduced to HEK293T cells (MOI = 0.3-0.4). After puromycin selection, cells were transiently transfected with a CRISPRoff plasmid concomitantly expressing the red fluorescent protein mScarletI (CRIS- PRoff-mScarletI). Two days later, cells double-positive for the sgRNA and CRISPRoff-mScarletI plasmids were sorted and maintained without antibiotic selection for ≥10 cell divisions, allowing sgRNAs silencing essential genes to drop out of the population. Guide abundance was compared between cell populations at baseline (immediately before the transfection of CRISPRoff-mScarletI) and at end point (after sorting and passaging) (**Fig. 7F**). The phenotype score γ (defined as log_2_(sgRNA enrichment)/the number of cell doublings) of each gene was calculated (**Fig. 7F**).

The dual-sgRNA CRISPRoff library induced a similar growth defect for the set of essential genes as in the original CRISPRoff study (Nunez et al., 2021) (see also **Fig. 7G** and **Fig. S7A**). We saw a similar fitness effect with the pooled T.gonfio library, but the CRISPRoff library showed a wider spread of phenotype scores for essential genes, whereas T.gonfio produced a more homogeneous distribution, with most scores for these genes lying close to the median. Similar differences in the variance were seen for non-essential control genes (**Fig. 7G** and **Fig. S7A**). Accordingly, we found a higher SSMD (strictly standardized mean difference) score for the T.gonfio library than the CRISPRoff library (**Fig. 7H**), indicating a stronger separation between essential and non-essential genes. Furthermore, we saw slightly greater accuracy and a lower false-positive rate for essential genes in the screen with the T.gonfio library (**Fig. 7I**). We conclude that the T.gonfio library can be adopted for efficient epigenetic silencing of individual genes in an arrayed format or at the genome scale in pooled format when combined with an appropriate dCas9 variant.

## Discussion

The arrayed libraries described here are meant to enhance the armamentarium necessary for the study of complex and non-autonomous phenotypes. We have addressed issues limiting the performance of CRISPR screens, such as the variable targeting efficacy of single sgRNAs and the impact of human genomic variability. The sgRNA selection algorithm was adapted to ensure that the sgRNAs were non-overlapping and would tolerate to the largest possible extent DNA polymorphisms found among 77,781 human genomes with minimal off-target effects.

We found that arrayed CRISPR screens can be less laborious than one might presume, and can be performed rapidly by standardizing workflows and by deploying inexpensive automation steps such as 384-well pipetting.

Covering the entire protein-coding genome with ablating, activating and silencing tools required the generation of >42,000 individual plasmids. This would have required prohibitive resources, since agar-plating, colony-picking and gel extraction of desired DNA fragments (McCarty et al., 2020) are time-consuming and refractory to automation. The APPEAL plasmid construction method, which features an additional antibiotic resistance element within the clonable insert, radically simplifies the cloning procedure and renders it massively parallel. Importantly, APPEAL is generically applicable to any kind of molecular cloning task. When combined with adjustments aimed at minimizing the likelihood of illegitimate recombination events, such as the insertion of unique promoter and tracrRNA sequences for each of the four sgRNA cassettes, APPEAL allowed generation of ∼2,000 individual 4sgRNA plasmids per week with high accuracy by two full time employees.

### Efficacy of CRISPR-Based Gene Perturbation

Although the algorithms for predicting sgRNA activity are continuously improving (Hanna and Doench, 2020), the efficacy of gene-perturbation screens can still be jeopardized by suboptimal guide efficiency, leading to false-negative hit calls. Pooled libraries can partially circumvent this by increasing the number of sgRNAs targeting the same gene. The strategy is cost-inefficient for arrayed genome-wide libraries. Thus, augmenting the efficiency and robustness of gene perturbation is key for high-performance arrayed genome-scale libraries. The integration of four sgRNAs into each targeting vector dramatically improves CRISPR-mediated gene activation, ablation, and silencing.

### Versatility of 4sgRNA-Based Libraries

To ensure the broadest possible adaptability to multiple experimental model systems (immortal cell lines, hiPSCs, organoids, or primary cells), we included two selection markers in our 4sgRNA vector (puromycin resistance and TagBFP) and the motifs necessary for lentiviral packaging and transposon-mediated integration. Furthermore, each sgRNA is driven by a different housekeeping promoter, ensuring activity in the broadest range of cells and tissues and minimizing the risk of transcriptional silencing by promoter methylation. The sgRNA selection algorithm was tuned to identify the least polymorphic regions of each gene, thereby extending the likelihood of perturbation to patient-derived cells that may substantially differ from the human reference genome. Consequently, our libraries proved useful for editing organoids.

Somewhat unexpectedly, we found that T.gonfio library can be efficiently used for epigenetic silencing (CRISPRoff). This opens the possibility of performing both loss-of-function and gain-of-function screens using the same library in cell lines expressing the appropriate dCas9 proteins, further saving time, cost and labor in the execution of gene-perturbation screens. Genome-wide pairwise screens showed that T.spiezzo and T.gonfio provided increased signal/noise ratios than the existing pooled libraries, Brunello (Sanson et al., 2018) and CRISPRoff (Nunez et al., 2021), respectively. What’s more, our pooled libraries have a much smaller size compared to existing libraries that require up to 10 guides per gene. This can not only reduce workload and cost, but enables screens when cell numbers are limiting – which is a frequent problem with human primary cells.

### Quality and Homogeneity of the 4sgRNA Vectors

By dramatically lowering the percentage of incorrect plasmid assemblies, APPEAL eliminates the necessity of isolating clonal bacterial colonies. Consequently, each APPEAL reaction product generally represents a polyclonal pool of plasmids. This source of variability was quantitatively assessed by sequencing: in the average well, 90% of the plasmid population contained ≥3 intact sgRNAs. Some 4sgRNA plasmids showed mutations in the region of the sgRNA and tracrRNA sequences, possibly originating from oligonucleotide synthesis and Taq DNA ligase reactions. The mutation rate found in our sgRNA vector sequences is consistent with the expected error rate occurring during Gibson assembly, leading to mutations in approximately 10% of the plasmids (Gibson et al., 2009). Despite the use of four different promoters and tracrRNA variants, we observed a recombination between sgRNA expression cassettes in ∼10% of reads, resulting in a deletion of the intervening sequence. However, 85% of recombining plasmids retained ≥1 correct sgRNA sequence, and in the median well, 99.7% of reads had at least one sgRNA+tracrRNA module that was 100% correct. Accordingly, the plasmid collections remained highly functional even in wells affected by recombination events. The average percentage of entirely correct protospacer and tracrRNA sequences for each of the four sgRNAs was ∼90%.

Mutated sgRNAs may target genomic sites illegitimately and induce off-target effects. The latter occurrence affected <0.5% of mutated sgRNAs, and targeted additional genes in only ∼0.01% of cases (**Fig. S4E**-**F**). Hence the sequence alterations in APPEAL-generated plasmid pools had no practical effect apart from a slight reduction in the number of active sgRNAs. Notably, the 74 single colony-derived and Sanger sequencing-confirmed control plasmids displayed several errors attributable to PCR or sequencing steps, suggesting that the error rates reported for T.gonfio and T.spiezzo are likely overestimates.

### An Arrayed Assessment of Transcription Factors Regulating PrP^C^ Expression

The cellular prion protein PrP^C^ encoded by the *PRNP* gene is required for the development and pathogenesis of prion disease as evidenced by studies that mice devoid of PrP^C^ coding gene *PRNP* is resistant to prion infection (Bueler et al., 1993) and neural tissues lacking PrP^C^ are resistant to scrapie-induced toxicity (Brandner et al., 1996). Thus, genes or drugs modifying the level of PrP^C^ expression are of general interest for both the basic biological understanding and therapeutic treatment of prion diseases. We have previously identified a wealth of modulators of PrP^C^ expression using arrayed genome-wide screens with microRNAs (Pease et al., 2019) and siRNAs (Heinzer et al., 2021). However, these screens did not identify any transcription factors regulating PrP^C^ expression, and the transcriptional regulation of *PRNP* has remained largely unexplored. Our arrayed CRISPR activation screen focusing on the TF sublibrary has thus partially filled the gap by successfully identifying 11 TFs regulating PrP^C^ expression, and these TFs provide an opportunity for further study to close the gap.

### A Pooled Screen for Genetic Modifiers of Autophagy

Autophagy is a highly-regulated biological flux that include the formation of autophagosomes, autophagosomal engulfment of cytoplasmic components and organelles, fusion of autophagosomes with lysosomes to produce autolysosomes, and the degradation of intra-autophagosomal components by lysosomal hydrolases (Tanida et al., 2008). Approx. 200 genes have been identified as core components of autophagic flux (Bordi et al., 2021).

Dysfunction of autophagy has been implied in diseases including neurodegenerative disease, diabetes, tumors, and immune diseases (Bordi et al., 2021). The accumulation of GFP-tagged SQSTM1 provides a reliable proxy of autophagic activity, motivating us to compare the sensitivity and specificity of a pooled T.spiezzo version with the two pooled CRISPRko libraries, Brunello and Cellecta. Not only did T.spiezzo exhibit a much higher signal/noise ratio than the Brunello and Cellecta libraries, but it also identified several novel autophagy modifiers that were not uncovered by the other libraries, 5 of which were subsequently validated using orthogonal methods. Beyond contributing to our understanding of autophagy and providing a wealth of new potential therapeutic targets, these results add to the evidence that the T.spiezzo and T.gonfio libraries represent powerful tools for genome-wide interrogations in both arrayed or pooled modalities.

## Supporting information

T. gonfio library sgRNA information

T. spiezzo library sgRNA information

## Contributorship

J-A.Y. designed, supervised, and coordinated the research, invented the APPEAL cloning method, performed validation experiments of APPEAL with the assistance of A.S. and K.M., developed the custom-made Gibson assembly mix for the two libraries with assistance of A.S., produced (∼30%) custom-made competent cells with Y.W. (∼30%), L.Y. (∼30%), K.G. (∼8%) and A.S. (∼2%), transformed the Gibson assembly products (100%) of the two libraries into competent cells together with Y.W. (∼50%) and L.Y. (∼50%), stored (100%) bacterial glycerol stock of the two libraries into 384- well deep-well plate, developed the 96-well plate deep-well magnetic-beads based plasmid miniprep together with Y.W. and L.Y., setup the lentiviral production of the 4gRNA plasmids together with K.M., performed the 1sgRNA vs 4sgRNA gene activation real-time quantitative PCR with A.S. and K.M., performed 1sgRNA vs 4sgRNA gene knockout efficiency on CD47, IFNGR1 and MCAM together with J.G., analysed the 4sgRNA knockout efficiency obtained from SMRT long-read sequencing, performed transfection and transduction of non-transfectable cells with 4sgRNA vector together with K.M. and M.L., contributed to L.F. for the design of new algorithms for 4sgRNA selection, set up the barcoded 384-well plate library plasmid sequencing with A.S., performed CRISPRoff test experiments on ITGB1, CD81, CD151, CD47, IFNGR1, MCAM together with J.G, analysed data, performed the hits validation experiment of autophagy with F.B. and Andrea A., helped T.L. for the genome-wide CRISPRoff pooled screen, and wrote most of the manuscript. L.F. developed the 4sgRNA selection algorithm and analysed the in-silico features of the libraries, analysed the SMRT long-read sequencing of the two libraries, analysed the arrayed screen data on PrP^C^, aligned the sgRNA targeting window of T. gonfio library to the targeting window of sgRNAs for efficient targeted epigenetic silencing, performed the data analyses of CRISPRoff screens, wrote part of the manuscript. M.S. performed the benchmarking experiment on 4sgRNA knockout efficiency with commercially available resources (synthetic sgRNA and lentiviruses) in HCT116, iPSCs and kidney organoids with various vector delivery methods including transfection, transduction and electroporation, analysed the data, and performed the genome-wide autophagy screens with the assistance of B.K., wrote part of the manuscript. T.L. prepared samples for 4sgRNA knockout assay with SMRT longread sequencing of the 9 genes with assistance of K.M., performed the genome-wide CRISPRoff screens. C.T. performed the TF sublibrary arrayed screen TR-FRET experiment, validated the hits with help of J-A.Y. and T.L., and wrote a first draft of the respective section of the manuscript. A.D. performed cell culture and transduction of 4sgRNA lentiviruses into iNeurons, packaged the T.gonfio TF sublibrary plasmids into lentiviruses together with J.T. and S.R. A.S. performed maxiprep, *Bbs*I digestion of the pYJA5 vector, and purification of the digested pYJA5, performed three fragment PCRs of the entire two libraries, Gibson assembly of the three PCR amplicons with digested pYJA5 of the entire two libraries, tested Taq DNA ligase together with J-A.Y., and the above-mentioned experiments together with J-A.Y. Y.W. produced (∼30%) custo,made competent cells, transformed the Gibson assembly products (∼50%) of the two libraries into competent cells, performed ∼50% of bacterial glycerol stock of the two libraries into 96-well deep-well plates, and miniprepped ∼50% of plasmids of the two libraries. L.Y. produced (∼30%) custom-made competent cells, transformed the Gibson assembly products (∼50%) of the two libraries into competent cells, performed ∼50% of bacterial glycerol stock of the two libraries into 96-well deep-well plates, and miniprepped ∼50% of plasmids of the two libraries. D.L.V. performed barcoded PCR of the entire two libraries and pooled down each plate of the PCR products into a single tube and purified the PCR products for SMRT long-read sequencing. B.K. executed the pooled autophagy screens. E.D.C produced Taq DNA ligase used for Gibson assembly reaction. K.G. performed real-time quantitative PCR of cDNAs from iNeurons, assisted sometimes A.S. for three fragments PCR and Gibson assembly, and the above-mentioned experiments. Andrea A. helped J-A.Y. the design of experiment for autophagy hits validation, and performed the LC3-II western blot and corresponding analyses. E.O., J.J., and J.H. performed sequencing library preparations of the autophagy screens and sequencing. F.N. and M.P. performed data analysis of autophagy pooled screens. J.T. and S.R. helped A.D. for lentiviral production. J.G. performed 1sgRNA vs 4sgRNA knockout efficiency and CRISPRoff assay with flow cytometry analyses. E.B. performed cell-cell heterogeneity assay, the 4sgRNA toxicity competition assay, and virus titration in 384-well plates. R.R. helped J-A.Y. or others with plasmid/lentiviral preparation for cell-cell heterogeneity assay, knockout efficiency assay, 4sgRNA toxicity competition assay, autophagy hits validation, and CRISPRoff assay. F.B. helped with autophagy hits validation and performed virus titration of lentiviruses. M.S.B performed puncta area analysis of GFP-SQSTM1 and YFP-LC3. S.B. assisted the method development for comparing activity of 1sgRNA and 4sgRNA for activation. M.L. assisted the flow cytometry analyses of 4sgRNA virus titration and delivery rate with transfection and transduction to non-transfectable cells. S.H. supported the production of Taq DNA ligase and the production of FRET antibodies for PrP^C^ detection. M.K. helped the design of sgRNAs for the 1^st^ trial 384-well plate cloning of 4sgRNA plasmids. L.P. and D.H. supervised research. P.H. supervised research, appropriated the funding, supervised the planning and the execution of the experiments, offered continuous feedback and mentoring at DZNE. A.A. conceived the primary idea of generating arrayed libraries, appropriated the funding, supervised the planning and the execution of the experiments, offered continuous feedback and mentoring, coordinated the activities of the research team, and wrote the paper with input from all authors.

## Acknowledgements

A.A. is the recipient of grants from the Nomis Foundation, the Swiss National Research Foundation, the Swiss Personalized Health Network (SPHN, 2017DRI17), an ERC (European Research Council) Advanced grant, and a donation from the estate of Dr. Hans Salvisberg. J-A.Y. is the recipient of the postdoc grant Forschungskredit from University of Zurich and the Career Development Awards grant of the Synapsis Foundation – Alzheimer Research Switzerland ARS. The work of A.D. J.T., S.R and P.H. was supported by funds from the DZNE. Y.W. and L.Y. were supported by China Scholarship Council. Andrea A. is the recipient of Marie Curie postdoctoral fellowship (MSCA GF, 101033310). We thank Drs. Patrick Hsu (University of California, Berkeley), John Doench (the Broad Institute of MIT and Harvard) and Jacob Corn (ETH Zurich) for their help and suggestions, Dr. Allan Bradley (University of Cambridge) for sharing the Lenti-PB plasmid, Dr. Erwei Zuo (Institute of Agricultural Genome Research, Chinese Academy of Agricultural Sciences) for sharing the transposase plasmid, Dr. David M. Sabatini for suggestions on autophagy hits validation, Dr. Luke A. Gilbert and Mr. Greg C. Pommier (University of California, San Francisco) and Dr. Jonathan S. Weissman (Massachusetts Institute of Technology) for providing reagents and advice for experiments on targeted epigenetic silencing (CRISPRoff). We thank Mr Kevin Maggi, Mr Stefano Sellitto and Mr Gregor Ehmer for technical assistance, Drs. Anna Bratus-Neuenschwander and Weihong Qi, and Dr Susanne Kreutzer and her team at the Functional Genomics Center Zurich (FGCZ) for support with SMRT sequencing, Illumina pair-end sequencing, respectively, and Mr Mario Wickert at the Cytometry Facility of University of Zurich for technical assistance, and Drs. Merve Avar and Daniel Heinzer for supporting C.T. with the PrP^C^ FRET assays.

## Materials and Methods

### DNA constructs

The DNA constructs (except for the 4sgRNA expression plasmids, whose construction by APPEAL will be described separately) used in the study originated from Addgene-deposited plasmids including lentiCas9-Blast (#52962), SP-dCas9-VPR (#63798), lenti-dCas9-VPR-Blast(#96917), PB-TRE-dCas9-VPR (#63800), psPAX2 (#12260), VSV-G (#8454), lentiGuide-Hygro-eGFP (#99375), lentiGuide-Hygro-dTomato (#99376) and pLVX-LC3-YFP (#99571), or as gifts, including the CRISPRoff and CRISPRoff-D3A mutant plasmids by Drs. Luke A. Gilbert and Jonathan S. Weissman from University of California, San Francisco, and the transposase plasmid by Dr. Erwei Zuo from Institute of Agricultural Genome Research, Chinese Academy of Agricultural Sciences, or were constructed in-house (the pYJA5, CRISPRoff-mScarletI, and single sgRNA plasmids).

The pYJA5 construct was created by modifying the lenti-PB vector (Metzakopian et al., 2017) (a gift from Dr. Allan Bradley) in two steps. First, the DNA fragment flanked by the recognition sites for the restriction enzymes *Mlu*I and *Age*I in the lenti-PB vector was replaced by a synthesized DNA fragment that included the human U6 promoter and the fourth variants of tracrRNA, as well as an ampicillin resistance gene (β-lactamase expression cassette). Two *Bbs*I (type II restriction enzyme) recognition sites flanking the β-lactamase expression cassette were introduced into the new fragment, in order to facilitate the removal of the β-lactamase expression cassette. In a second step, the original ampicillin resistance (β-lactamase) expression cassette in the lenti-PB vector was removed between the two *Bsp*HI restriction enzyme recognition sites. After its removal, the insertion of 4sgRNA expression cassettes containing a trimethoprim resistance gene (dihydrofolate reductase) achieves APPEAL cloning. Furthermore, all *BsmB*I recognition sites were mutated. Detailed sequences of the pYJA5 and 4sgRNA-pYJA5 constructs are included in the supplementary information.

The CRISPRoff-mScarletI plasmid was constructed by replacing the TagBFP cDNA fragment with the mScarletI cDNA sequences. The original CRISPRoff plasmid DNA (except the region encoding TagBFP) and mScarletI cDNA from the pmScarlet-i_C1 plasmid (Addgene # 85044) were amplified by polymerase chain reaction (PCR) with the Phusion High-fidelity DNA Polymerase (New England Biolabs #M0530L), then both amplicons were assembled by Gibson assembly to produce the desired CRISPRoff-mScarletI plasmid.

Single sgRNAs were cloned into the pYJA5-modified vector individually via the previously established method (Koike-Yusa et al., 2014).

All plasmids were sequence confirmed.

### Quantification of gene activation by real-time quantitative PCR

HEK293 cells were seeded in 24-well plates at a density of 4.0 × 10^5^ cells per well. The second day, cells at 80–90% confluency were co-transfected with 0.25 µg of single-sgRNA (or 4sgRNA) plasmids and 0.25 µg of dCas9-VPR plasmids using Lipofectamine 3000 (Thermo Fisher Scientific). Three days post-transfection, the cells were lysed, and their total RNA was isolated using TRIzol Reagent (Thermo Fisher Scientific) according to the manual. For iPSCs-derived neurons which stably express dCas9-VPR, 4sgRNA lentiviruses were transduced at a multiplicity of infection (MOI) of 1.4. Seven days post-transduction, cells were lysed, and RNA extracted using the same method as described for the HEK293 cells. 600 ng of RNA was reversed transcribed into cDNA via the QuantiTect Reverse Transcription Kit (Qiagen). Real-time quantitative PCR was done with SYBR green (Roche) according to the manual with the primer sets for each gene as follows. *GAPDH*, *ACTB* and *HMBS* were used as internal controls.

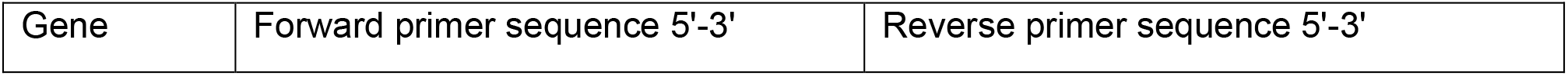
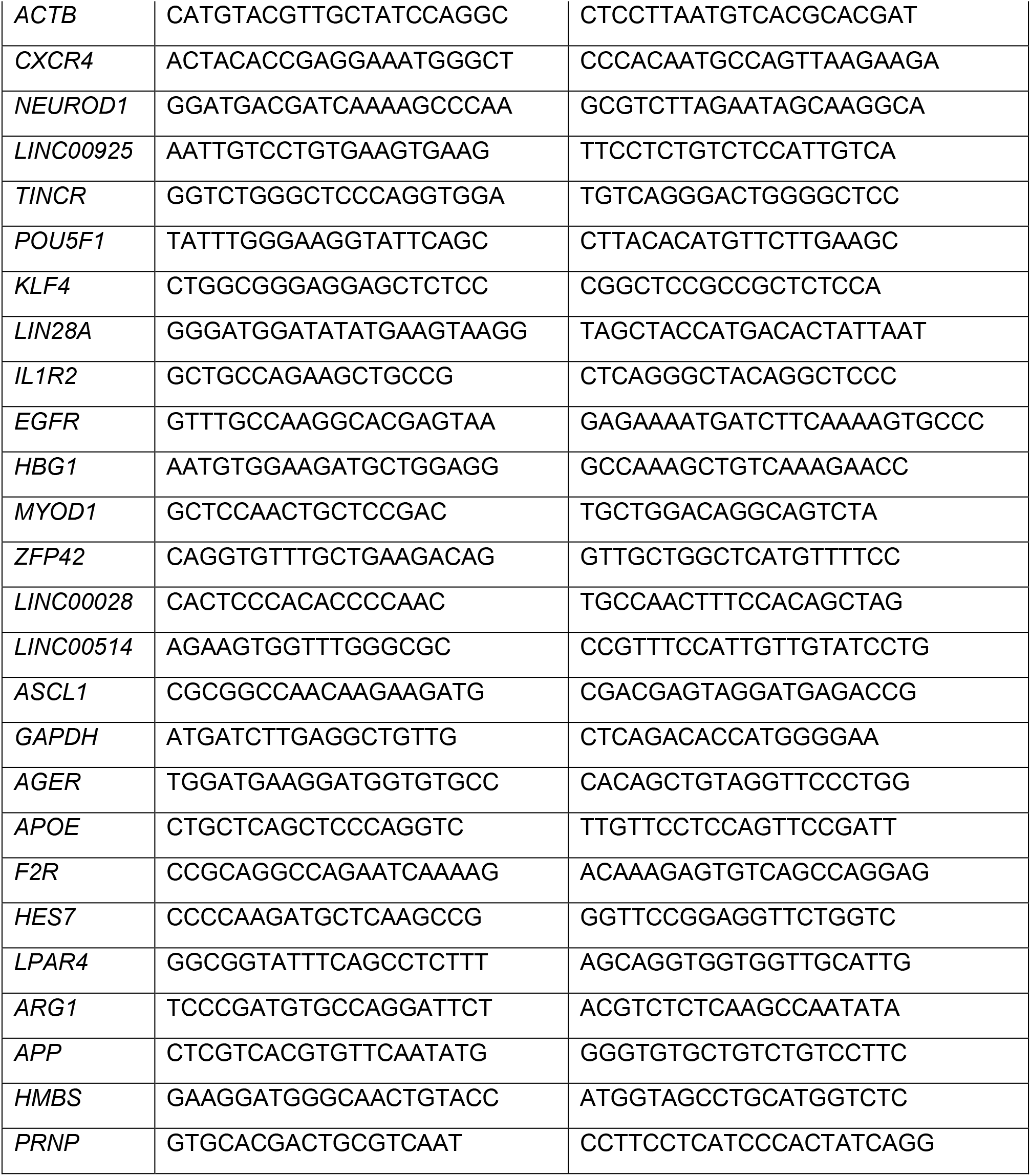

### Gene activation-associated cell-cell heterogeneity assay

Polyclonal HEK293 cells stably expressing doxycycline-inducible dCas9-VPR (PB-TRE-dCas9-VPR, hygromycin selection) were generated by co-transfection of the PB-TRE-dCas9-VPR (500 ng) and transposase (500 ng) plasmids with Lipofectamine 3000 in 6-well plates. Cells were split the next day to Hygromycin B (100 μg/ml, GIBCO) containing medium for 5–7 days. Then the cells were split and transduced with single sgRNAs, 4sgRNAs or non-targeting control (hNTa) lentiviruses at an MOI of 0.3. After 2 days, cells were split with puromycin (1 μg/ml, GIBCO) containing medium and cells were again selected for 5–7 days with medium change every other day. Then cells were seeded in 6-well plates at a density of 2.0 × 10^5^ cells per well. Cells were grown for 3 days in doxycycline hyclate (1μg/ml, Sigma Aldrich) containing medium, and the medium was refreshed every day.

At day 3 post-doxycycline induction of dCas9-VPR expression, cells were trypsinized and resuspended as a single cell suspension in Cell Staining buffer (BioLegend Cat. No. 420201) at a density of 6 × 10^5^ cells in 100 µl. Cells were pre-incubated with 5 µl of Human TruStain FcX™ (Fc Receptor Blocking Solution, BioLegend Cat. No. 422301) per 100µl of cell suspension for 5–10 minutes at room temperature. Afterwards, APC conjugated antibodies to human CD2 (Biolegend, 300214), CD4 (Biolegend, 357408) or CD200 (Biolegend, 329208) were added and incubated on ice for 15–20 minutes in the dark. After 20 minutes, cells were washed twice with 2 ml of Cell Staining Buffer by centrifugation at 350g for 5 minutes. The cell pellet was resuspended in 400 µl of Cell Staining Buffer, transferred to FACS tubes, centrifuged at 1000 rpm for 1 min and finally analyzed with a Fortessa (BD) LSR II analyzer at the core facility center of the University of Zurich. Non-targeting sgRNA-treated cells were used as controls.

### Quantification of gene ablation efficiency by live-cell antibody staining

HEK293 cells at 80–90 % confluency were co-transfected with the lentiCas9-Blast (250 ng per well) and sgRNA plasmids (250 ng per well) using Lipofectamine 3000 in 24-well plates. 24 hours post-transfection, the cells were split to puromycin (1μg/ml) containing medium for 72 hours. Then cells were cultured in medium without selection for around one week. During the whole procedure, cells were maintained at a confluency of no more than 80%. On the day of the assay, cells were trypsinized and single cells were stained with the respective CD47 (BioLegend Cat. No. 323124), IFNGR1 (also known as CD119, BioLegend Cat. No. 308606), or MCAM (also known as CD146, BioLegend Cat. No. 361016) antibody and analyzed using a similar approach as described for the gene activation-associated cell-cell heterogeneity assay mentioned above. Non-targeting sgRNA- treated cells were used as controls.

### Quantification of gene editing efficiency via SMRT long-read sequencing

HEK293 cells were cultured, transfected, and maintained in the same manner as for the gene ablation efficiency measurement by live-cell antibody staining assay. On the day of the assay, instead of being stained with antibodies, the cells were harvested for genomic DNA isolation using the DNeasy Blood & Tissue Kit (Qiagen, 69506). Then barcoded primers (flanking 4sgRNA targeting sites) were synthesized to amplify the genome-edited region of the corresponding genes using Phusion high-fidelity DNA Polymerase (New England Biolabs, M0530L). For each PCR reaction of 50 μl volume, 150 ng genomic DNA, 0.5 μl Phusion DNA polymerase, 10 μM forward/reverse primers, 10 mM dNTP, and 10 μl 5× Phusion HF buffer were used, and the following temperature conditions: Initial denaturation at 98 °C for 30 seconds, 36 cycles at 98 °C for 10 seconds, 60 °C for 30 seconds, and 72 °C for 30 seconds per Kb, and final extension at 72 °C for 6 min. Then the PCR products were purified with gel extraction using the NucleoSpin gel and PCR clean-up kit (Macherey Nagel, 40609.250). Purified PCR amplicons were pooled in approximately equal molar amounts (determined by Nanodrop) and subjected to SMRT long-read sequencing. The individual reads were demultiplexed and aligned to their corresponding amplicon sequences amplified from cells treated with non-targeting sgRNAs. Gene editing efficiency was calculated as the percentage of mutated reads compared to the corresponding non-targeting controls. For each gene, their Fwd bc1, bc2, bc3, and bc4 primers was designed with 4 distinct 10-bp barcodes (lower case nucleotides in the following table) to index the four biological repeats, and their Rev primers bc1 and bc2 were designed with 2 distinct 10-bp barcodes (lower case nucleotides in the following table) to index non-targeting and 4sgRNAplasmid-s transfected cells, respectively.

The barcoded primers used were HPLC purified and are listed below:

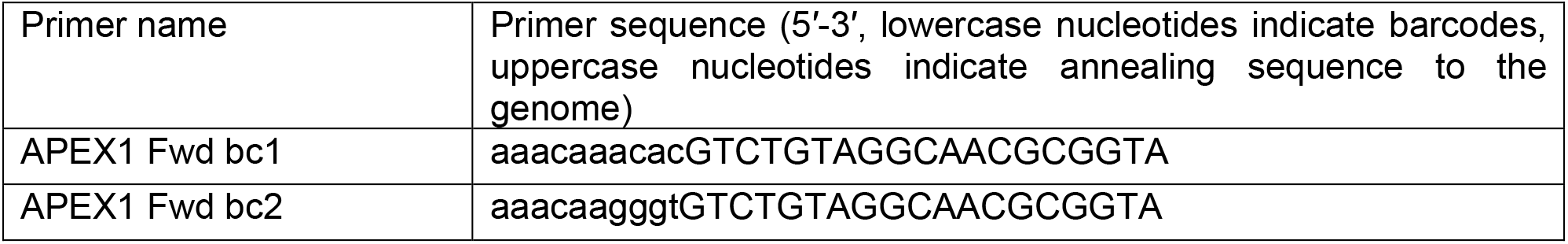
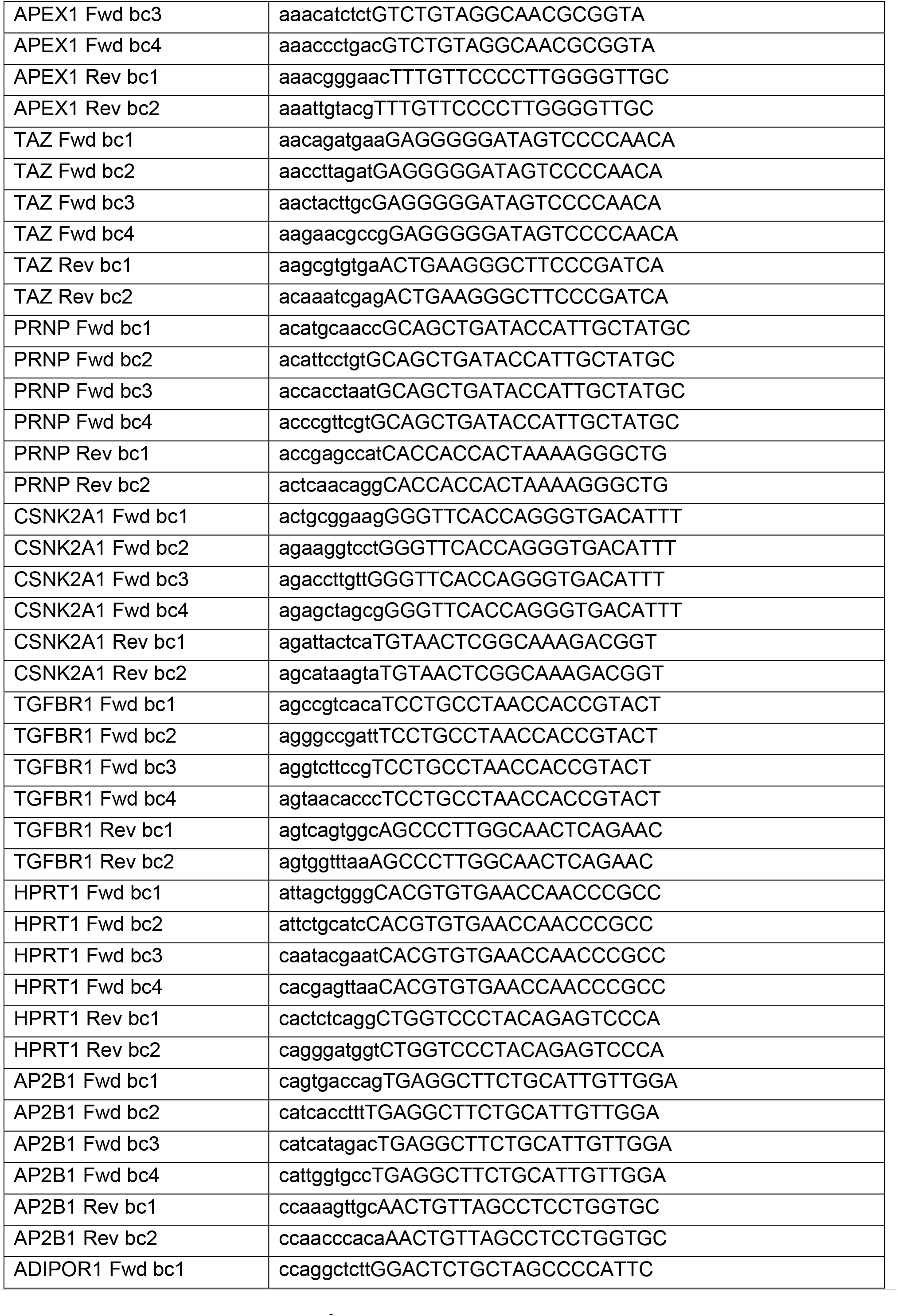
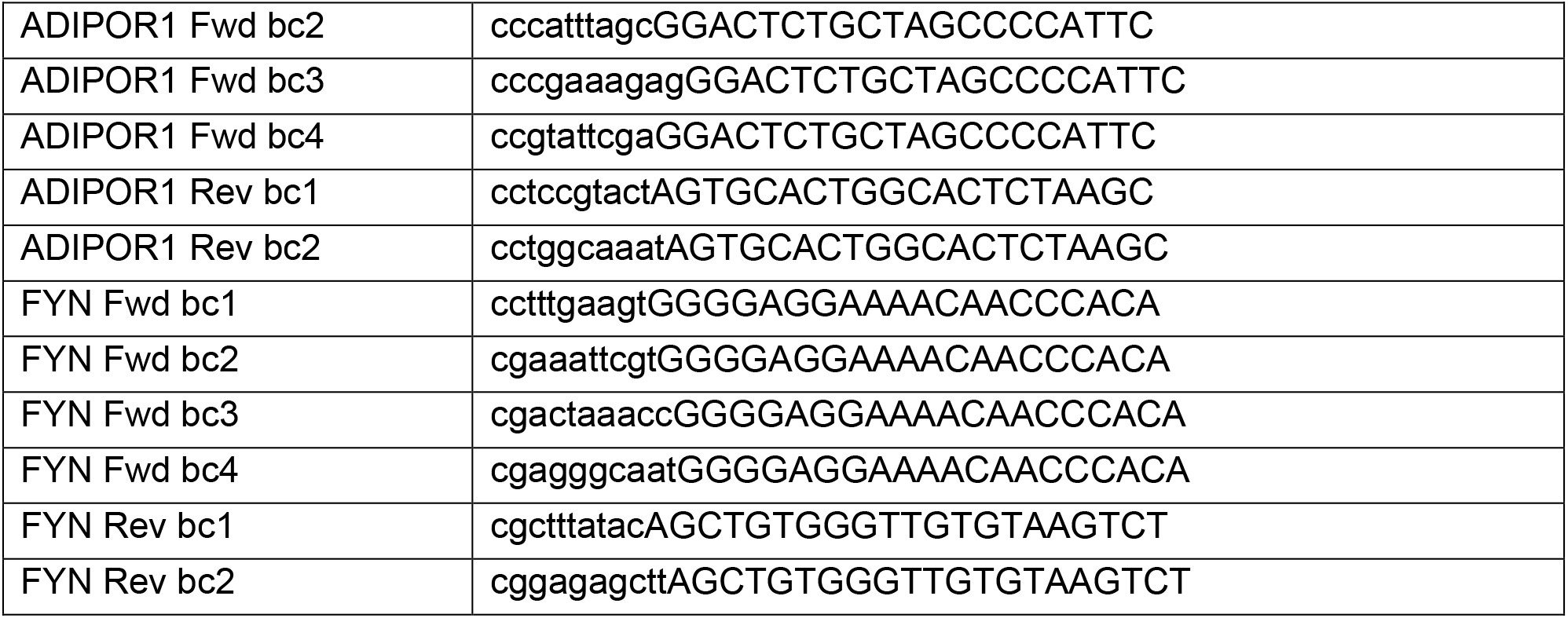

### Cell-growth competition assay

HEK293 cells stably expressing dTomato were generated by transducing the lentiGuide-Hygro-dTomato lentivirus. HEK293-Cas9-EGFP cells were generated by co-transducing HEK293 cells with lentiCas9-Blast and lentiGuide-Hygro-eGFP lentiviruses. Non-transduced cells were eliminated with hygromycin (100 μg/ml, GIBCO) and blasticidin (10 μg/ml, GIBCO). Polyclonal cells of both lines were used in the assay. The two stable cell lines were then mixed and seeded in a ratio of around 1:1. The next day, cells were analyzed with a Fortessa (BD) LSRII analyzer at the core facility center of the University of Zurich to validate the ratio of EGFP/dTomato of the starting cells. Afterwards, cell mixtures were seeded on 12-well plates and transduced at an MOI of 0.3 with lentivirus containing different 4sgRNA knockout plasmids targeting different genes or non-targeting controls. Two days post-transduction, cells were split with puromycin (1μg/ml, GIBCO) containing medium and selected for 5–7 days with medium change every other day. At day 14 after the lentiviral transduction of sgRNAs, cells were collected, resuspended in PBS, transferred to FACS tubes, centrifuged at 1000 rpm for 1 minute and analyzed with a Fortessa (BD) LSR II analyzer at the core facility center of the University of Zurich to determine the ratio of EGFP/dTomato positive cells. The final ratio was normalized to the starting ratio measured on day 2 after seeding the cell mixtures.

### Lentiviral packaging of individual or small number of plasmids

HEK293T cells were grown at 80–90% confluency in DMEM + 10% FBS medium on poly-D-lysine-coated 24-well plates and transfected with the 3 different plasmids (sgRNA plasmid, psPAX2, and VSV-G; ratios: 5:3:2) with Lipofectamine 3000 for lentivirus production. After 6 hours, or overnight incubation, the medium was changed to virus harvesting medium (DMEM + 10% FBS + 1% BSA). The supernatant containing the lentiviral particles was then harvested 48–72 hours after the change to virus harvesting medium. Suspended cells or cellular debris was pelleted with centrifugation at 1500 rpm for 5 min. Then clear supernatant was collected and stored at −80 °C.

For the titration of the lentiviral particles, 2 × 10^5^ HEK293T cells were grown in 24-well plates, and infected by adding small volumes (V) of the abovementioned viral supernatant (-e.g. 3 µl). A representative batch of cells was used to determine the cell count at the time of infection (N). 72 hours after infection, the cells were harvested and analysed by flow cytometry to quantify the fraction of infected cells (BFP positive). The percentage of positive cells (P) is then used to calculate the titre (T) of the virus according to the following formula: 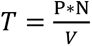

### High-throughput lentiviral production in 384-well plates

#### Cell seeding

A monoclonal HEK293T cell line that produces higher titers of lentivirus was generated in-house and used for high-throughput lentiviral production. 6,800 cells were seeded with Integra Viaflo in 50 µl of DMEM + 10% FBS medium per well.

#### Plasmid transfection

Around 24 hours after seeding, when cells reached 80–90% confluency, the mix was prepared according to the following table (the amount was for three 384-well plates of plasmid set, each plasmid set for three technical repeats, each plasmid set a 384-well plate of plamids) for transfection:

|  | Opti-MEM | psPAX2 | VSV-G | P3000 | Lipofectamine |
| --- | --- | --- | --- | --- | --- |
| Tube 1 | 14.7 ml | 69.34 µg | 46.23 µg | 441.3 µl |  |
| Tube 2 | 18.3 ml |  |  |  | 441.3 µl |

Cells were transfected with the following procedure within 1 hour after the mix had been prepared:

1. Tube 1 was prepared and 38 µl of the mix was transferred to each well of a 384-deep-well-plate using a multichannel pipette (plate with Mix 1).
2. 10.1 µl (per well) of the above-mentioned Mix 1 was then transferred to three 384-well PCR-plates using the Viaflo (12.5 µl-tips).
3. Tube 2 was prepared and 46 µl (per well) of the mix was transferred to a 384-deep-well-plate using a multichannel pipette.
4. 72 ng (2.4 µl of 30 ng/µl dilution) of 4sgRNA plasmid (from the library in 384-well-format) was added to the plate prepared at Step 2 and mixed for 10 cycles (Viaflo; speed up/down: 8), then incubated for 5 minutes at room temperature (RT).
5. 12.5 µl (per well) of the Lipofectamine 3000 mix prepared at Step 3 was transferred to the plates prepared at Step 4 to obtain 25 µl of transfection mix per well and mixed for 20 cycles (speed up/down: 8), then incubated for 20 minutes at RT.
6. 36 µl of medium was removed from the cell culture plate (ViaFlo setting: 31.5 mm; speed up 3).
7. The plate prepared at Step 5 was centrifuged at 1500 rpm for 2 minutes.
8. 8 µl of transfection mix was added to each replicate of the three replicate cell culture plates (ViaFlo setting: 16 mm; speed down 3); and incubated at 37 °C for 6–12 hours.

#### Virus harvesting medium addition

Virus Harvesting Medium (composition for a 500 ml: 434 ml of DMEM + 59 ml of 10% FBS + 6.82 grams of BSA + 6.8 ml of penicillin or streptomycin) was prepared and warmed up to 37 °C. Then 44 µl per well of Virus Harvesting Medium was added to the plates with transfected cells (ViaFlo settings: 37mm, speed up: 8, speed down: 2).

#### Virus collection

48–60 hours after the addition of the virus harvesting medium, 45 µl of viruses of each replicate transfected plate was collected, and viruses from the three replicates of the same plasmid set were pooled together, mixed for 20 cycles and then centrifuged at 1500 rpm for 2 minutes to pellet cells. Viral supernatant was aliquoted into PCR plates for storage at −80°C, for later use and titer determination.

#### Virus titration

One day before virus titration, 6,000 HEK293T cells were seeded in 384-well plates. Lentiviruses produced in 384-well plates were thawed from −80 °C and 0.5 µl of viruses (per well) were transduced into the HEK293T cells. Three days post-transduction, cell culture medium was removed, and cells were dissociated with 15 µl of PBS-EDTA (2 mM EDTA in 1× PBS) for 15–20 min at 37 °C. Later, 20 µl of FACS buffer (10 mM EDTA + 5% FBS in 1× PBS) was added to each well and mixed with the Viaflo until a single-cell suspension was obtained. Then the percentage of TagBFP-positive cells was analysed by flow cytometry.

### APPEAL high-throughput generation of libraries

#### Oligo synthesis

Twenty-nucleotide sgRNA sequences were incorporated into oligonucleotide sequences with appended constant sequences and synthesized in 384-well plates using the high affinity purification (HAP) purification method by Sangon Biotech (China). The sgRNA1 (sgRNA1 sequence, N_20sg1_) oligonucleotide sequence is: 5′- ttgtggaaaggacgaaacaccGN_20sg1_GTTTAAGAGCTAAGCTG-3′; the sgRNA2 (sgRNA2 sequence, N_20sg2_) oligo sequence is: 5′- cttggagaaaagccttgtttGN_20sg2_GTTTGAGAGCTAAGCAGA-3′; the sgRNA3 (sgRNA3 sequence, N_20sg3_) oligo sequence is: 5′ - gtatgagaccactctttcccGN_20sg3_GTTTCAGAGCTAAGCACA-3′; and the sgRNA4 (reverse complement sequence of sgRNA4, N_20 crsg4_) oligo sequence is 5′ - ATTTCTGCTGTAGCTCTGAAACN_20crsg4_Cgaggtacccaagcggc-3′. The oligonucleotides were diluted with ultrapure water to a working concentration of 4 μM.

#### Three-fragment PCRs

A total of 10 µL PCR reaction per well was produced in 384-well plates. The C1 fragment (amplicon size 761 bp) PCR mix was prepared as follows:

| Volume/well | Reagent |
| --- | --- |
| 0.2 μL | C1 fragment 1 ng/μL |
| 0.2 μL | mU6 Rev primer 10 μM<br>(common primer, sequence attached) |
| 2 μL | 5X HF buffer |
| 0.2 μL | dNTPs 10 mM |
| 0.1 μL | Phusion High-Fidelity DNA polymerase |
| 6.8 μL | ddH<sub>2</sub>O |

9.5 µL of the mix were aliquoted in each well of the 384-well plate, and 0.5 µl of sgRNA1 primer (at 4 µM concentration) was added to each well and mixed.

The M fragment (amplicon size 360 bp) PCR mix was prepared as follows:

| volume/well | Reagent |
| --- | --- |
| 0.2 μL | M fragment 1 ng/μL |
| 0.2 μL | M Rev primer 10 μM<br>(common primer, sequence attached) |
| 2 μL | 5X HF buffer |
| 0.2 μL | dNTPs 10 mM |
| 0.1 μL | Phusion High-Fidelity DNA polymerase |
| 6.8 μL | ddH<sub>2</sub>O |

9.5 µL of the mix were aliquoted in each well of the 384-well plate, and then 0.5 µl of sgRNA2 primer (at 4 µM concentration) was added to each well and mixed.

C2s fragment (amplicon size 422 bp) PCR mix was prepared as follows:

| vol/well | Reagent |
| --- | --- |
| 0.2 μL | C2s fragment 1ng/μL |
| 2 μL | 5x HF buffer |
| 0.2 μL | dNTPs 10 mM |
| 0.1 μL | Phusion High-Fidelity DNA polymerase |
| 6.5 μL | ddH<sub>2</sub>O |

9 µL of the mix were aliquoted in each well of the 384-well plate, and then 0.5 µL of sgRNA3 primer (at 4 µM concentration) and 0.5 µL of sgRNA4 primer (also at 4 µM concentration) were added to each well and mixed.

The Integra ViaFlo 384-well pipetting system was used for all 384-well liquid handling. All PCR plates were sealed tightly and centrifuged at 2000 rpm for 2 minutes, and placed in thermocyclers with the following program: Preheat the lid at 99 °C; Initial denaturation at 98 °C for 30 seconds, 36 cycles comprising 98 °C for 10 seconds, 60 °C for 30 seconds, and 72 °C for 25 seconds, and final extension at 72 °C for 5 minutes, followed by cooldown to 20 °C. All PCR products were then diluted with 9 µL of ultrapure water for later Gibson assembly. The success of PCR on each plate was confirmed by DNA agarose gel electrophoresis of several random samples on the plate.

#### Gibson assembly

Assembly of the three fragment PCR products into the pYJA5 vector was performed in a 384-well plate by Gibson assembly, with the following reaction mix:

| Volume | Reagent |
| --- | --- |
| 2 μL | C1 amplified fragment (estimated around 16 ng/μL) |
| 1 μL | M amplified fragment (estimated around 16 ng/μL) |
| 1 μL | C2s amplified fragment (estimated around 20 ng/μL) |
| 1 μL | pYJA5 *Bbs*I digested purified vector, diluted to 120 ng/μL |
| 5 μL | 2x homemade HiFi Gibson master mix |

The mix was incubated in the thermocycler at 50 °C for 1 hour, and then used for the transformation of competent cells or stored immediately at −20 °C.

#### Transformation and bacterial storage

Transformation was carried out in 96-well deep-well plates (2.3 mL, Axygene P-DW-20-C) in the cold room. 5 µL (per well) of Gibson mix from the 384-well plate was transferred into four 96-well plates and spun down to the bottom of each well. 50 µL (per well) of homemade competent cells (NEB stable competent cells) were dispensed and mixed twice with the Gibson mix. The plates were then kept immersed in ice for 30 minutes. Heat shock was performed for 30 seconds at 42 °C by placing the plate into a water bath. Plates were placed back on ice for 5 minutes. 300 µL of homemade SOC medium (0.5% Yeast Extract, 2% Tryptone, 10 mM NaCl, 2.5 mM KCl, 10 mM MgCl_2_, 10 mM MgSO_4_, 20 mM Glucose) were then added to the plate and incubated for 1 hour at 37 °C under shaking at 900 rpm using a thermo-shaker. Then, 900 µL (per well) of Terrific Broth (TB) medium (https://openwetware.org/wiki/Terrific_Broth) containing 15 µg/mL trimethoprim and 15 µg/mL tetracycline was added to the transformation mix, and incubated at 30 °C under shaking at 900 rpm for 40–48 hours.

Bacteria were then stored at a final concentration of 16.7% (v/v) glycerol in both 96--well plates (300 µL final storage volume) and 384-well plates (150 µL final storage volume) at −80 ℃.

#### Magnetic bead-based 96-well plasmids miniprep

Fifty microliters of the Gibson assembly product-transformed bacteria were transferred into 1.2 mL of TB medium (with 15 µg/mL trimethoprim and 15 µg/mL tetracycline in 96-well deep well plate) immediately before the storage of the bacteria, and grown at 30 °C at 900 rpm for 40–48 hours. The bacteria were then subjected to in-house magnetic beads-based plasmids miniprep procedures, which were adopted from the canonical plasmids miniprep protocols (Birnboim and Doly, 1979). Briefly, the bacteria were pelleted at 4000 rpm for 10 min and resuspended in 200 µl of P1 buffer [50 mM glucose, 10 mM EDTA, 25 mM Tris (pH 8.0)], and subsequently lysed in 200 µl of P2 buffer [0.2 M NaOH, 1% SDS (w/v)], and the lysis mixture was neutralized in 200 µl of P3 buffer (3 M KOAc, pH 6.0) and subjected to centrifugation at 4000 rpm for 10 min at 4 °C. Then 400 µl of the supernatant were transferred into a new deep-well plate and 1000 µl of cold absolute ethanol were added and mixed, then centrifuged at 4000 rpm for 10 min at 4 °C. The supernatant was discarded and 50 µl of ddH_2_O was added to the plasmid pellet and mixed to dissolve the plasmids. Then 75 µl of beads buffer (2.5 M NaCl, 10 mM Tris base, 1mM EDTA, 3.36 mM HCl, 20% (w/v) PEG8000, 0.05% (w/v) Tween 20) and 50 µl of SpeedBeads™ magnetic carboxylate modified particles (GE Healthcare 65152105050250, 1:50 dilution in beads buffer) were added to the plasmids, mixed and incubated for 5 min on a magnetic rack to separate the beads from the supernatant. The beads were then washed twice with 70% ethanol and dried in a water bath (65 °C). Plasmid DNA was then eluted by 150 µl of sterile tris-EDTA buffer [1 mM EDTA, 10 mM Tris-HCl (pH 8.0)] from the beads at 65 °C for 10 min and transferred to a new low-profile 96-well plate. To ensure the full cloning procedure was correct, two wells of plasmids from each 96-well plate were subjected to Sanger sequencing.

### In silico 4sgRNA libraries design

#### Pooling existing libraries

To provide a starting point for guide RNA selection, we collected sgRNAs from previously published and validated libraries and tools, each of which employed their own algorithms to select sgRNAs with high predicted on-target efficacy. We included the Calabrese (Sanson et al., 2018) and hCRISPRa v2 (Horlbeck et al., 2016) libraries for CRISPRa, and the TKOv3 (Hart et al., 2017) and Brunello (Doench et al., 2016; Sanson et al., 2018) libraries for CRISPRko. We complemented these source libraries with sgRNAs from the CRISPick tool (formerly GPP sgRNA Designer, https://portals.broadinstitute.org/gppx/crispick/public), to ensure optimal coverage of difficult-to-target and newly annotated genes (the website was accessed in April 2020, following the update of 20 March 2020).

#### Gene definitions

Entrez gene identifiers were used to provide common gene definitions for sgRNAs from all sources. If the source library did not provide Entrez identifiers, the official gene symbols were mapped to Entrez IDs, and the genomic location was used to disambiguate gene symbols when necessary. Genes that were not defined as protein-coding by NCBI or Ensembl were excluded according to the following annotation files:

ftp://ftp.ncbi.nih.gov/gene/DATA/GENE_INFO/Mammalia/Homo_sapiens.gene_info.gz ftp://ftp.ensembl.org/pub/release-99/tsv/homo_sapiens/Homo_sapiens.GRCh38.99.entrez.tsv.gz

Both files were downloaded on 25 March 2020. The final libraries included 19839 protein-coding genes for CRISPRa and 19820 for CRISPRko; the difference in gene counts arises from genes that were present for only one modality in our source libraries. For example, highly polymorphic genes related to adaptive immunity, such as the T Cell Receptor Alpha Locus (TRA) gene, are available for CRISPRa, but not CRISPRko.

#### Transcriptional start site (TSS) definitions

To ensure good coverage of alternative transcripts and the broad applicability of the CRISPRa library in multiple cell lines, we adopted the alternative transcription start site (TSS) definitions from the hCRISPRa-v2 library (Horlbeck et al., 2016). The authors of this library used the FANTOM5 CAGE-seq dataset (Consortium et al., 2014), supplemented by Ensembl (Yates et al., 2020) transcript models, to define TSS positions. Additional TSSs were targeted by a separate set of sgRNAs if the FANTOM5 scores indicated significant transcriptional activity, and if they were spaced more than one kilobase apart from the primary TSS. We chose a separate set of four sgRNAs for each TSS, treating multiple TSSs as if they were distinct genes. To group sgRNAs by TSS, we mapped sgRNAs from all sources (including the top five sgRNAs from the CRISPick sgRNA Designer) to their genomic locations. We then iterated through each sgRNA, starting with the lowest genomic coordinate; a new TSS group was defined if the distance from one guide to the next exceeded 1000 base pairs. Additional TSSs were only targeted if a valid combination of four guides was available. Multiple TSSs were included for 2,311 genes using 4,803 four-guide combinations, whereas a single TSS was targeted for the remaining 17,528 genes.

#### Avoidance of genetic polymorphisms

For each sgRNA, we checked for overlaps with regions of frequent genetic polymorphism in human populations. We considered both the 20-nucleotide target sequence and the two guanosine nucleotides of the NGG PAM; however, the first nucleotide of the PAM was allowed to vary. We avoided sgRNAs whose target region contained any genetic polymorphisms with frequencies greater than 0.1%. Variant frequencies were derived from the Kaviar database (Glusman et al., 2011), which includes curated genomic data on single nucleotide variants, indels, and complex variants from over 77,000 individuals (including over 13,000 whole genomes). The dataset (only variants seen more than 3 times, version 160204-hg38) was downloaded on 7 August 2019. The polymorphism frequencies in the Kaviar database were generally similar to those from TOPMED, gnomAD, and the 1000 Genomes Project.

#### Specificity scores

In order to select a four-guide combination with minimal off-target effects, we computed specificity scores for each sgRNA from our source libraries. We used the approach introduced by the authors of the GuideScan (Perez et al., 2017) tool: For each guide, potential off-target sites were weighted by their CFD (cutting frequency determination) scores (Doench et al., 2016), and CFD scores were aggregated into a single score using the formula: 1 / (1 + sum of CFD scores from all off-target sites (Hsu et al., 2013). Because the pre-computed GuideScan Cas9 database does not contain all sgRNAs (it excludes those with perfect-match or one-mismatch off-target sites in the reference genome), we annotated sgRNAs using both the GuideScan and CRISPOR (Concordet and Haeussler, 2018; Haeussler et al., 2016) tools. Local installations of these tools were used, and the source code was downloaded in December 2020 (GuideScan version 2018-05-16, and CRISPOR version 4.97). The output of the local installations was confirmed to be identical to that of the web-based tools. When available, we used GuideScan specificity scores (considering up to three mismatches); otherwise, CRISPOR specificity scores were used (considering up to four mismatches). CRISPOR three-mismatch (3MM) and four-mismatch (4MM) specificity scores analogous to those from GuideScan were computed, using the detailed output files listing each off-target site. GuideScan and CRISPOR specificity scores were highly correlated, but not identical, due to slight differences in the number of off-target sites identified for the same sequence. When selecting sgRNAs, we avoided low-specificity guides with 3MM scores below 0.2; this cut-off point was recently shown to have good predictive power for identifying sgRNAs with significant off-target activity (Tycko et al., 2019). However, this criterion had to be relaxed in cases where all eligible sgRNAs had specificity scores below 0.2, for example, when targeting genes present in multiple copies in the genome, or those belonging to large gene families with many closely related paralogs and pseudogenes. Finally, in order to choose among all eligible four-guide combinations, we computed an aggregate specificity score using the formula: 1 / (1 + sum of CFD scores from all four guides), and picked the combination with the highest score, indicating high predicted specificity.

#### sgRNA spacing

To allow for unhindered multiple binding for synergistic effect, we selected, whenever possible, four sgRNAs whose “cut” locations were spaced at least 50 base pairs apart. However, for CRISPRa, target sequences should be located within a window of about 400 base pairs upstream of the TSS for optimal activity (Gilbert et al., 2014), which is reflected in the selection of sgRNAs in the source libraries. Thus, overlaps were unavoidable for some genes. For CRISPRko, on the other hand, overlaps were often inevitable when targeting genes with very short coding sequences. In those cases, we nevertheless aimed to minimize the total number of overlaps between neighbouring guides. Furthermore, all four-guide combinations strictly adhered to another criterion: No two sgRNAs were allowed to share identical sub-sequences of more than seven base pairs. This was done primarily to minimize recombination events between identical regions during Gibson assembly of the plasmid. However, this also enforced minimal spacing of the four selected guides.

#### Selection of four sgRNAs

After integration and annotation of sgRNAs from the source libraries, we selected the final combination of four sgRNAs for each gene or TSS. First, sgRNAs containing a stretch of four or more T nucleotides were excluded, since this sequence can induce termination of transcription. Next, all possible four-guide combinations for each gene were generated, and combinations that shared identical subsequences greater than seven base pairs in length were excluded. The potential combinations were then ranked, using a list of criteria that were applied in order; if multiple combinations were tied in first place, the decision was made using the next criterion down the list. The criteria were as follows: 1) Maximize the number of sgRNAs (from zero to four) that fulfil certain minimal requirements – the sgRNA can be mapped to a defined genomic location in the reference genome with an N(GG) PAM; there are no overlaps with frequent genetic polymorphisms (>0.1%); the 3MM specificity score is at least 0.2; and for CRISPRko only, the guide conforms to the criteria of Graf et al. (Graf et al., 2019); 2) maximize the number of sgRNAs with exactly one perfect match location in the reference genome, 3) minimize the number of overlaps between two neighbouring sgRNAs spaced fewer than 50 base pairs apart, 4) minimize the number of sgRNAs derived from the CRISPick sgRNA Designer tool, rather than the previously published libraries, 5) for CRISPRa, minimize the number of sgRNAs derived from the “supplemental 5” rather than “top 5” sgRNAs for the hCRISPRa-v2 library, and for CRISPRko, minimize the number of CRISPick-derived sgRNAs ranked outside the top 10, and 6) maximize the aggregate specificity score from all 4 guides. The highest-ranked four-guide combination was chosen. Since the aggregate specificity score was the only quantitative criterion, it acted as a tiebreaker and had the greatest impact on the choice of guides.

#### Sublibrary allocation

To facilitate focussed screens of a subset of the genome, we divided the entire set of protein-coding genes into mutually exclusive sub-libraries. Two of our sub-libraries – Transcription Factors, and Secretome – were based on recent publications that combined bioinformatics analyses with expert curation to arrive at a comprehensive list of genes in those categories (Lambert et al., 2018; Uhlen et al., 2015). These lists were obtained from the publication’s supplemental data (for the secretome) or the authors’ website (for the transcription factors; humantfs.ccbr.utoronto.ca, database version 1.01). Ensembl gene IDs were translated to Entrez gene IDs, making use of HUGO gene symbols to disambiguate one-to-many mappings for a few genes. A third sub-library was based on a list of G-protein coupled receptors, curated by the HUGO Gene Nomenclature Committee (HGNC) (Braschi et al., 2019) (https://www.genenames.org/cgi-bin/genegroup/download?id=139&type=branch, accessed on 11 Marc 2020). An additional seven thematic sub-libraries were adopted from the hCRISPRa-v2 library (Horlbeck et al., 2016): Membrane Proteins, Kinases/Phosphatases/Drug Targets, Mitochondria/Trafficking/Motility, Stress/Proteostasis, Cancer/Apoptosis, Gene Expression, and Unassigned. The first two of these thematic sub-libraries were updated to incorporate a small number of additional transmembrane receptors, transporters, kinases and phosphates, using Gene Ontology terms (exported from BioMart (Smedley et al., 2009) on 25 March 2020) and a list of membrane proteins provided by the Human Protein Atlas project (Uhlen et al., 2019) (https://www.proteinatlas.org/search/protein_class:Predicted+membrane+proteins, accessed on 11 March 2020). If a gene belonged to multiple categories, it was assigned to the first sub-library (in the order in which they are listed in this section), and all remaining genes were added to the Unassigned sub-library.

#### Classification of unintended gene perturbations

Some sgRNAs are expected to perturb additional genes other than the intended target gene. In certain cases, this occurs at a different locus than the intended target site: For example, gene families of very close paralogs can often only be targeted with sgRNAs that have multiple perfect-match binding sites in the genome. However, in most cases, this involves a single locus – the intended binding site – where a sgRNA may perturb more than one gene. In the case of CRISPRa, the same promoter region is often shared by two genes located on opposite strands of the chromosome, so that their transcription start sites (TSSs) lie only a few hundred base pairs apart. In this case, guide RNAs that effectively activate one gene would inevitably also activate the other. As a guide for users of the library, and to aid the interpretation of hit genes, we annotated sgRNAs with a complete list of all genes they target. For the purpose of summarizing this phenomenon across the entire library, we classified each sgRNA as 1) only targeting the intended gene, 2) targeting unintended genes, but in a single location (on-site off-target effects), or 3) targeting unintended genes at other locations (off-site off-target effects). If two perfect-match sgRNA binding sites had any target genes in common, they were considered to target unintended genes at the same location. This ensured that sgRNAs targeting the pseudoautosomal region of chromosomes X and Y were classified correctly. Genes in this region technically map to two different chromosomes, but they are present in only two copies in diploid cells, just like any other gene. Unless they have additional target sites, they should not be included in the “off-site” category.

#### Annotation of unintended target genes

To annotate each sgRNA with all its potential target genes, a database of TSS locations was constructed by merging the FANTOM5 dataset (lifted over to the hg38 genome (Abugessaisa et al., 2017), version 3) with data from BioMart (Smedley et al., 2009) (exported on 25 March 2020), using Entrez gene IDs as a common identifier. Similarly, data on coding sequence (CDS) and exon locations were compiled from BioMart, the “TxDb.Hsapiens.UCSC.hg38.knownGene” Bioconductor package (version 3.10.0), and GENCODE (Frankish et al., 2019) annotation data (Release 33), and location data were merged using Entrez gene identifiers (if available) or Ensembl gene identifiers. Genes annotated as pseudogenes, or whose categorization was unclear, were excluded from further analysis. For CRISPRa, perfect-match sgRNA binding sites within a window of 1000 base pairs around TSSs were considered. For CRISPRko, sgRNA cut locations had to lie within the coding sequences (CDSs) of protein-coding genes, or within the exons of non-coding RNAs.

#### Annotation of predicted deletions

In the case of CRISPRko, when four sgRNAs are active within the same cell, the multiple, closely spaced double-strand breaks commonly lead to the loss of a DNA segment between the sgRNA cut locations. Thus, in addition to annotating individual sgRNAs, we also determined which genes are affected by the predicted deletion – the segment between the first and last cut site. We also took deletions induced by (perfect-match) off-target binding sites into consideration. Because deletions may be less likely to occur if the cut sites are very far apart, we imposed a maximum distance of one megabase between cut sites, so that multiple predicted deletions (or isolated cut positions) on the same chromosome were possible.

#### In silico comparison of CRISPR libraries

To compare in silico characteristics of existing libraries and the 4sgRNA libraries, the top four guides per gene were selected. Whereas the Brunello (Doench et al., 2016; Sanson et al., 2018) and TKOv3 (Hart et al., 2017) libraries were designed to contain four sgRNAs per gene, the Calabrese library (Sanson et al., 2018)was divided by the authors into Set A and Set B, each containing three sgRNAs per gene. To define the top four sgRNAs, the sgRNAs from Set A were supplemented with a randomly selected sgRNA from Set B. For the hCRISPRa-v2 (Horlbeck et al., 2016) and CRISPick tool, the four highest-ranked sgRNAs were chosen (using the “Pick Order” column in the output from the CRISPick sgRNA designer tool). Since the libraries differed in the genes they covered, and since different genes vary in the availability of potential sgRNAs with high predicted activity and specificity, only genes present in all libraries were used for benchmarking. Furthermore, for genes for which the T.gonfio and hCRISPRa-v2 libraries included more than one transcription start site (TSS), only the sgRNAs targeting the main TSS was included, defined as the TSS with the highest score in the FANTOM5 dataset, or – if data were unavailable for that gene – the most upstream TSS. To compare the expected number of sgRNA binding sites affected by genetic polymorphisms, the frequency of the most common polymorphism overlapping each sgRNA was considered. This is a conservative estimate, since SNPs with frequencies below 0.1% were excluded. Furthermore, in the case of multiple single nucleotide polymorphisms (SNPs) overlapping with a sgRNA, only the most frequent was considered. Due to linkage disequilibrium between SNPs affecting the same sgRNA, a precise estimation of the total probability of overlaps with polymorphisms would require access to the individual-level sequencing data underlying the SNP databases.

#### Software and code

The annotation and selection of sgRNAs for the library design was performed using the R statistical programming environment (R Core Team, 2020), version 3.6.3, and the Bioconductor suite (Huber et al., 2015), version 3.10. Source code is available at https://github.com/Lukas-1/CRISPR_4sgRNA.

### SMRT long-read next-generation sequencing of libraries

#### Barcoding, amplification, and long-read sequencing

Library plasmids were firstly diluted to be 1.3 ng/µl and then 0.5 µl of each diluted plasmid was used as template for per 10 µl PCR reaction with barcoded primers that uniquely identified each plasmid on a 384-well plate. This was achieved using a combination of 16 different forward primers and 24 different reverse primers (distinguishing the rows and columns of the 384-well plate, respectively). The amplified region was 2225 base pairs in length, encompassing the entire 4sgRNA expression cassette (containing all four promoter, guide RNA and tracrRNA sequences, as well as the trimethoprim resistance element), and was flanked by two 10-bp paired barcode sequences. The high-fidelity Phusion DNA polymerase was used and PCR was performed with amplification of 16 cycles. Then PCR amplicons of the same 384-well plate was pooled down and purified by magnetic beads, then individual purified pools were uniquely barcoded, purified, and further pooled with near equal molar of DNA amplicons and subjected to single-molecule real-time (SMRT) sequencing data using the PacBio Sequel IIe instrument.

The barcoded primers were HPLC-purified and are listed below (lowercase nucleotides indicate barcodes, uppercase nucleotides indicate the sequence annealing to the plasmid vector):

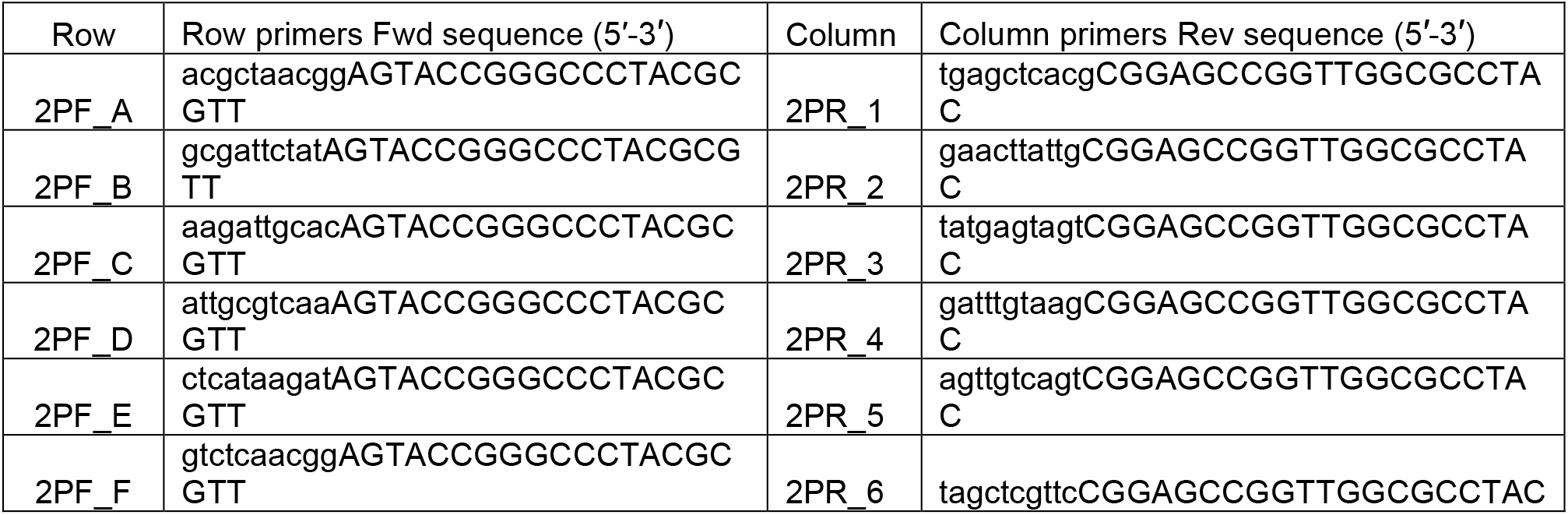
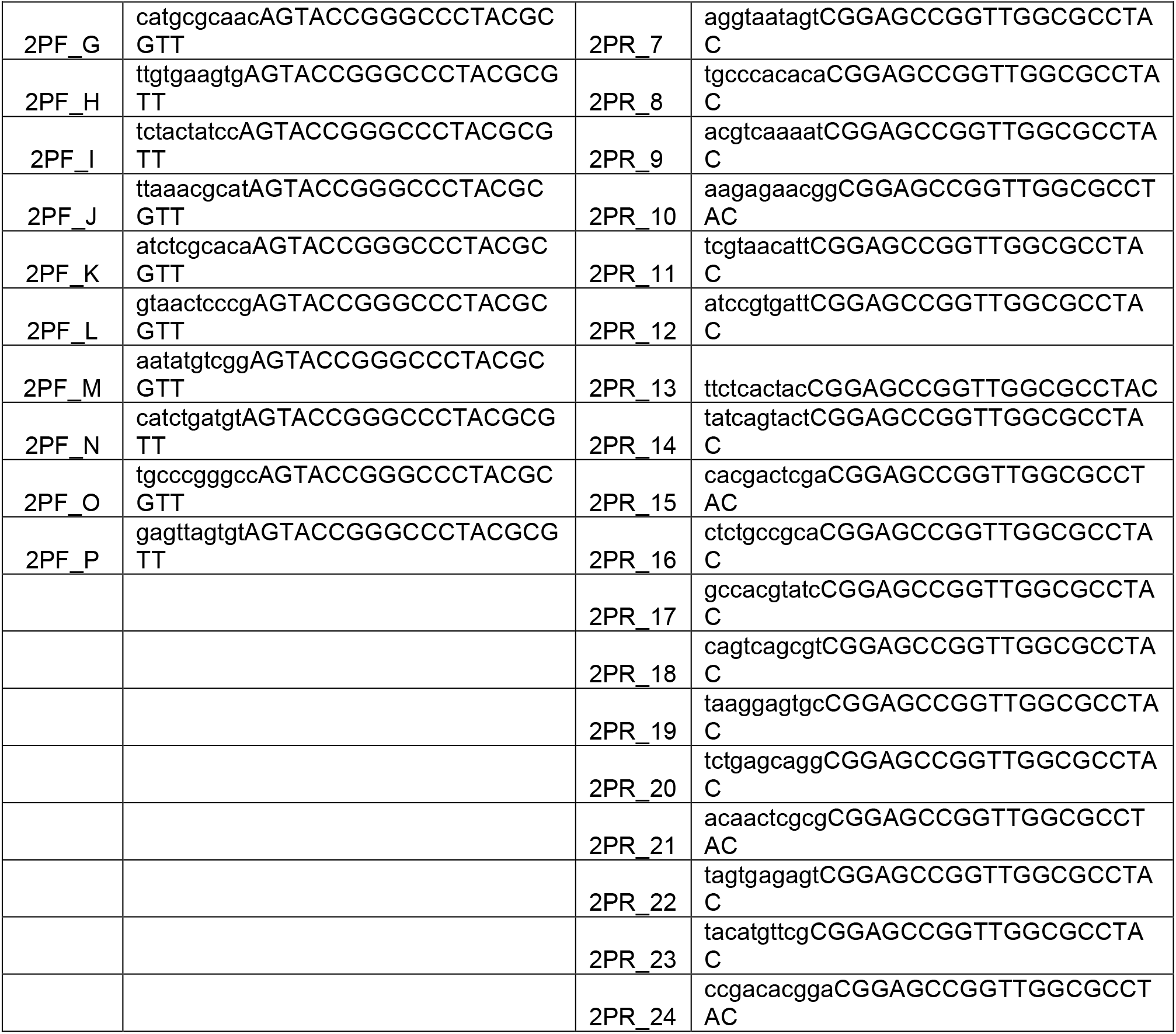

#### Processing of raw sequencing data

Consensus reads were generated from raw subreads with SMRT Link software (Pacific Biosystems) using standard settings. Barcode demultiplexing of the consensus reads was performed using SMRT Link software, version 9.0.0.92188, with two consecutive calls to the “lima” tool for plate and well barcodes, respectively. Only consensus reads that met the following criteria were retained: At least 80% of both barcode sequences must be present, and both barcodes must have a quality score of 25 or higher. This was done by specifying the following non-default parameters to the SMRT Link “lima” tool: “--min-score 25 --min-ref-span 0.8 --min-scoring-regions 2”. For plate barcodes, the “--same” parameter was given to indicate that the same barcode was present on either side of the read, whereas the “--different” parameter was given for well barcodes, which were flanked by one row and one column barcode. The data file containing the sequences and metadata for all consensus reads was converted from .bam to .sam format using SAMtools, version 1.9.

#### Filtering for read and barcode quality

Additional filtering for read and barcode quality was performed using custom R scripts. For each consensus read, the number of full passes (number of complete subreads) and the estimated read quality were extracted from the metadata of the .sam file and integrated with data from the two “.lima.report” files generated by demultiplexing the plate and well barcodes. In order to avoid incorrect assignments to plates and wells, all reads had to meet the following additional criteria: For both plate and well barcodes, the “ScoreCombined” and “ScoreLead” values from the “.lima.report” file had to reach a minimum of 60 and 30, respectively.

For the 10-nucleotide well barcodes, we imposed the additional restriction that either the correct barcode sequence must be present in full, or the barcode sequence must be at least eight nucleotides in length and flanked by an entirely correct 20-nucleotide constant region. We further filtered by read quality by applying the following minimal standards: seven full passes, a read quality of 0.9999, and a mean per-base Phred Quality Score of 85 (out of a maximum achievable score of 93). For 99.9% of plasmids in our libraries, at least one high-quality consensus read fulfilled these criteria. If there were no consensus reads for a specific well, we relaxed the requirements to include reads with at least three full passes and a read quality of 0.99 or higher; these criteria were used for 15 plasmids. Only 23 plasmids in our libraries (0.05%) were not represented by any reads.

#### Analysis of consensus reads

To quantify the percentage of correct guide RNA sequences and to identify well-to-well contaminations, each read was searched for the combined sgRNA and tracrRNA sequences in the forward and reverse directions, and all perfect matches were counted. To further characterise incorrect sequences, each consensus read was aligned to the correct barcoded reference sequence for that well. Alignment was done with the “pairwiseAlignment” function of the Biostrings package from the Bioconductor project, version 2.54.0, with a gap opening penalty of 30. Once all reads were aligned to a unified set of coordinates, the sequences corresponding to each combined sgRNA and tracrRNA region were extracted. Each region was classified as a) entirely correct, b) a contamination (if it was a perfect match for a sgRNA sequence from another plasmid), c) a large deletion (if more than 50% of the aligned sequence was composed of gaps), or d) a mutation (all other alterations). For each well, aggregate statistics were computed on the number of correct sgRNA modules, cross-well and cross-plate contaminations, and the number of deletions affecting each element of the 4sgRNA expression cassette. These metrics were displayed in heatmaps and two-dimensional bar plots and used to identify plates with quality issues.

### Benchmarking 4sgRNA ablation plasmids in cells and organoids

#### Cell culture

HCT116-Cas9 cells were grown in DMEM medium (GIBCO) supplemented with 10% FBS (GIBCO) and 1× penicillin/streptomycin (GIBCO) in a humidified incubator at 37°C with 5% CO_2_. For passaging, HCT116-Cas9 cells were washed once with D-PBS (GIBCO) and detached using 0.25% Trypsin (GIBCO). iPSC-iCas9 cells were cultured in mTeSR (Stem Cell Technologies) supplemented with penicillin/streptomycin (GIBCO) and doxycycline (200 ng/ml; Clontech) on laminin-521 (Biolamina) coated plates at 37 °C and 5% CO_2_. For routine maintenance, approx. 70% confluent cultures were dissociated into single cells with TrypLE (GIBCO) and seeded at a seeding density of 10,000–25,000 cells/cm^2^ in mTeSR supplemented with 2 μM ROCK inhibitor Y27632 (Tocris) onto laminin-521 coated plates. After 24h, the medium was replaced with mTeSR without ROCKi followed by daily medium changes. Kidney organoid differentiation and maintenance were performed as described (Ungricht et al., 2022). HEK293T cells were cultured in DMEM medium (GIBCO) supplemented with 10% FBS (GIBCO) in a humidified incubator at 37 °C with 5% CO_2_.

#### Transduction

Each batch of virus was titrated by transduction of 1.5 × 10^5^ HCT116-Cas9 or iPSC- iCas9 cells per well in a 6-well plate or 2.5 × 10^3^ NPC-iCas9 cells per well in an ultra-low attachment (ULA) 384**-**well plate with different dilutions of the virus in each well. For each viral concentration and the no-virus control, three to four replicate wells were seeded. After 24 hours, medium containing 2 µg/ml puromycin and 8 µg/ml polybrene (HCT116-Cas9) and 1 µg/ml (iPSC-iCas9) or no (NPCs- iCas9) puromycin (GIBCO) was exchanged. A no-virus control was always included and untransduced control cells did not survive 72 hours of puromycin treatment. After puromycin selection, live and dead cells were counted. The viral titer of each batch was identified by calculating the percentage of puromycin surviving cells relative to the no-virus control. For NPCs, all viral volumes were functionally read out via FACS analysis. For all subsequent transductions, the viral volume was calculated to reach an MOI of 5. Cells were cultured as described above and knockout was measured 4 or 8 days (HCT116-Cas9 or iPSC-iCas9) or >14 days (NPC-iCas9) after transduction.

#### Transfection

HCT116-Cas9 or iPSC-iCas9 cells were seeded so that cells reached 60–80% confluency 24 hours post-seeding. On the day of transfection, growth medium was exchanged for medium without penicillin or streptomycin. 4sgRNA plasmids or synthetic guides complexed with the tracrRNA following the manufacturer’s protocol (IDT) as described below were diluted (5 µg of plasmid or 10 µM tracr-complexed synthetic guide RNAs) in OptiMEM (GIBCO). Lipofectamine 2000 (Invitrogen) was diluted in OptiMEM according to manufacturer’s protocol and incubated for 5 minutes at RT. DNA or RNA and diluted Lipofectamine 2000 were mixed dropwise and incubated for 20 minutes at RT. The DNA/Lipofectamine 2000 mixture was added gently to the cells. Growth medium was exchanged 24h post-transfection. Cells were cultured as described above and knockout was measured 4 or 8 days after transfection.

#### Nucleofection

2 × 10^5^ HCT116-Cas9 and iPSC-iCas9 were resuspended in 20 µl SE cell line nucleofection solution (Lonza) (HCT116-Cas9) or P3 primary cell nucleofection solution (Lonza) (iPSC-iCas9). Cells were mixed and incubated at room temperature for 2 min in PCR tubes. 5 µg of the 4sgRNA plasmid or 10 µM synthetic guide RNAs were mixed and the cell/reagent/nucleofection mix was transferred to Nucleofection cuvette strips (Lonza). Cells were electroporated using a 4D nucleofector (4D-Nucleofector Core Unit: Lonza, AAF-1002B; 4D-Nucleofector X Unit: AAF-1002X; Lonza). Programs were adapted for the different cell types (HCT116-Cas9: EN-113, iPSC-iCas9: CD-118). After nucleofection, prewarmed cell-specific growth media was used to transfer transfected cells in culture plates containing pre-warmed cell-specific growth media. Cells were cultured as described above and knockout was measured 4 or 8 days post-nucleofection.

#### Live immunostaining and FACS analysis

Cells were harvested and resuspended at a concentration of 1 × 10^6^ cells/100µl in FACS buffer (1× PBS (GIBCO), 5 mM EDTA (Sigma) and 1% FBS (GIBCO)). Afterwards, per 1 × 10^6^ cells, 1 µl of Alexa488 anti-human EPCAM (Abcam, ab112067) or Alexa647 anti-mouse/human CD44 (Biolegend, 103018) was added. After incubation for 10–20 minutes at RT, cells were washed 2× with 2 ml FACS buffer. Afterwards, cells were resuspended in 250 µl FACS buffer and analyzed with a Fortessa (BD) or Canto (BD) analyzer.

#### Preparation of crRNA–tracrRNA duplex and precomplexing of Cas9/RNP

To prepare the duplex, each Alt-R crRNA and Alt-R tracrRNA (IDT) was reconstituted to 200 µM with Nuclease-Free Duplex Buffer (IDT). Oligos were mixed at equimolar concentrations in a sterile PCR tube (e.g., 10 µl Alt-R crRNA and 10 µl Alt-R tracrRNA). Oligos were annealed by heating at 95 °C for 5 min in a PCR thermocycler and the mix was slowly cooled to room temperature.

#### The FLAER assay

NPCs were transduced with lentiviruses carrying the 4sgRNA plasmid targeting the gene *PIGA* or with a pool of four individual lentiviruses each carrying a sgRNA targeting the gene *PIGA* as described above. After 46 days post-transduction, organoids were dissociated into single cells and stained with FLAER-488 reagent (Biozol) in 3% BSA (blocking solution) according to the manufacturer’s protocol. Subsequently, the percentage of FLAER-negative cells in each condition were analyzed using a Fortessa FACS analyzer (BD).

### Arrayed activation screen for PrP^C^ regulating transcription factors

#### Cell culture

The U-251 MG human cells (Kerafast, AccessionID: CVCL_0021) stably expressing dCas9-VPR were cultured in T150 tissue culture flasks (TPP) in OptiMEM without Phenol (Gibco) supplemented with 10% FBS (Takara), 1% NEAA (Gibco), 1% GlutaMax (Gibco), 1% Penicillin/Streptomycin (P/S) (Gibco) and 10 µg/mL of blasticidin (Gibco). Once the cells reach a confluency of 80–90%, they were harvested with Accutase (Gibco), washed with PBS and resuspended in medium, pooled, and counted using TC20 (BioRad) Cell Counter with trypan blue (Gibco).

#### PrP^C^ screening

The screen was performed as previously described (Heinzer et al., 2021). 5’000 cells (per well) of U-251 MG dCas9-VPR were seeded in 30 µl of medium into white 384-well cell culture plates (Greiner Bio-One, #781080). The plates were incubated in a rotating tower incubator (LiCONiC StoreX STX, Schaanwald, Liechtenstein). 24 hours later, cells were transduced with lentiviruses containing the 4sgRNA vector targeting each TF at an MOI of 3. Each plate contained 14 wells with non-targeting (NT) and another 14 wells with *PRNP* targeting controls. Experiments were performed in triplicates. Plates were further incubated for four days. On the day of the assay, one replicate was used to determine cell viability using CellTiter-Glo® (Promega) according to the manuals. The luminescence was measured with the EnVision plate reader (Perkin Elmer).

The other two replica plates were used to assess PrP^C^ levels by the TR-FRET method (Heinzer et al., 2021). Cell culture medium was removed by inverting the plates, and cells were lysed in 10 μL of lysis buffer (0.5% Na-Deoxycholate (Sigma Aldrich St. Louis, MO, USA), 0.5% Triton X (Sigma Aldrich), supplemented with EDTA-free cOmplete Mini Protease Inhibitors (Roche, Basel, Switzerland) and 0.5% BSA (Merck, Darmstadt, Germany). Following lysis, plates were incubated on a plate shaker (Eppendorf ThermoMixer Comfort) for 10 min (4° C, 400 rpm shaking conditions) prior to centrifugation at 1000×g for 1 minute and incubated at 4° C for two additional hours. Following incubation, plates were centrifuged once more under the same conditions mentioned above and 5 μL of each FRET antibody pair was added (2.5 nM final concentration for the donor and 5 nM for the acceptor, diluted in 1× Lance buffer (Perkin Elmer)). POM1 (binding to amino acid residue a.a 144– 152) and POM2 (binding to a.a 43–92) (Polymenidou et al., 2008), targeting different epitopes of PrP^C^, were coupled to a FRET donor, Europium (EU) and a FRET acceptor, Allophycocyanin (APC), respectively, following previously reported protocols (Ballmer et al., 2017) . Plates were centrifuged once more and incubated overnight at 4° C. Then, TR-FRET measurements were read out using previously reported parameters (Ballmer et al., 2017) on an EnVision multimode plate reader (Perkin Elmer).

#### Data analysis

The first and last columns of each 384-well plate were reserved for blanks (wells containing only one of the antibodies, or buffer only), which were used to calculate net FRET values and for background subtraction as previously described (Ballmer et al., 2017).

For plate-wise quality control, the separation between positive and negative controls was assessed using the Z′-factor 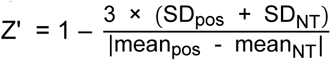, where SD_pos_ and SD_NT_ denote the standard deviation of positive and non-targeting controls, respectively (Zhang et al., 1999).

To obtain logarithmised and normalised PrP^C^ expression values (log_2_ fold change values), raw values were log_2_ transformed and normalised by subtracting the median expression value of all genes on that plate. Mixed-moment estimates of sample SSMD values were computed using the formula 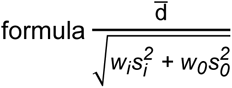, and *t* values were calculated as 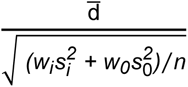 (<u>Zhang, 2011</u>). In our case, d̅ was the mean of normalised expression values of the two replicates for each plasmid. The weights (w_i_ and w_0_) were both set to 0.5; the variable s_i_^2^ referred to the variance of the two replicates for each plasmid, and s_0_ ^2^ was the median of all variances for pairs of non-targeting control wells on replicate plates. Two-sided *p* values were derived from sample *t* values using the Student’s *t*-distribution with one degree of freedom.

#### Hits validation

The 36 primary hits that passed the cutoff criteria were further repeated with the same FRET-based assay in 384-well plates with 5 technical repeats. Then the respective top 10 hits of PrP^C^ upregulators and downregulators were individually confirmed in 6-well plates in U-251MG dCas9-VPR cells with their corresponding 4sgRNA lentiviruses four days post-viral transduction and 1 µg/ml puromycin selection via western blotting. The POM2 primary antibody against PrP^C^ was used for the assay. Vinculin was used as a loading control. The levels of PrP^C^ were quantified by ImageJ and normalized to Vinculin.

### Pooled genome-wide CRISPR knockout autophagy screens

#### Cell culture

H4 (ATCC, HTB-148) cells were kept in a humidified incubator at 37 °C and 5% CO_2_ and were maintained in DMEM + 10% FBS, 1% L-glutamine and 1% Penicillin/Streptomycin. Cell culture reagents were obtained from Invitrogen. H4-Cas9-GFP-SQSTM1 and H4-Cas9 cells were generated in the previous study (DeJesus et al., 2016).

#### Pooled CRISPR screens

sgRNA plasmids of pooled T.spiezzo, an optimized Brunello library that covers 18’360 genes (5 sgRNAs per gene whenever possible, split into two sgRNA sub-pools named CHIP1 to CHIP6) (Estoppey et al., 2017) and Cellecta pooled knockout libraries were amplified and packaged into lentiviral particles using HEK293T cells. Packaging was scaled up by growing cells in CellSTACK flasks (Corning #3313). For each cell stack, 2.1 × 10^7^ cells were transfected 24h after plating using 510.3 µl of TransIT reagent (#MIR2300, Mirus) diluted in 18.4 ml of OPTI-MEM that was combined with 75.6 µg of each sgRNA libraries and 94.5 µg of lentiviral packaging mix (#CPCP- K2A, Cellecta; containing psPAX2 and pMD2 plasmids that encode Gag/Pol and VSV-G, respectively). 72h post-transfection, lentivirus was harvested, aliquoted, and frozen at −80 °C. Viral titers of the sgRNA libraries were determined by a 6-point dose response in 6-well plates by puromycin selection and determination of live and dead cells (T.spiezzo, Cellecta) or flow cytometry of RFP+ cells (Brunello) using H4-Cas9-GFP-SQSTM1 cells 3 days after infection.

H4-Cas9-GFP-SQSTM1 cells were infected at an MOI of 0.4 with at least 300 coverage of each sgRNA at any time during the screen. This resulted in 50–60 million H4-Cas9-GFP-SQSTM1 in CellSTACK flasks (#3313, Corning) that were infected with the corresponding volume of lentiviral supernatant. 24h post-infection, cells were selected with puromycin (2 μg/ml, GIBCO) for 3 days and then maintained in culture and split as needed to ensure confluence did not exceed 90% for a further 7 days. Then cells were harvested and resuspended at a density of 30 million cells/mL, and live and single cells were sorted (BD ARIA III) from the lower GFP quartile (GFP^low^) or from the upper GFP quartile (GFP^high^) and subsequently fixed in 4% PFA in PBS. For each screen, 20–25 million cells were isolated by FACS in the GFP^low^ and GFP^high^ category. 25 million unsorted cells were also collected as an input sample. For all samples, genomic DNA (gDNA) was isolated using phenol chloroform extraction. In short, cells were resuspended in 5 mL TNES (10 mM Tris-Cl pH 8.0, 100 mM NaCl, 1 mM EDTA,1% SDS) and incubated overnight at 65 °C to reverse PFA crosslinks. After allowing the samples to cool, samples were incubated with 100 μl RNase A (QIAGEN) for 30 min at 37°C, followed by addition of 100 μl of proteinase K (QIAGEN) for 1 h at 45 °C. 5 mL PCIA (Phenol:Cholorform:Isoamyl alcohol pH8) (Thermo Fisher) were added and samples vortexed, spun at max speed for 2 min, and the aqueous phase was transferred to 5 mL of PCIA. Samples were vortexed again and spun at max speed for 2 min. The aqueous phase was transferred to 4.5 mL of chloroform. A third time, samples were vortexed, spun at max speed for 2 min, and the aqueous phase was transferred to 400 μl 3M NaAcetate pH 5.2. Later, 10 mL of 100% EtOH was added, samples were mixed and DNA precipitated for 1 h on ice, followed by spinning at max speed for 10 min. EtOH was decanted and pellets washed with 10 mL 70% EtOH. Finally, samples were spun at max speed for 10 min, pellets air-dried and resuspended in 1 mL of nuclease free water.

#### Sequencing libraries preparation

For gDNAs of GFP^high^, GFP^low^ and unsorted samples from the T.spiezzo pooled screen, we developed an Illumina paired-end sequencing strategy. Briefly, sgRNA2 and sgRNA3 from the 4sgRNA library were amplified (595 bp) by PCR using NEB Q5 DNA polymerase (NEB, Ipswitch MA, USA) with a universal P7 primer and individual P5 primer, each with a unique index. We designed a custom sequencing primer (Read-1 primer) for read 1 detecting the sequence of sgRNA2 and designed a custom index sequencing primer (Index-1 sequencing primer) detecting the sequence of sgRNA3, and an index sequencing primer (Index-2 sequencing primer) for demultiplexing samples. See below the schematic of the sequencing strategy.

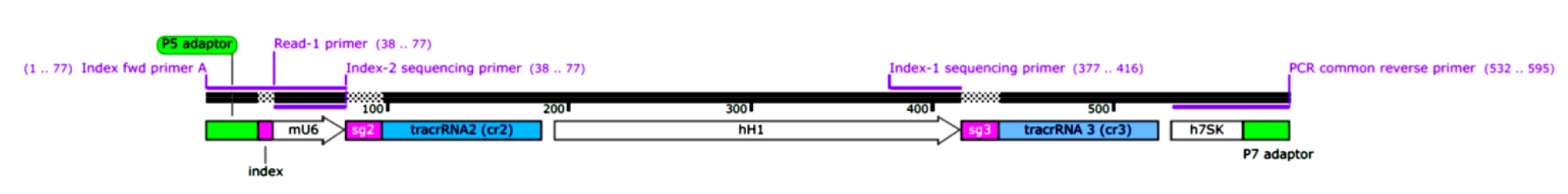

For each PCR, the amplification parameters are 98 °C for 30 seconds, 98 °C for 10 s, 55 °C for 30 s, 72 °C for 120 s, and 72 °C for 120 seconds. We used 30 to 60 µg of gDNA from each sample and performed 16 to 31 individual PCR reactions, each with 2 µg gDNA as the input, which were pooled and reamplified with the same primers.

The amplification and sequencing primers used are listed as follows:

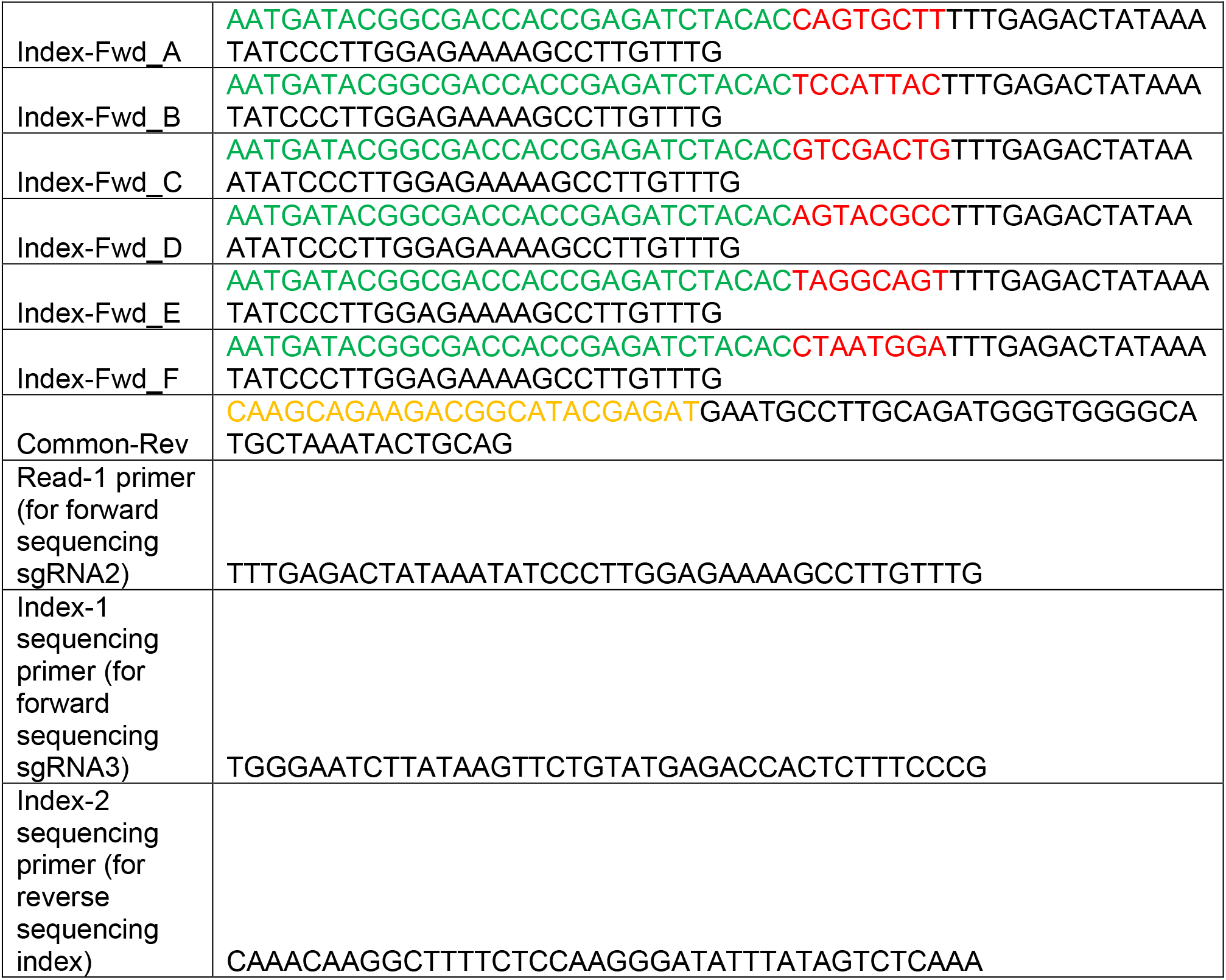

Green, red, and yellow colored nucleotides indicate P5 adapter, index, and P7 adapter sequences of the primers, respectively.

The sequencing library preparations for Brunello and Cellecta library screens were performed according to the published protocols (Doench et al., 2016; Sanson et al., 2018).

All library samples were sequenced on an Illumina NovaSeq 6000 using S1 flowcells and the XP workflow.

#### Pooled screen data analysis

All FASTQ files were aligned using bowtie2 with the following parameters: -L10 -N0 -iS,1,2; seed length is 10, zero mismatches allowed in seed alignment, interval between seeds is 1 + 2 * sqrt(read length) which is smaller than the default (-iS,1,2.5). This is adapted to the short sequences (20 nt) that are being aligned and provided better results that standard bowtie2 settings (which are for longer reads of around 50 nt). For T.spiezzo, two bowtie2 indices were built, one for sgRNA2 and one for sgRNA3. For the Brunello library two bowtie2 indices were built, one for CHIP1-3 and one for CHIP4-6 of the Brunello library. For the Cellecta library data (DeJesus et al., 2016), historical counts from the original screen were used, those were obtained with bowtie1.

All alignments were written to SAM files which were then further processed with samtools (statistical analysis, filtering, counting). Counts were obtained by counting the reference names, i.e. sgRNA annotations from SAM files. For T.spiezzo, each sample was processed to most accurately reflect the Illumina-based sequencing protocol that was employed: Read 1 of each sample should only be from the sgRNA2, i.e. the reference sequence should contain a sg2 in its name; reference sequences of the second read should contain a sg3 in their name; due to the alignments using two separate indices for the FASTQ files from read 1 and 2, all alignments from reads 1 and 2 must contain sgRNA2 and sgRNA3 sequences, respectively. The intersection of sgRNA2 first reads and sgRNA3 second reads provides the names of all valid reads across the FASTQ files for read 1 and read 2. FASTQ files are filtered to these common read names (using the -N option to samtools) and unaligned reads are discarded (-F4 option to samtools). Because of this filtering, the number of reads from the respective SAM files are the same. Additionally, the two corresponding reads are then checked for targeting the same gene, and discarded if this is not the case. The reported valid guide rate takes into account correct sequences and cohesiveness of both sgRNA sequences.

Fold changes were estimated using edgeR. These were obtained for the comparison of the GFP^high^ versus the GFP^low^ samples in all three screens used for analysis (T.spiezzo, Brunello, and Cellecta).

Enrichment analysis were performed using a thresholded approach (overrepresentation analysis, ORA). The top or bottom 200 genes with the highest or lowest fold change were used for ORA. Gene sets from the biological process branch of the Gene Ontology were used as reference sets. Enrichment calculations were done using the clusterProfiler package in R.

Boxplots for enriched gene sets and/or gene sets related to autophagy were prepared using the absolute fold-change. This was done in order to be able to quantify the effect of the screening library via analysis of variance modeling. In short, the absolute fold change was modeled as a linear function of gene and screening library (abs(logFC) ∼ gene + screen) R formula), and the significance of the screen coefficient was determined using ANOVA (aov in R).

#### Hits validation

Among the top hits uniquely identified by the T.spiezzo pooled library, 10 potential novel regulators together with 6 known autophagy regulators were selected for further validation. Lentiviruses of the corresponding 4sgRNA plasmids targeting the 16 genes or non-targeting controls (NT) were individually packaged, titrated, and transduced to H4-Cas9-GFP-SQSTM1 cells at an MOI of around 0.3 in 24-well plates. Then cells were split and selected in 2 μg/ml puromycin medium for 3 days and further cultured in normal medium without antibiotic selection for a week in duplicates. One replicate of cells was analyzed by flow cytometry to quantify the log_2_ fold change of cell numbers in GFP^high^ vs GFP^low^ populations, as in the primary pooled screen. The other replicate of cells was imaged under a fluorescent microscope to analyze the distribution of GFP-SQSTM1 puncta. NT-treated cells were used as controls for both conditions. The area of GFP-SQSTM1 puncta was determined with CellProfiler, and the percentage of cells with GFP-SQSTM1 puncta was analyzed with ImageJ (NIH).

To further confirm the potential novel regulators of autophagy, H4-Cas9 cells were individually transduced with 4sgRNA lentiviruses against these genes or NT, and cultured the same as the GFP- SQSTM1 assay in 10-cm dishes in duplicates. 6 hours before collection of samples, one replicate of cells was treated with vehicle (medium) and one replicate was treated with a final concentration of 100 µM chloroquine (ChQ). At the end point, cells were washed twice with cold PBS before adding ice cold 1× RIPA buffer (Cell Signaling) with protease (Complete, Roche) and phosphatase inhibitors (PhosSTOP, Roche) directly to the well. After cell scraping, the suspension was collected in a tube and incubated on ice for 30 minutes before centrifugation at maximum speed for 10 minutes. Supernatant containing cell lysates was collected in a new tube and the protein concentration assessed through BCA assay (Pierce). 5 μg of total protein was separated in a 12% Bis-Tris gel with MOPS running buffer and subsequently transferred to a PVDF membrane. The following antibodies were used for immunoblotting: LC3b (CellSignaling #3868) and GAPDH (CellSignaling #2118). The experiments were repeated twice. The intensities of LC3-II were quantified with ImageJ and normalized to GAPDH intensities.

For the YFP-LC3 reporter assay, H4-Cas9 cells were cultured following a similar protocol as used for the GFP-SQSTM1 assay. 48–60 hours before the experiment, 2’000 gene-edited or NT-treated H4-Cas9 cells were seeded into 8-well chamber and 6 hours later (or the next day), they were transduced with YFP-LC3 lentiviruses at an MOI of 3. Two days later, each well of cells was treated with vehicle or 100 µM of ChQ for 6 hours, before cells were imaged with Leica LAS X confocal imaging system. The puncta area of YFP-LC3 was analyzed in a similar manner to that of the GFP- SQSTM1 experiments.

### CRISPRoff expriments

#### CRISPRoff tests with individual 4sgRNA plasmids

Individual 4sgRNA plasmids for CRISPRoff were assessed in HEK293T cells via plasmid transient transfection. A pool of three single sgRNAs (150 ng) or individual 4sgRNA plasmids (150 ng) were co-transfected with the CRISPRoff (300 ng) or CRISPRoff-D3A mutant (300 ng) plasmids into HEK293T cells in 24-well plates with Lipofectamine 3000 once the cells reached 80–90% confluency. The second day, cells were split into 1 µg/ml puromycin containing medium and selected for three days to eliminate non-transfected cells. Then cells were maintained in normal growth medium without antibiotic selection for one week. Cells were trypsinized and single live-cell solutions were stained using the same method as described for the gene ablation efficiency assay with the corresponding fluorophore-conjugated primary antibodies and analyzed with flow cytometry. Gene ablation efficiencies of 4sgRNA plasmids from the T.spiezzo library were determined as described above. Non-targeting sgRNA-treated cells were used as controls.

#### Genome-wide CRISPRoff screens

The T.gonfio pooled library was amplified with Lucigen E. cloni 10G Elite electrocompetent cells through electroporation. Briefly, 150 ng plasmid of the library was added to 200 μl competent cells, followed by 4 aliquots with 50 μl loaded into each electroporation cuvette (0.1 cm gap) for further electroporation using Gene Pulser Electroporation System (voltage in 1600 V, capacitance in 25 μF, and resistance in 200 Ω). Afterwards electroporations were mixed with chilled recovery medium, followed by shaking at 250 rpm for 1 hour at 37 ℃. Then all transformations were pooled and distributed in 4 pre-warmed 24.5 cm^2^ bioassay plates (trimethoprim resistance) and grown at 30 ℃ for 16 hours. Finally, bacteria in plates were harvested for plasmid extraction with QIAGEN EndoFree Plasmid Maxi Kit (Catalog # 12362). Finally, amplified sgRNA libraries were packaged into lentiviruses with HEK293T cells followed by titration with flow cytometry analysis as described above. For the dual-sgRNA CRISPRoff library, DH5α competent cells (3 × 10^8^ cfu/μg in 75 μl) were used to transform 100 ng plasmids to amplify the library as previously described (Nuñez et al, 2021).

The CRISPRoff genome-wide dropout screen was performed as described (Nuñez et al, 2021). Briefly, 60 million HEK293T cells for each library were seeded in 2 × T300 culturing flasks, followed by virus transduction at an MOI 0.3 with a coverage of about 1000 cells per plasmid 6 hours post-seeding. 2 days later, the cells were passaged with 1:3 ratio and maintained for 4 days in the presence of puromycin, during which the percentage of GFP-positive (for dual-sgRNA library) or BFP-positive (for T.gonfio 4-sgRNA library) cells were monitored with flow cytometry analysis until a proportion of 90% was achieved. Next, 60 million cells were seeded in 2 × T300 flasks without puromycin. About 24 hours later, cells with a confluency of about 80% (T0 time point) in each flask were transfected with 57 µg of the CRISPRoff-mScarletI plasmid using Lipofectamine 3000. On the second day, transfected cells were passaged at a 1:2 ratio for cell sorting. One day later, 27 million cells double-positive for mScarletI-GFP (for the CRISPRoff library screen) or mScarletI-TagBFP (for the T.gonfio library screen) were sorted for each library, seeded in T300 flasks and maintained for 10–12 cell passages (T10 time point). 60 million cells for each data point were collected for gDNA extraction and subsequent sequencing analyses.

PCR amplicons of the screen with the CRISPRoff library were prepared according to established methods (Nunez et al., 2021). PCR amplicons of the T.gonfio library screen were prepared the same as the pooled screen for autophagy using the T.spiezzo pooled library. For each sample of the CRISPRoff screens, 130 µg of gDNA was used in 65 × 50 µl PCR reactions (2 µg gDNA per 50 µl PCR reaction), and all PCR products from the same sample were pooled and purified. All PCR amplicons from the two libraries were sequenced on a Illumina NextSeq500 device at the Functional Genomics Center Zurich.

#### Data analysis

First, sequences were mapped to sgRNAs from the reference CRISPR libraries. If no perfect match could be found, we relaxed the search criteria to include sgRNAs with a maximum edit distance of 1 (a single-nucleotide mismatch, insertion, or deletion was tolerated). Finally, mapped sgRNA sequences were allocated to plasmids. For a few genes in the T.gonfio and dual-sgRNA CRISPRoff libraries, the same sgRNAs were included for multiple plasmids. The inclusion of unspecific sgRNAs is sometimes inevitable when targeting genes with closely related paralogues. In such cases, the information from neighboring mapped sgRNA sequences was used for disambiguation, whenever possible. Sometimes, however, sgRNAs had to be assigned to multiple plasmids. When computing plasmid counts, these were weighted by the number of possible plasmids, resulting in fractional counts. This was separate from the handling of reads with template switches, which were excluded from further analysis.

Normalisation of plasmid counts: Only reads with two mapped sgRNAs without a template switch were included. For each screen, raw read counts were adjusted to account for any differences in the number of reads per sample using the median ratio method (Li et al., 2014). Initially, the geometric mean across samples was calculated for each plasmid. Next, counts were divided by the geometric mean across samples to produce a matrix of count ratios. A sample-specific size factor was computed, defined as the median of count ratios across all plasmids. Finally, raw counts were divided by the sample-specific size factor to produce normalized counts. Because the geometric mean is defined as the antilog of the sum of log values divided by the number of samples, and the logarithm of zero is undefined, only plasmids with non-zero counts for all samples were used for estimating the size factor. Sample size factors ranged from 0.81 to 1.21.

Comparison of genome-wide CRISPRoff screens: Both the T.gonfio and dual-sgRNA CRISPRoff libraries contained multiple plasmids for genes with major alternative transcription start sites (TSSs). To ensure a fair comparison between libraries, only one plasmid per gene was included, and guide RNAs targeting the principal TSS were preferred. The primary TSS was defined by the activity score from the FANTOM5 project (Abugessaisa et al., 2017). The sgRNA sequences of the dual-sgRNA CRISPRoff library were aligned to the hg38 reference genome and re-annotated with additional information, including Entrez gene identifiers and their location relative to the TSS. To define the gene sets used for the comparison, we downloaded the “common_essentials.csv” and “nonessentials.csv” data files from the Public 20Q2 release of the Cancer Dependency Map (DepMap, https://depmap.org/portal/download/all/).

The robust estimate of the strictly standardised mean difference (SSMD) was computed as a measure of the separation between essential and non-essential genes (Zhang, 2007, 2011) as 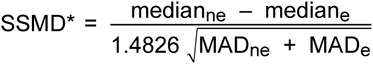, where MAD_ne_ and MAD_e_ denote the median absolute deviation of non-essential and essential genes, respectively.

## Supplementary Figures

**Figure S1.**
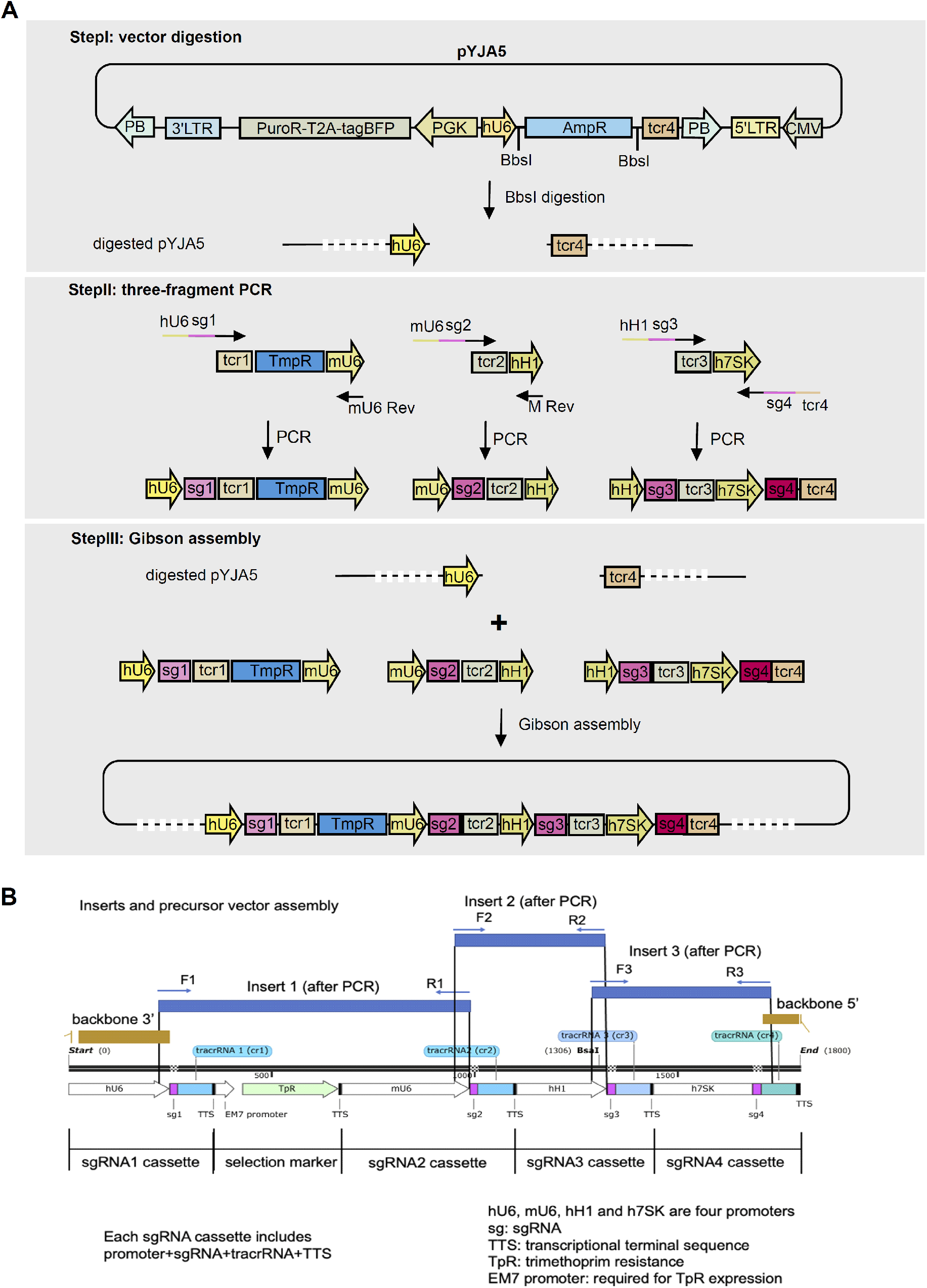
Details of APPEAL cloning method. **A**, Step-by-step construction of 4sgRNA plasmids using APPEAL cloning method. **B**, Zoom-in illustration of homologous ends overlapping among the three amplicons and the digested vector pYJA5.

**Figure S2.**
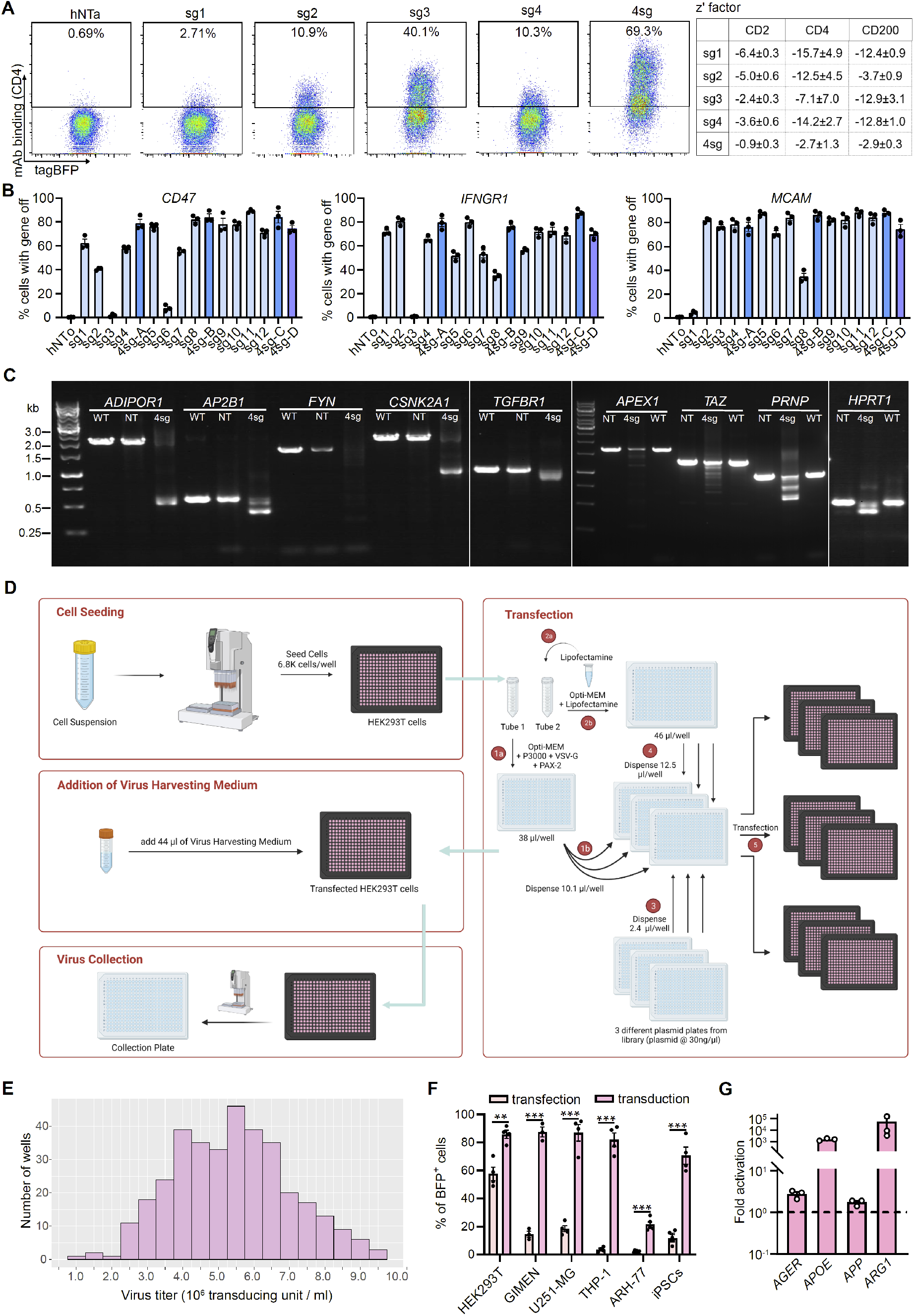
High efficiency of 4sgRNAs in gene activation/ablation and high-throughput lentiviral production. **A**, left, Surface expression of CD4 in HEK293 cells expressing dCas9-VPR and single sgRNA (sg1-4) or 4sgRNA (4sg). hNTa, non-targeting control plasmid; right, z’ factor of single sgRNAs-or 4sgRNAs-activated conditions compared with their corresponding hNTa-treated conditions. **B**, Gene disruption efficiency by single sgRNAs vs 4sgRNAs in HEK293 cells via co-transfection with Cas9 plasmid. 12 single sgRNAs (sg1-12) from the Brunello, GeCKOv2 and TKOv3 libraries (light blue) were tested, 4sg-A, B and C (dark blue) and 4sg-D (purple bule) were 4sgRNA plasmids assembled with sg1-4, sg5-8, sg9-12 or the 4 worst-performed single sgRNAs among the 12 sgRNAs tested; hNTo, non-targeting control plasmid. **C**, An example of gel images of PCR amplicons amplified from 4sgRNA (4sg) knockout plasmid-edited genomic DNAs. For each of the genes tested, a pair of primers flanking all the 4sgRNA targeting sites was designed (see details in materials and methods). PCR amplicons of *ADIPOR1*, *AP2B1*, *CSNK2A1*, *FYN*, *HPRT1*, *TGFBR1*, *APEX1*, *TAZ*, and *PRNP* conditions amplified from wildtype (WT) or non-targeting (NT) plasmid treated cells showed expected size of 2095, 558, 2225, 1975, 514, 987, 1663, 1288, and 959 bp on agarose gels, whereas for amplicons from 4sg-edited cells in each condition a great majority of them showed shorter sizes, indicating DNA deletions in the corresponding edited genes by 4sg knockout plasmids. **D**, A schematic of lentivirus production in 384-well plates (see details in materials and methods). **E**, Titers of lentiviral particles packaged from an example of 384-well plate of library plasmids. Viral particles were produced in 384-well plates with HEK293T cells, and transduced to further HEK293T cells. TagBFP+ cells were quantified by flow cytometry 3 days post transduction. **F**, Gene delivery efficiency of 4sgRNA vector into poorly transfectable cells, measured by flow cytometry of TagBFP^+^ cells 3 days post transduction. **G**, Gene activation in neurons derived from human induced pluripotent stem cells, measured by qRT-PCR after transduction of 4sgRNA lentiviruses (multiplicity of infection: 1.4). Target neurons expressed stably dCas9-VPR. Assays were performed at day 7 post infection.

**Figure S3.**
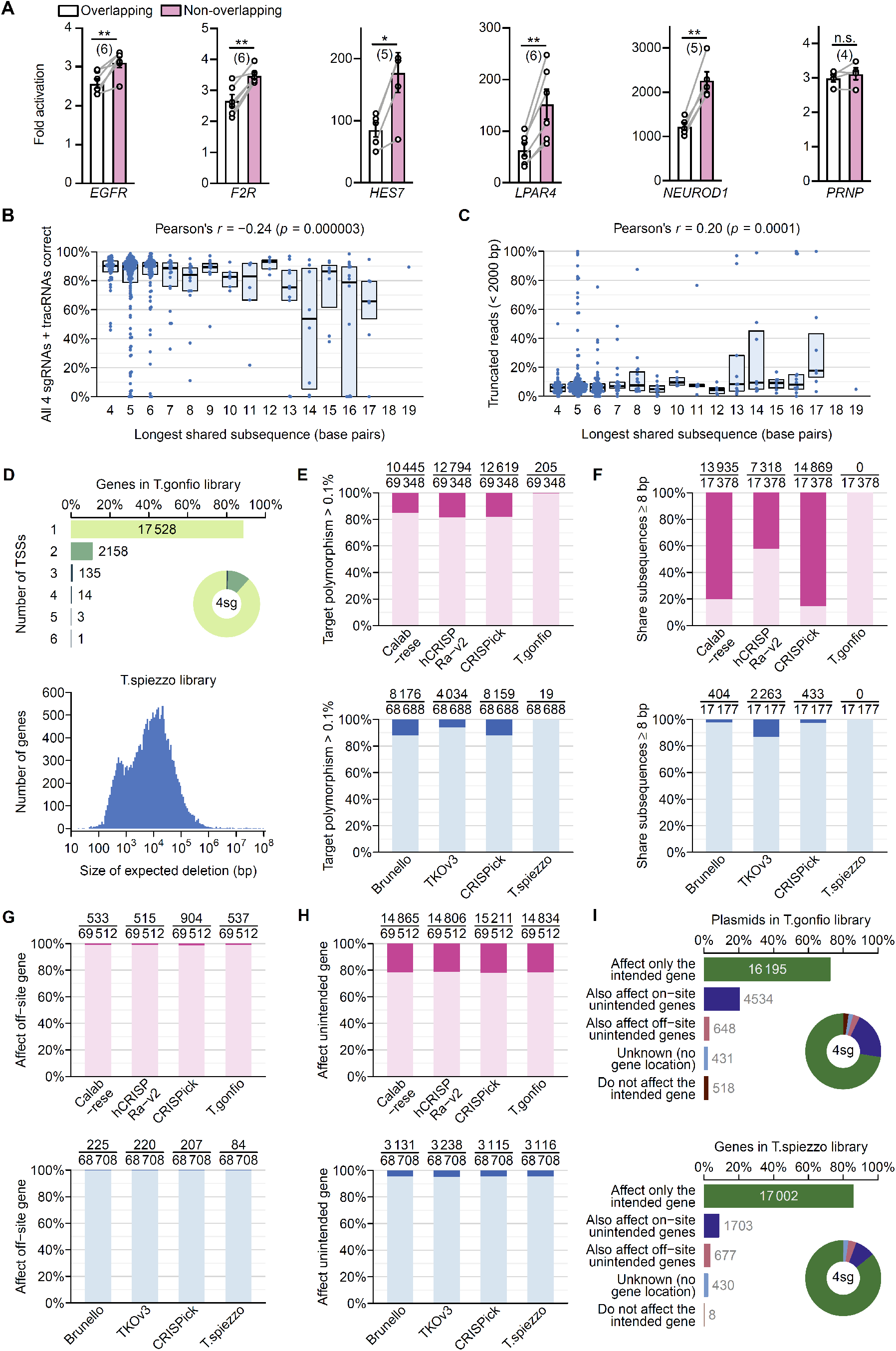
The effect of sgRNA spacing and homology on 4sgRNA plasmids and other features of T.spiezzo and T.gonfio libraries. **A**, Comparison of the effect of overlapping and non-overlapping sgRNAs on gene activation in HEK293 cells. **B**, Correlation between the extent of homology among the 4 sgRNAs and the percentage of correct plasmids. **C**, Correlation between the extent of homology and the frequency of shortened amplicon regions (indicating deletions). **D**, Summary of the number of transcription start sites (TSSs) per gene that are each targeted by a separate plasmid in the T.gonfio library (top), and the estimated size of deletions between the first and last cut sites of each 4 sgRNA plasmid in the T.spiezzo library (bottom). **E**, Percentage of sgRNAs that target genomic site affected by a polymorphism with frequency higher than 0.1% in the T.spiezzo and T.gonfio libraries in comparison with the top 4 sgRNAs from existing resources. **F**, Percentage of sgRNAs that share 8 or more base pairs of homology in the T.spiezzo and T.gonfio libraries in comparison with the top 4 sgRNAs from existing resources. **G** and **H**, Comparison of the percentage of sgRNAs predicted to target unintended genes at off-site locations (**G**) and all locations (**H**) – the latter include mostly sgRNAs with *on-site* unintended targets. **I**, All plasmids in the T.spiezzo and T.gonfio libraries were assigned to mutually exclusive categories, based on whether any of the 4 sgRNAs may target additional, unintended genes.

**Figure S4.**
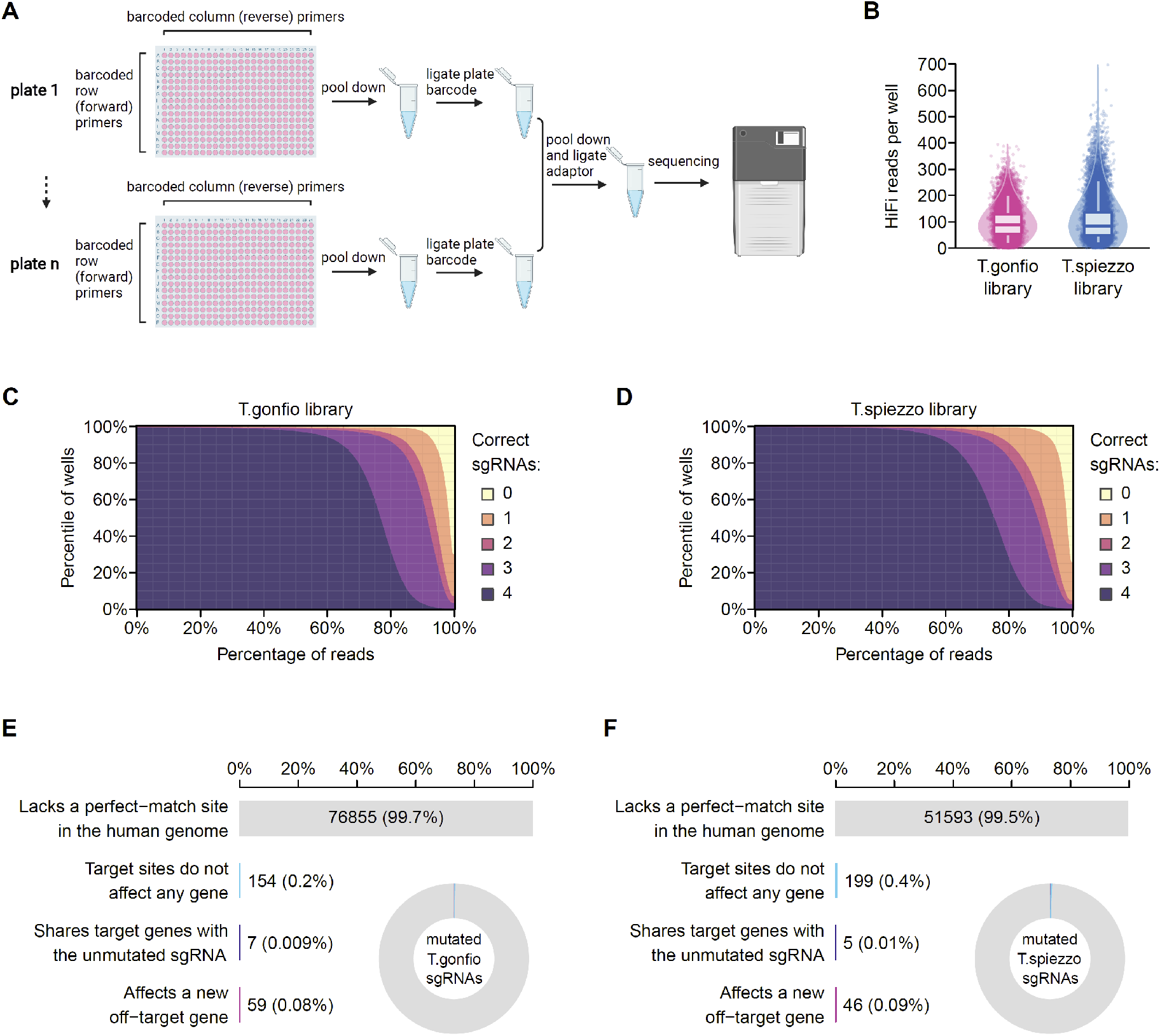
Genome-wide sequencing of the T.spiezzo and T.gonfio libraries. **A,** PacBio long-read sequencing workflow: polymerase chain reaction (PCR) was performed in each well of a 384- well plate using primers appended with row- and column-specific barcodes. All wells from one plate were pooled and ligated with plate-specific barcodes, and multiple plates were further pooled for sequencing. **B**, High-quality read count for each well in the T.spiezzo and T.gonfio libraries. **C** and **D**, Cumulative distribution of each well of plasmids with 0, 1, 2, 3, 4 entirely correct sgRNA and tracrRNA sequences, as well as an associated promoter sequence that was at least 95% correct, in the T.spiezzo and T.gonfio libraries. **E** and **F**, Predicted off-target effects for mutated sgRNAs in the T.spiezzo and T.gonfio libraries. Guide RNAs were considered to target a gene if they lay within coding sequences or exons (for CRISPR knockout plasmids) or within 1000 base pairs of a transcription start site (for CRISPR activation plasmids).

**Figure S5.**
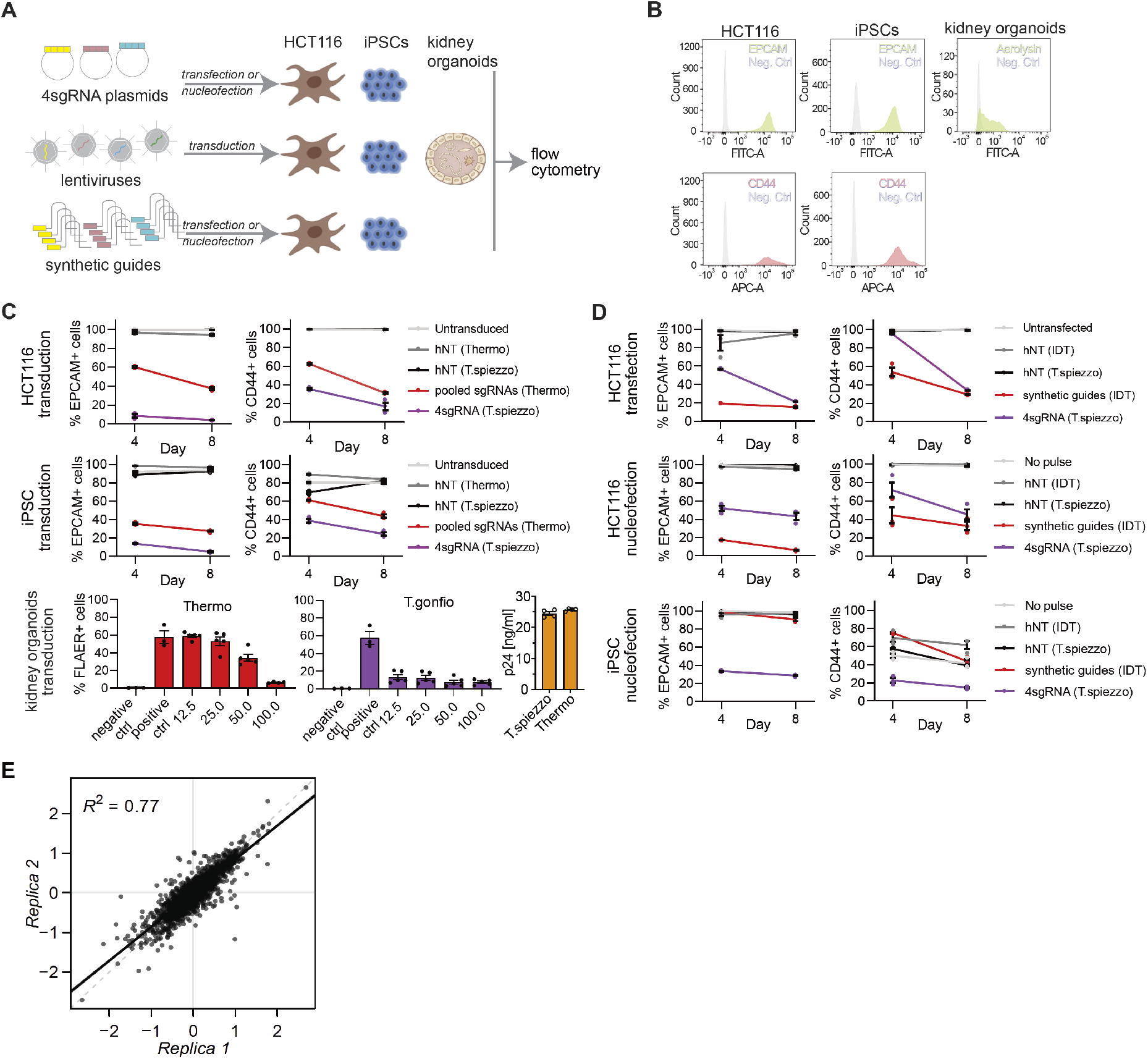
Benchmarking of 4sgRNA ablation plasmids in cells and organoids. **A**, Schematic of the experiment. 4sgRNA plasmids, synthetic guide RNAs, or lentivirally packaged sgRNAs were either transfected, nucleofected or transduced into Cas9 expressing HCT116, iPSCs and nephron progenitor cells which were matured to kidney organoids. **B**, Flow-cytometry histograms of Cas9- expressing HCT116 and iPSC cells stained with antibodies to EPCAM (*green; top left and middle*) or CD44 (*red; bottom left and middle*), and single-cell-dissociated kidney organoids stained with fluorescently labelled aerolysin (*green, top right*). **C**, upper and middle, Percentages of EPCAM^+^ or CD44^+^ HCT116-Cas9 (upper) and iPSC-iCas9 cells (middle) transduced with lentiviruses carrying the T.spiezzo 4sgRNA vector or a mixture of four individual lentiviruses (*Thermo*) targeting *EPCAM* or *CD44* or non-targeting (hNT) controls at 4 and 8 days post transduction (*n*=3; error bars represent S.E.M.); bottom (left and middle), Percentage of fluorescently labelled aerolysin (FLAER) positive cells dissociated from kidney organoids transduced with T.spiezzo lentiviruses or four individual lentiviruses (*Thermo*) targeting *PIGA* at increasing viral volumes compared to the unstained (negative ctrl) and untransduced (positive Ctrl) controls (*n*=4-5; error bars represent S.E.M.). bottom (right), p24 ELISA of supernatants containing *T.spiezzo* or *Thermo* lentiviruses targeting *PIGA* (*n*=4; error bars represent SEM). **D**, Percentages of EPCAM^+^ or CD44^+^ HCT116-Cas9 cells transfected (upper panels), electroporated (middle panels) or iPSC-iCas9 cells nucleofected (lower panels) with *T.spiezzo* (5 µg) or 4 synthetic guide RNAs (*Integrated DNA Technologies,* 10 µM) targeting *EPCAM* or *CD44* at 4 and 8 days post transfection (*n*=3; error bars: S.E.M.) **E**, Duplicate correlation across samples from the primary TF screen.

**Figure S6.**
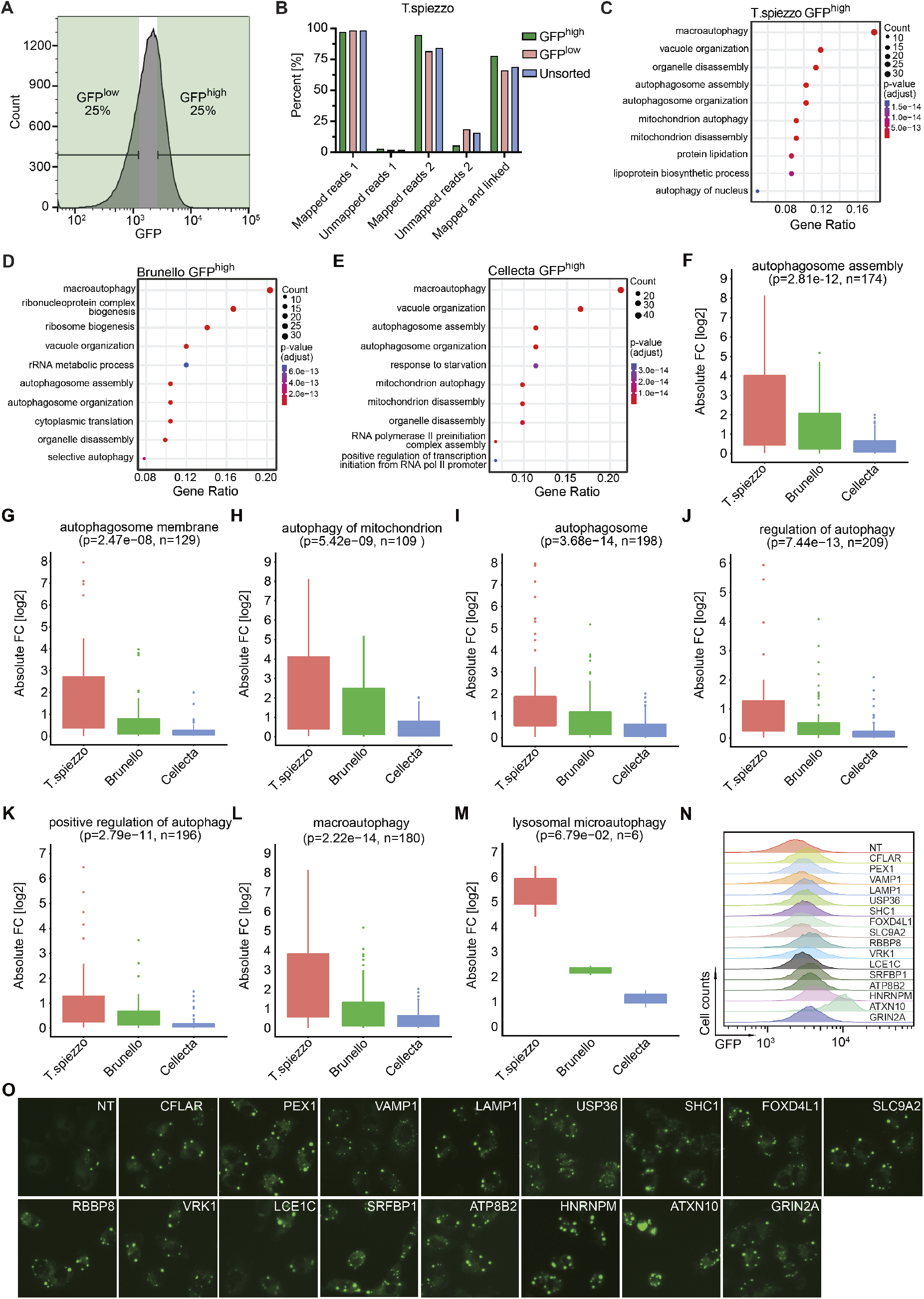
Template switching in pooled T.spiezzo and autophagy-related genes with higher fold changes compared to existing CRISPR knockout libraries. **A**, Histogram showing gating strategy to isolate GFP^high^ and GFP^low^ (upper and lower quartile of GFP fluorescence, respectively) cell populations. **B**, Percentages of sequencing reads in GFP^high^, GFP^low^ and unsorted samples from the T.spiezzo pooled screen that correctly aligned to sgRNA2 (mapped reads 1) or sgRNA3 (mapped reads 2), that did not align to sgRNA2 (unmapped reads 1) or sgRNA3 (unmapped reads 2) and those that aligned and had the correct linkage between sgRNA2 and sgRNA3 (mapped and linked). **C**-**E**, Overrepresentation analysis of the top 200 genes enriched in GFP^high^ cell populations from the T.spiezzo (**C**), Brunello (**D**) and Cellecta (**E**) screens. Gene counts and adjusted p-value are represented in each figure. The 10 most significant GO biological processes are shown. **F-M**, Autophagy-related gene sets including autophagosome assembly (GO:0000045, n=174, **F**), autophagosome membrane (GO:0000421, n=129, **G**), autophagy of mitochondrion (GO:0000422, n=109, **H**), autophagosome (GO:0005776, n=198, **I**), regulation of autophagy (GO:0010506, n=209, **J**), positive regulation of autophagy (GO:0010508, n=196, **K**), macroautophagy (GO:0016236, n=180, **L**), and lysosomal microautophagy (GO:0016237, n=6, **M**) using absolute Log2 fold changes in GFP^high^ cell populations from the T.spiezzo, Brunello and Cellecta screens. p value was determined by one-way ANOVA. **N**, An example of flow-cytometry histograms of GFP-SQSTM1 intensity of H4- Cas9-GFP-SQSTM1-cells transduced with T.spiezzo 4sgRNA lentivirus against nontargeting (NT) control or each of the 16 genes selected for validation. N = 3 biological repeats. **O**, An example of GFP-SQSTM1 puncta in H4-Cas9-GFP-SQSTM1-cells transduced with T.spiezzo 4sgRNA lentivirus against NT or each of the 16 genes selected for validation. N = 3 biological repeats.

**Figure S7.**
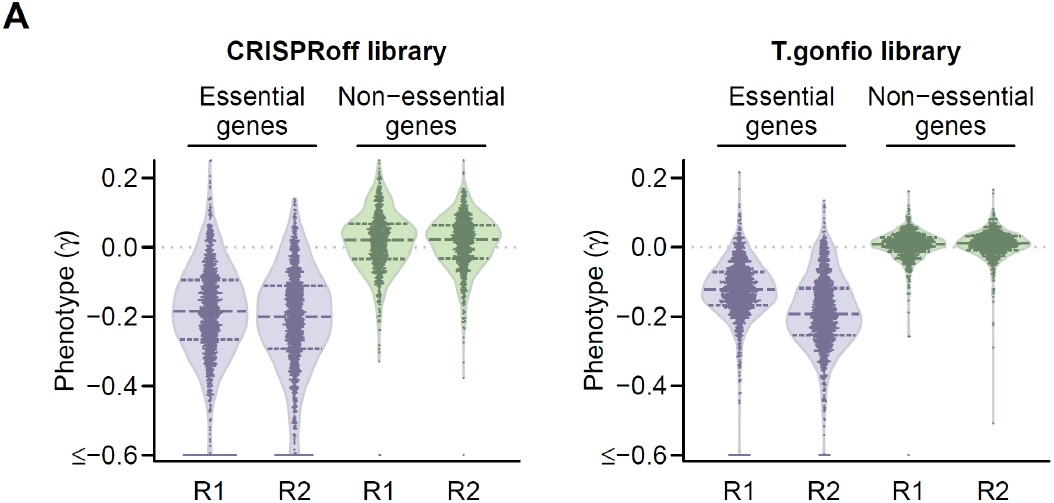
Genome-wide CRISPRoff screens. **A**, Violin plots of the phenotype scores (γ) for essential and non-essential genes in the pair-wise genome-wide epigenetic silencing screens. R1 and R2 represent each individual biological repeat of the screens.

## Supplementary information

### 1. Sequence of the empty pYJA5 vector

CAATTGGAGAAGTGAATTATATAAATATAAAGTAGTAAAAATTGAACCATTAGGAGTAGCACCCAC CAAGGCAAAGAGAAGAGTGGTGCAGAGAGAAAAAAGAGCAGTGGGAATAGGAGCTTTGTTCCTT GGGTTCTTGGGAGCAGCAGGAAGCACTATGGGCGCAGCGTCAATGACGCTGACGGTACAGGCC AGACAATTATTGTCTGGTATAGTGCAGCAGCAGAACAATTTGCTGAGGGCTATTGAGGCGCAACA GCATCTGTTGCAACTCACAGTCTGGGGCATCAAGCAGCTCCAGGCAAGAATCCTGGCTGTGGAA AGATACCTAAAGGATCAACAGCTCCTGGGGATTTGGGGTTGCTCTGGAAAACTCATTTGCACCAC TGCTGTGCCTTGGAATGCTAGTTGGAGTAATAAATCTCTGGAACAGATTTGGAATCACACGACCT GGATGGAGTGGGACAGAGAAATTAACAATTACACAAGCTTAATACACTCCTTAATTGAAGAATCG CAAAACCAGCAAGAAAAGAATGAACAAGAATTATTGGAATTAGATAAATGGGCAAGTTTGTGGAA TTGGTTTAACATAACAAATTGGCTGTGGTATATAAAATTATTCATAATGATAGTAGGAGGCTTGGT AGGTTTAAGAATAGTTTTTGCTGTACTTTCTATAGTGAATAGAGTTAGGCAGGGATATTCACCATT ATCGTTTCAGACCCACCTCCCAACCCCGAGGGGACCCGACAGGCCCGAAGGAATAGAAGAAGA AGGTGGAGAGAGAGACAGAGACAGATCCATTCGATTAGTGAACGGATCGGCACTGCGTGCGCC AATTCTGCAGACAAATGGCAGTATTCATCCACAATTTTAAAAGAAAAGGGGGGATTGGGGGGTAC AGTGCAGGGGAAAGAATAGTAGACATAATAGCAACAGACATACAAACTAAAGAATTACAAAAACA AATTACAAAAATTCAAAATTTTCGGGTTTATTACAGGGACAGCAGAGATCCAGTTTGGTTAGTACC GGGCCCTACGCGTTACTTAACCCTAGAAAGATAATCATATTGTGACGTACGTTAAAGATAATCAT GCGTAAAATTGACGCATGTGTTTTATCGGTCTGTATATCGAGGTTTATTTATTAATTTGAATAGATA TTAAGTTTTATTATATTTACACTTACATACTAATAATAAATTCAACAAACAATTTATTTATGTTTATTT ATTTATTAAAAAAAAACAAAAACTCAAAATTTCTTCTATAAAGTAACAAAGCaaaaaaaGCACCGACT CGGTGCCACTTTTTCAAGTTGATAACGGACTAGCCTTATTTCAACTTGCTACAGCATTTCTGCTGT AGCTCTGAAACccGTCTTCTTACCAATGCTTAATCAGTGAGGCACCTATCTCAGCGATCTGTCTAT TTCGTTCATCCATAGTTGCCTGACTCCCCGTCGTGTAGATAACTACGATACGGGAGGGCTTACCA TCTGGCCCCAGTGCTGCAATGATACCGCGAGACCCACGCTCACCGGCTCCAGATTTATCAGCAA TAAACCAGCCAGCCGGAAGGGCCGAGCGCAGAAGTGGTCCTGCAACTTTATCCGCCTCCATCCA GTCTATTAATTGTTGCCGGGAAGCTAGAGTAAGTAGTTCGCCAGTTAATAGTTTGCGCAACGTTG TTGCCATTGCTACAGGCATCGTGGTGTCACGCTCGTCGTTTGGTATGGCTTCATTCAGCTCCGGT TCCCAACGATCAAGGCGAGTTACATGATCCCCCATGTTGTGCAAAAAAGCGGTTAGCTCCTTCG GTCCTCCGATCGTTGTCAGAAGTAAGTTGGCCGCAGTGTTATCACTCATGGTTATGGCAGCACTG CATAATTCTCTTACTGTCATGCCATCCGTAAGATGCTTTTCTGTGACTGGTGAGTACTCAACCAAG TCATTCTGAGAATAGTGTATGCGGCGACCGAGTTGCTCTTGCCCGGCGTCAATACGGGATAATA CCGCGCCACATAGCAGAACTTTAAAAGTGCTCATCATTGGAAAACGTTCTTCGGGGCGAAAACTC TCAAGGATCTTACCGCTGTTGAGATCCAGTTCGATGTAACCCACTCGTGCACCCAACTGATCTTC AGCATCTTTTACTTTCACCAGCGTTTCTGGGTGAGCAAAAACAGGAAGGCAAAATGCCGCAAAAA AGGGAATAAGGGCGACACGGAAATGTTGAATACTCATACTCTTCCTTTTTCAATATTATTGAAGCA TTTATCAGGGTTATTGTCTCATGAGCGGATACATATTTGAATGTATTTAGAAAAATAAACAAATAG GGGTTCCGCGGAAGACccCggtgtttcgtcctttccacaagatatataaagccaagaaatcgaaatactttcaagttacggtaagca tatgatagtccattttaaaacataattttaaaactgcaaactacccaagaaattattactttctacgtcacgtattttgtactaatatctttgtgtttacagt caaattaattctaattatctctctaacagccttgtatcgtatatgcaaatatgaaggaatcatgggaaataggccctcTTCCTGCCCGACC TTGGgGATCCAATTCTACCGGGTAGGGGAGGCGCTTTTCCCAAGGCAGTCTGGAGCATGCGCTT TAGCAGCCCCGCTGGGCACTTGGCGCTACACAAGTGGCCTCTGGCCTCGCACACATTCCACATC CACCGGTAGGCGCCAACCGGCTCCGTTCTTTGGTGGCCCCTTCGCGCCACCTTCTACTCCTCCC CTAGTCAGGAAGTTCCCCCCCGCCCCGCAGCTCGCGTCGTGCAGGACGTGACAAATGGAAGTA GCAGTCTCACTAGTCTCGTGCAGATGGACAGCACCGCTGAGCAATGGAAGCGGGTAGGCCTTT GGGGCAGCGGCCAATAGCAGCTTTGCTCCTTCGCTTTCTGGGCTCAGAGGCTGGGAAGGGGTG GGTCCGGGGGCGGGCTCAGGGGCGGGCTCAGGGGCGGGGCGGGCGCCCGAAGGTCCTCCG GAGGCCCGGCATTCTGCACGCTTCAAAAGCGCACGTCTGCCGCGCTGTTCTCCTCTTCCTCATC TCCGGGCCTTTCGACCTGCATCCATCTAGATCTCGAGCAGCTGAAGCTTACCATGACCGAGTAC AAGCCCACGGTGCGCCTCGCCACCCGCGACGACGTCCCCAGGGCCGTACGCACCCTCGCCGC CGCGTTCGCCGACTACCCCGCCACGCGCCACACCGTCGATCCGGACCGCCACATCGAGCGGGT CACCGAGCTGCAAGAACTCTTCCTCACGCGCGTCGGGCTCGACATCGGCAAGGTGTGGGTCGC GGACGACGGCGCCGCGGTGGCGGTCTGGACCACGCCGGAGAGCGTCGAAGCGGGGGCGGTG TTCGCCGAGATCGGCCCGCGCATGGCCGAGTTGAGCGGTTCCCGGCTGGCCGCGCAGCAACA GATGGAAGGCCTCCTGGCGCCGCACCGGCCCAAGGAGCCCGCGTGGTTCCTGGCCACCGTCG GgGTCTCGCCCGACCACCAGGGCAAGGGTCTGGGCAGCGCCGTCGTGCTCCCCGGAGTGGAG GCGGCCGAGCGCGCCGGGGTGCCCGCCTTCCTGGAGACCTCCGCGCCCCGCAACCTCCCCTT CTACGAGCGGCTCGGCTTCACCGTCACCGCCGACGTCGAGGTGCCCGAAGGACCGCGCACCT GGTGCATGACCCGCAAGCCCGGTGCCGGCGGCGGGTCCGGAGGAGAGGGCAGAGGAAGTCTC CTAACATGCGGTGACGTGGAGGAGAATCCTGGCCCAATGAGCGAGCTGATTAAGGAGAACATGC ACATGAAGCTGTACATGGAGGGCACCGTGGACAACCATCACTTCAAGTGCACATCCGAGGGCGA AGGCAAGCCCTACGAGGGCACCCAGACCATGAGAATCAAGGTGGTCGAGGGCGGCCCTCTCCC CTTCGCCTTCGACATCCTGGCTACTAGCTTCCTCTACGGCAGCAAGACCTTCATCAACCACACCC AGGGCATCCCCGACTTCTTCAAGCAGTCCTTCCCTGAGGGCTTCACATGGGAGAGAGTCACCAC ATACGAGGACGGGGGCGTGCTGACCGCTACCCAGGACACCAGCCTCCAGGACGGCTGCCTCAT CTACAACGTCAAGATCAGAGGGGTGAACTTCACATCCAACGGCCCTGTGATGCAGAAGAAAACA CTCGGCTGGGAGGCCTTCACCGAGACtCTGTACCCCGCTGACGGCGGCCTGGAAGGCAGAAAC GACATGGCCCTGAAGCTCGTGGGCGGGAGCCATCTGATCGCAAACATCAAGACCACATATAGAT CCAAGAAACCCGCTAAGAACCTCAAGATGCCTGGCGTCTACTATGTGGACTACAGACTGGAAAG AATCAAGGAGGCCAACAACGAGACCTACGTCGAGCAGCACGAGGTGGCAGTGGCCAGATACTG CGACCTCCCTAGCAAACTGGGGCACAAGCTTAATTGAGCGGCCGCTAGGTACCTTTAAGACCAA TGACTTACAAGGCAGCTGTAGATCTTAGCCACTTTTTAAAAGAAAAGGGGGGACTGGAAGGGCT AATTCACTCCCAAAGAAGTCAAGATCTGCTTTTTGCCTGTACTGGGTCTCTCTGGTTAGACCAGA GTCTCTCTGGTTAGACCAGATCTGAGCCTGGGAGCTCTCTGGCTAACTAGGGAACCCACTGCTT AAGCCTCAATAAAGCTTGCCTTGAGTGCTTCAAGTAGTGTGTGCCCGTCTGTTGTGTGACTCTGG TAACTAGAGATCCCTCAGACCCTTTTAGTCAGTGTGGAAAATCTCTAGCAGTTTAAACCCGCTGA TCAGCCTCGACTGTGCCTTCTAGTTGCCAGCCATCTGTTGTTTGCCCCTCCCCCGTGCCTTCCTT GACCCTGGAAGGTGCCACTCCCACTGTCCTTTCCTAATAAAATGAGGAAATTGCATCGCATTGTC TGAGTAGGTGTCATTCTATTCTGGGGGGTGGGGTGGGGCAGGACAGCAAGGGGGAGGATTGGG AAGTCAATAGCAGGCATGCTGGGGATGCGGTGGGCTCTATGGGCGGCCGTTAATGATATCTATA ACAAGAAAATATATATATAATAAGTTATCACGTAAGTAGAACATGAAATAACAATATAATTATCGTA TGAGTTAAATCTTAAAAGTCACGTAAAAGATAATCATGCGTCATTTTGACTCACGCGGTCGTTATA GTTCAAAATCAGTGACACTTACCGCATTGACAAGCACGCCTCACGGGAGCTCCAAGCGGCGACT GAGATGTCCTAAATGCACAGCGACGGATTCGCGCTATTTAGAAAGAGAGAGCAATATTTCAAGAA TGCATGCGTCAATTTTACGCAGACTATCTTTCTAGGGTTAAATTAAGCTTTTGTTCCCTTTAGTGA GGGTTAATTGCGCGCTTGGCGTAATCATGGTCATAGCTGTTTCCTGTGTGAAATTGTTATCCGCT CACAATTCCACACAACATACGAGCCGGgAGCATAAAGTGTAAAGCCTGGGGTGCCTAATGAGTG AGCTAACTCACATTAATTGCGTTGCGCTCACTGCCCGCTTTCCAGTCGGGAAACCTGTCGTGCCA GCTGCATTAATGAATCGGCCAACGCGCGGGGAGAGGCGGTTTGCGTATTGGGCGCTCTTCCGC TTCCTCGCTCACTGACTCGCTGCGCTCGGTCGTTCGGCTGCGGCGAGCGGTATCAGCTCACTCA AAGGCGGTAATACGGTTATCCACAGAATCAGGGGATAACGCAGGAAAGAACATGTGAGCAAAAG GCCAGCAAAAGGCCAGGAACCGTAAAAAGGCCGCGTTGCTGGCGTTTTTCCATAGGCTCCGCC CCCCTGACGAGCATCACAAAAATCGACGCTCAAGTCAGAGGTGGCGAAACCCGACAGGACTATA AAGATACCAGGCGTTTCCCCCTGGAAGCTCCCTCGTGCGCTCTCCTGTTCCGACCCTGCCGCTT ACCGGATACCTGTCCGCCTTTCTCCCTTCGGGAAGCGTGGCGCTTTCTCATAGCTCACGCTGTA GGTATCTCAGTTCGGTGTAGGTCGTTCGCTCCAAGCTGGGCTGTGTGCACGAACCCCCCGTTCA GCCCGACCGCTGCGCCTTATCCGGTAACTATCGTCTTGAGTCCAACCCGGTAAGACACGACTTA TCGCCACTGGCAGCAGCCACTGGTAACAGGATTAGCAGAGCGAGGTATGTAGGCGGTGCTACA GAGTTCTTGAAGTGGTGGCCTAACTACGGCTACACTAGAAGaACAGTATTTGGTATCTGCGCTCT GCTGAAGCCAGTTACCTTCGGAAAAAGAGTTGGTAGCTCTTGATCCGGCAAACAAACCACCGCT GGTAGCGGTGGTTTTTTTGTTTGCAAGCAGCAGATTACGCGCAGAAAAAAAGGATCTCAAGAAGA TCCTTTGATCTTTTCTACGGGGTCTGACGCTCAGTGGAACGAAAACTCACGTTAAGGGATTTTGG TCATGAGCGGATACATATTTGAATGTATTTAGAAAAATAAACAAATAGGGGTTCCGCGCACATTTC CCCGAAAAGTGCCACCTAAATTGTAAGCGTTAATATTTTGTTAAAATTCGCGTTAAATTTTTGTTAA ATCAGCTCATTTTTTAACCAATAGGCCGAAATCGGCAAAATCCCTTATAAATCAAAAGAATAGACC GAGATAGGGTTGAGTGTTGTTCCAGTTTGGAACAAGAGTCCACTATTAAAGAACGTGGACTCCAA CGTCAAAGGGCGAAAAACCGTCTATCAGGGCGATGGCCCACTACGTGAACCATCACCCTAATCA AGTTTTTTGGGGTCGAGGTGCCGTAAAGCACTAAATCGGAACCCTAAAGGGAGCCCCCGATTTA GAGCTTGACGGGGAAAGCCGGCGAACGTGGCGAGAAAGGAAGGGAAGAAAGCGAAAGGAGCG GGCGCTAGGGCGCTGGCAAGTGTAGCGGTCACGCTGCGCGTAACCACCACACCCGCCGCGCTT AATGCGCCGCTACAGGGCGCGTCCCATTCGCCATTCAGGCTGCGCAACTGTTGGGAAGGGCGA TCGGTGCGGGCCTCTTCGCTATTACGCCAGCTGGCGAAAGGGGGATGTGCTGCAAGGCGATTA AGTTGGGTAACGCCAGGGTTTTCCCAGTCACGACGTTGTAAAACGACGGCCAGTGAGCGCGCG TAATACGACTCACTATAGGGCGAATTGACTAGTTATTAATAGTAATCAATTACGGGGTCATTAGTT CATAGCCCATATATGGAGTTCCGCGTTACATAACTTACGGTAAATGGCCCGCCTGGCTGACCGC CCAACGACCCCCGCCCATTGACGTCAATAATGACGTATGTTCCCATAGTAACGCCAATAGGGACT TTCCATTGACGTCAATGGGTGGAGTATTTACGGTAAACTGCCCACTTGGCAGTACATCAAGTGTA TCATATGCCAAGTACGCCCCCTATTGACGTCAATGACGGTAAATGGCCCGCCTGGCATTATGCC CAGTACATGACCTTATGGGACTTTCCTACTTGGCAGTACATCTACGTATTAGTCATCGCTATTACC ATGGTGATGCGGTTTTGGCAGTACATCAATGGGCGTGGATAGCGGTTTGACTCACGGGGATTTC CAAGTCTCCACCCCATTGACGTCAATGGGAGTTTGTTTTGGCACCAAAATCAACGGGACTTTCCA AAATGTCGTAACAACTCCGCCCCATTGACGCAAATGGGCGGTAGGCGTGTACGGTGGGAGGTCT ATATAAGCAGCGCGTTTTGCCTGTACTGGGTCTCTCTGGTTAGACCAGATCTGAGCCTGGGAGC TCTCTGGCTAACTAGGGAACCCACTGCTTAAGCCTCAATAAAGCTTGCCTTGAGTGCTTCAAGTA GTGTGTGCCCGTCTGTTGTGTGACTCTGGTAACTAGAGATCCCTCAGACCCTTTTAGTCAGTGTG GAAAATCTCTAGCAGTGGCGCCCGAACAGGGACTTGAAAGCGAAAGGGAAACCAGAGGAGCTC TCTCGACGCAGGACTCGGCTTGCTGAAGCGCGCACGGCAAGAGGCGAGGGGCGGCGACTGGT GAGTACGCCAAAAATTTTGACTAGCGGAGGCTAGAAGGAGAGAGATGGGTGCGAGAGCGTCAG TATTAAGCGGGGGAGAATTAGATCGCGATGGGAAAAAATTCGGTTAAGGCCAGGGGGAAAGAAA AAATATAAATTAAAACATATAGTATGGGCAAGCAGGGAGCTAGAACGATTCGCAGTTAATCCTGG CCTGTTAGAAACATCAGAAGGCTGTAGACAAATACTGGGACAGCTACAACCATCCCTTCAGACAG GATCAGAAGAACTTAGATCATTATATAATACAGTAGCAACCCTCTATTGTGTGCATCAAAGGATAG AGATAAAAGACACCAAGGAAGCTTTAGACAAGATAGAGGAAGAGCAAAACAAAAGTAAGACCAC CGCACAGCAAGCGGCCGGCCGCTGATCTTCAGACCTGGAGGAGGAGATATGAGGGA

### 2. Sequence of the 4sgRNA-pYJA5 plasmid (N20 indicates sgRNA sequence)

CAATTGGAGAAGTGAATTATATAAATATAAAGTAGTAAAAATTGAACCATTAGGAGTAGCACCCAC CAAGGCAAAGAGAAGAGTGGTGCAGAGAGAAAAAAGAGCAGTGGGAATAGGAGCTTTGTTCCTT GGGTTCTTGGGAGCAGCAGGAAGCACTATGGGCGCAGCGTCAATGACGCTGACGGTACAGGCC AGACAATTATTGTCTGGTATAGTGCAGCAGCAGAACAATTTGCTGAGGGCTATTGAGGCGCAACA GCATCTGTTGCAACTCACAGTCTGGGGCATCAAGCAGCTCCAGGCAAGAATCCTGGCTGTGGAA AGATACCTAAAGGATCAACAGCTCCTGGGGATTTGGGGTTGCTCTGGAAAACTCATTTGCACCAC TGCTGTGCCTTGGAATGCTAGTTGGAGTAATAAATCTCTGGAACAGATTTGGAATCACACGACCT GGATGGAGTGGGACAGAGAAATTAACAATTACACAAGCTTAATACACTCCTTAATTGAAGAATCG CAAAACCAGCAAGAAAAGAATGAACAAGAATTATTGGAATTAGATAAATGGGCAAGTTTGTGGAA TTGGTTTAACATAACAAATTGGCTGTGGTATATAAAATTATTCATAATGATAGTAGGAGGCTTGGT AGGTTTAAGAATAGTTTTTGCTGTACTTTCTATAGTGAATAGAGTTAGGCAGGGATATTCACCATT ATCGTTTCAGACCCACCTCCCAACCCCGAGGGGACCCGACAGGCCCGAAGGAATAGAAGAAGA AGGTGGAGAGAGAGACAGAGACAGATCCATTCGATTAGTGAACGGATCGGCACTGCGTGCGCC AATTCTGCAGACAAATGGCAGTATTCATCCACAATTTTAAAAGAAAAGGGGGGATTGGGGGGTAC AGTGCAGGGGAAAGAATAGTAGACATAATAGCAACAGACATACAAACTAAAGAATTACAAAAACA AATTACAAAAATTCAAAATTTTCGGGTTTATTACAGGGACAGCAGAGATCCAGTTTGGTTAGTACC GGGCCCTACGCGTTACTTAACCCTAGAAAGATAATCATATTGTGACGTACGTTAAAGATAATCAT GCGTAAAATTGACGCATGTGTTTTATCGGTCTGTATATCGAGGTTTATTTATTAATTTGAATAGATA TTAAGTTTTATTATATTTACACTTACATACTAATAATAAATTCAACAAACAATTTATTTATGTTTATTT ATTTATTAAAAAAAAACAAAAACTCAAAATTTCTTCTATAAAGTAACAAAGCaaaaaaaGCACCGACT CGGTGCCACTTTTTCAAGTTGATAACGGACTAGCCTTATTTCAACTTGCTACAGCATTTCTGCTGT AGCTCTGAAACNNNNNNNNNNNNNNNNNNNNCgaggtacccaagcggcgcacaagctatataaacctgaaggaagt ctcaactttacacttaggtcaagttgcttatcgtactagagcttcagcaggaaatttaactaaaatctaatttaaccagcatagcaaatatcatttatt cccaaaatgctaaagtttgagataaacggacttgatttccggctgttttgacactatccagaatgccttgcagatgggtggggcatgctaaatact gcagaaaaaaaGCACCCGACTCGGGTGCCACTTTTTCAAGTTGTAAACGGACTAGCCTTATTTCAAC TTGCTATGCACTCTTGTGCTTAGCTCTGAAACNNNNNNNNNNNNNNNNNNNNCgggaaagagtggtctc atacagaacttataagattcccaaatccaaagacatttcacgtttatggtgatttcccagaacacatagcgacatgcaaatattgcagggcgcc actcccctgtccctcacagccatcttcctgccagggcgcacgcgcgctgggtgttcccgcctagtgacactgggcccgcgattccttggagcgg gttgatgacgtcagcgttcaaaaaaaGCAGCCGACTCGGCTGCCACTTTTTCAAGTTGTGTACGGACTAGCC TTATTTGAACTTGCTATGCAGCTTTCTGCTTAGCTCTCAAACNNNNNNNNNNNNNNNNNNNNCaaa caaggcttttctccaagggatatttatagtctcaaaacacacaattactttacagttagggtgagtttccttttgtgctgttttttaaaataataatttagta tttgtatctcttatagaaatccaagcctatcatgtaaaatgtagctagtattaaaaagaacagattatctgtcttttatcgcacattaagcctctatagtt actaggaaatattatatgcaaattaaccggggcaggggagtagccgagcttctcccacaagtctgtgcgagggggccggcgcgggcctaga gatggcggcgtcggatcaaaaaaattaggccacacgttcaagtgcagccacaggataaatttgcactgagcctgggtgggattcggactcga ccgcatagccttcaggagtgagttttgtgcaataccaaccgacgacttgaccctgccaagcggcaccagatttcttgcgtacgcgatcccctaa gccaaaggtggcactcaggggaagcgcaaactgccctgcaacgggagcgttggcttcatcgctactttgacccatggtttagttcctcaccttgt cgtattatactatgccgatatactatgccgatgattaattgtcaacaaaaaaaGCACCGACTCGGTGCCACTTTTTCAAGTT GATAACGGACTAGCCTTATTTAAACTTGCTATGCTGTTTCCAGCTTAGCTCTTAAACNNNNNNNNN NNNNNNNNNNNCggtgtttcgtcctttccacaagatatataaagccaagaaatcgaaatactttcaagttacggtaagcatatgatagtc cattttaaaacataattttaaaactgcaaactacccaagaaattattactttctacgtcacgtattttgtactaatatctttgtgtttacagtcaaattaatt ctaattatctctctaacagccttgtatcgtatatgcaaatatgaaggaatcatgggaaataggccctcTTCCTGCCCGACCTTGGgG ATCCAATTCTACCGGGTAGGGGAGGCGCTTTTCCCAAGGCAGTCTGGAGCATGCGCTTTAGCAG CCCCGCTGGGCACTTGGCGCTACACAAGTGGCCTCTGGCCTCGCACACATTCCACATCCACCG GTAGGCGCCAACCGGCTCCGTTCTTTGGTGGCCCCTTCGCGCCACCTTCTACTCCTCCCCTAGT CAGGAAGTTCCCCCCCGCCCCGCAGCTCGCGTCGTGCAGGACGTGACAAATGGAAGTAGCAGT CTCACTAGTCTCGTGCAGATGGACAGCACCGCTGAGCAATGGAAGCGGGTAGGCCTTTGGGGC AGCGGCCAATAGCAGCTTTGCTCCTTCGCTTTCTGGGCTCAGAGGCTGGGAAGGGGTGGGTCC GGGGGCGGGCTCAGGGGCGGGCTCAGGGGCGGGGCGGGCGCCCGAAGGTCCTCCGGAGGC CCGGCATTCTGCACGCTTCAAAAGCGCACGTCTGCCGCGCTGTTCTCCTCTTCCTCATCTCCGG GCCTTTCGACCTGCATCCATCTAGATCTCGAGCAGCTGAAGCTTACCATGACCGAGTACAAGCC CACGGTGCGCCTCGCCACCCGCGACGACGTCCCCAGGGCCGTACGCACCCTCGCCGCCGCGT TCGCCGACTACCCCGCCACGCGCCACACCGTCGATCCGGACCGCCACATCGAGCGGGTCACCG AGCTGCAAGAACTCTTCCTCACGCGCGTCGGGCTCGACATCGGCAAGGTGTGGGTCGCGGACG ACGGCGCCGCGGTGGCGGTCTGGACCACGCCGGAGAGCGTCGAAGCGGGGGCGGTGTTCGC CGAGATCGGCCCGCGCATGGCCGAGTTGAGCGGTTCCCGGCTGGCCGCGCAGCAACAGATGG AAGGCCTCCTGGCGCCGCACCGGCCCAAGGAGCCCGCGTGGTTCCTGGCCACCGTCGGgGTCT CGCCCGACCACCAGGGCAAGGGTCTGGGCAGCGCCGTCGTGCTCCCCGGAGTGGAGGCGGCC GAGCGCGCCGGGGTGCCCGCCTTCCTGGAGACCTCCGCGCCCCGCAACCTCCCCTTCTACGAG CGGCTCGGCTTCACCGTCACCGCCGACGTCGAGGTGCCCGAAGGACCGCGCACCTGGTGCAT GACCCGCAAGCCCGGTGCCGGCGGCGGGTCCGGAGGAGAGGGCAGAGGAAGTCTCCTAACAT GCGGTGACGTGGAGGAGAATCCTGGCCCAATGAGCGAGCTGATTAAGGAGAACATGCACATGA AGCTGTACATGGAGGGCACCGTGGACAACCATCACTTCAAGTGCACATCCGAGGGCGAAGGCA AGCCCTACGAGGGCACCCAGACCATGAGAATCAAGGTGGTCGAGGGCGGCCCTCTCCCCTTCG CCTTCGACATCCTGGCTACTAGCTTCCTCTACGGCAGCAAGACCTTCATCAACCACACCCAGGG CATCCCCGACTTCTTCAAGCAGTCCTTCCCTGAGGGCTTCACATGGGAGAGAGTCACCACATAC GAGGACGGGGGCGTGCTGACCGCTACCCAGGACACCAGCCTCCAGGACGGCTGCCTCATCTAC AACGTCAAGATCAGAGGGGTGAACTTCACATCCAACGGCCCTGTGATGCAGAAGAAAACACTCG GCTGGGAGGCCTTCACCGAGACtCTGTACCCCGCTGACGGCGGCCTGGAAGGCAGAAACGACA TGGCCCTGAAGCTCGTGGGCGGGAGCCATCTGATCGCAAACATCAAGACCACATATAGATCCAA GAAACCCGCTAAGAACCTCAAGATGCCTGGCGTCTACTATGTGGACTACAGACTGGAAAGAATC AAGGAGGCCAACAACGAGACCTACGTCGAGCAGCACGAGGTGGCAGTGGCCAGATACTGCGAC CTCCCTAGCAAACTGGGGCACAAGCTTAATTGAGCGGCCGCTAGGTACCTTTAAGACCAATGAC TTACAAGGCAGCTGTAGATCTTAGCCACTTTTTAAAAGAAAAGGGGGGACTGGAAGGGCTAATTC ACTCCCAAAGAAGTCAAGATCTGCTTTTTGCCTGTACTGGGTCTCTCTGGTTAGACCAGAGTCTC TCTGGTTAGACCAGATCTGAGCCTGGGAGCTCTCTGGCTAACTAGGGAACCCACTGCTTAAGCC TCAATAAAGCTTGCCTTGAGTGCTTCAAGTAGTGTGTGCCCGTCTGTTGTGTGACTCTGGTAACT AGAGATCCCTCAGACCCTTTTAGTCAGTGTGGAAAATCTCTAGCAGTTTAAACCCGCTGATCAGC CTCGACTGTGCCTTCTAGTTGCCAGCCATCTGTTGTTTGCCCCTCCCCCGTGCCTTCCTTGACCC TGGAAGGTGCCACTCCCACTGTCCTTTCCTAATAAAATGAGGAAATTGCATCGCATTGTCTGAGT AGGTGTCATTCTATTCTGGGGGGTGGGGTGGGGCAGGACAGCAAGGGGGAGGATTGGGAAGTC AATAGCAGGCATGCTGGGGATGCGGTGGGCTCTATGGGCGGCCGTTAATGATATCTATAACAAG AAAATATATATATAATAAGTTATCACGTAAGTAGAACATGAAATAACAATATAATTATCGTATGAGT TAAATCTTAAAAGTCACGTAAAAGATAATCATGCGTCATTTTGACTCACGCGGTCGTTATAGTTCA AAATCAGTGACACTTACCGCATTGACAAGCACGCCTCACGGGAGCTCCAAGCGGCGACTGAGAT GTCCTAAATGCACAGCGACGGATTCGCGCTATTTAGAAAGAGAGAGCAATATTTCAAGAATGCAT GCGTCAATTTTACGCAGACTATCTTTCTAGGGTTAAATTAAGCTTTTGTTCCCTTTAGTGAGGGTT AATTGCGCGCTTGGCGTAATCATGGTCATAGCTGTTTCCTGTGTGAAATTGTTATCCGCTCACAA TTCCACACAACATACGAGCCGGgAGCATAAAGTGTAAAGCCTGGGGTGCCTAATGAGTGAGCTA ACTCACATTAATTGCGTTGCGCTCACTGCCCGCTTTCCAGTCGGGAAACCTGTCGTGCCAGCTG CATTAATGAATCGGCCAACGCGCGGGGAGAGGCGGTTTGCGTATTGGGCGCTCTTCCGCTTCCT CGCTCACTGACTCGCTGCGCTCGGTCGTTCGGCTGCGGCGAGCGGTATCAGCTCACTCAAAGG CGGTAATACGGTTATCCACAGAATCAGGGGATAACGCAGGAAAGAACATGTGAGCAAAAGGCCA GCAAAAGGCCAGGAACCGTAAAAAGGCCGCGTTGCTGGCGTTTTTCCATAGGCTCCGCCCCCCT GACGAGCATCACAAAAATCGACGCTCAAGTCAGAGGTGGCGAAACCCGACAGGACTATAAAGAT ACCAGGCGTTTCCCCCTGGAAGCTCCCTCGTGCGCTCTCCTGTTCCGACCCTGCCGCTTACCGG ATACCTGTCCGCCTTTCTCCCTTCGGGAAGCGTGGCGCTTTCTCATAGCTCACGCTGTAGGTATC TCAGTTCGGTGTAGGTCGTTCGCTCCAAGCTGGGCTGTGTGCACGAACCCCCCGTTCAGCCCGA CCGCTGCGCCTTATCCGGTAACTATCGTCTTGAGTCCAACCCGGTAAGACACGACTTATCGCCA CTGGCAGCAGCCACTGGTAACAGGATTAGCAGAGCGAGGTATGTAGGCGGTGCTACAGAGTTCT TGAAGTGGTGGCCTAACTACGGCTACACTAGAAGaACAGTATTTGGTATCTGCGCTCTGCTGAAG CCAGTTACCTTCGGAAAAAGAGTTGGTAGCTCTTGATCCGGCAAACAAACCACCGCTGGTAGCG GTGGTTTTTTTGTTTGCAAGCAGCAGATTACGCGCAGAAAAAAAGGATCTCAAGAAGATCCTTTG ATCTTTTCTACGGGGTCTGACGCTCAGTGGAACGAAAACTCACGTTAAGGGATTTTGGTCATGAG CGGATACATATTTGAATGTATTTAGAAAAATAAACAAATAGGGGTTCCGCGCACATTTCCCCGAAA AGTGCCACCTAAATTGTAAGCGTTAATATTTTGTTAAAATTCGCGTTAAATTTTTGTTAAATCAGCT CATTTTTTAACCAATAGGCCGAAATCGGCAAAATCCCTTATAAATCAAAAGAATAGACCGAGATAG GGTTGAGTGTTGTTCCAGTTTGGAACAAGAGTCCACTATTAAAGAACGTGGACTCCAACGTCAAA GGGCGAAAAACCGTCTATCAGGGCGATGGCCCACTACGTGAACCATCACCCTAATCAAGTTTTTT GGGGTCGAGGTGCCGTAAAGCACTAAATCGGAACCCTAAAGGGAGCCCCCGATTTAGAGCTTGA CGGGGAAAGCCGGCGAACGTGGCGAGAAAGGAAGGGAAGAAAGCGAAAGGAGCGGGCGCTAG GGCGCTGGCAAGTGTAGCGGTCACGCTGCGCGTAACCACCACACCCGCCGCGCTTAATGCGCC GCTACAGGGCGCGTCCCATTCGCCATTCAGGCTGCGCAACTGTTGGGAAGGGCGATCGGTGCG GGCCTCTTCGCTATTACGCCAGCTGGCGAAAGGGGGATGTGCTGCAAGGCGATTAAGTTGGGTA ACGCCAGGGTTTTCCCAGTCACGACGTTGTAAAACGACGGCCAGTGAGCGCGCGTAATACGACT CACTATAGGGCGAATTGACTAGTTATTAATAGTAATCAATTACGGGGTCATTAGTTCATAGCCCAT ATATGGAGTTCCGCGTTACATAACTTACGGTAAATGGCCCGCCTGGCTGACCGCCCAACGACCC CCGCCCATTGACGTCAATAATGACGTATGTTCCCATAGTAACGCCAATAGGGACTTTCCATTGAC GTCAATGGGTGGAGTATTTACGGTAAACTGCCCACTTGGCAGTACATCAAGTGTATCATATGCCA AGTACGCCCCCTATTGACGTCAATGACGGTAAATGGCCCGCCTGGCATTATGCCCAGTACATGA CCTTATGGGACTTTCCTACTTGGCAGTACATCTACGTATTAGTCATCGCTATTACCATGGTGATGC GGTTTTGGCAGTACATCAATGGGCGTGGATAGCGGTTTGACTCACGGGGATTTCCAAGTCTCCA CCCCATTGACGTCAATGGGAGTTTGTTTTGGCACCAAAATCAACGGGACTTTCCAAAATGTCGTA ACAACTCCGCCCCATTGACGCAAATGGGCGGTAGGCGTGTACGGTGGGAGGTCTATATAAGCA GCGCGTTTTGCCTGTACTGGGTCTCTCTGGTTAGACCAGATCTGAGCCTGGGAGCTCTCTGGCT AACTAGGGAACCCACTGCTTAAGCCTCAATAAAGCTTGCCTTGAGTGCTTCAAGTAGTGTGTGCC CGTCTGTTGTGTGACTCTGGTAACTAGAGATCCCTCAGACCCTTTTAGTCAGTGTGGAAAATCTC TAGCAGTGGCGCCCGAACAGGGACTTGAAAGCGAAAGGGAAACCAGAGGAGCTCTCTCGACGC AGGACTCGGCTTGCTGAAGCGCGCACGGCAAGAGGCGAGGGGCGGCGACTGGTGAGTACGCC AAAAATTTTGACTAGCGGAGGCTAGAAGGAGAGAGATGGGTGCGAGAGCGTCAGTATTAAGCGG GGGAGAATTAGATCGCGATGGGAAAAAATTCGGTTAAGGCCAGGGGGAAAGAAAAAATATAAAT TAAAACATATAGTATGGGCAAGCAGGGAGCTAGAACGATTCGCAGTTAATCCTGGCCTGTTAGAA ACATCAGAAGGCTGTAGACAAATACTGGGACAGCTACAACCATCCCTTCAGACAGGATCAGAAG AACTTAGATCATTATATAATACAGTAGCAACCCTCTATTGTGTGCATCAAAGGATAGAGATAAAAG ACACCAAGGAAGCTTTAGACAAGATAGAGGAAGAGCAAAACAAAAGTAAGACCACCGCACAGCA AGCGGCCGGCCGCTGATCTTCAGACCTGGAGGAGGAGATATGAGGGA

Four sgRNA primer sequence (5’-3’, N20 in sgRNA1 primer Fwd, sgRNA2 primer Fwd, sgRNA3 primer Fwd is exactly the sgRNA sequence, however, in sgRNA4 primer Rev it should be the reverse complement sequence of the sgRNA sequence):

sgRNA1 primer Fwd: ttgtggaaaggacgaaacaccGN20GTTTAAGAGCTAAGCTG sgRNA2 primer Fwd: cttggagaaaagccttgtttGN20GTTTGAGAGCTAAGCAGA sgRNA3 primer Fwd: gtatgagaccactctttcccGN20GTTTCAGAGCTAAGCACA sgRNA4 primer Rev: ATTTCTGCTGTAGCTCTGAAACN20Cgaggtacccaagcggc

Common primers sequences (5’-3’):

mU6 Rev: CAAACAAGGCTTTTCTCCAAGGG

M Rev: Cgggaaagagtggtctcataca

Constant template sequences (5’-3’)

C1 sequence GTTTAAGAGCTAAGCTGGAAACAGCATAGCAAGTTTAAATAAGGCTAGTCCGTTATCAACTTGAA AAAGTGGCACCGAGTCGGTGCtttttttgttgacaattaatcatcggcatagtatatcggcatagtataatacgacaaggtgagga actaaaccatgggtcaaagtagcgatgaagccaacgctcccgttgcagggcagtttgcgcttcccctgagtgccacctttggcttaggggatcg cgtacgcaagaaatctggtgccgcttggcagggtcaagtcgtcggttggtattgcacaaaactcactcctgaaggctatgcggtcgagtccga atcccacccaggctcagtgcaaatttatcctgtggctgcacttgaacgtgtggcctaatttttttgatccgacgccgccatctctaggcccgcgccg gccccctcgcacagacttgtgggagaagctcggctactcccctgccccggttaatttgcatataatatttcctagtaactatagaggcttaatgtgc gataaaagacagataatctgttctttttaatactagctacattttacatgataggcttggatttctataagagatacaaatactaaattattattttaaaa aacagcacaaaaggaaactcaccctaactgtaaagtaattgtgtgttttgagactataaatatcccttggagaaaagccttgtttG

M sequence GTTTGAGAGCTAAGCAGAAAGCTGCATAGCAAGTTCAAATAAGGCTAGTCCGTACACAACTTGAA AAAGTGGCAGCCGAGTCGGCTGCtttttttgaacgctgacgtcatcaacccgctccaaggaatcgcgggcccagtgtcactag gcgggaacacccagcgcgcgtgcgccctggcaggaagatggctgtgagggacaggggagtggcgccctgcaatatttgcatgtcgctatgt gttctgggaaatcaccataaacgtgaaatgtctttggatttgggaatcttataagttctgtatgagaccactctttcccG

C2s sequence GTTTCAGAGCTAAGCACAAGAGTGCATAGCAAGTTGAAATAAGGCTAGTCCGTTTACAACTTGAA AAAGTGGCACCCGAGTCGGGTGCtttttttctgcagtatttagcatgccccacccatctgcaaggcattctggatagtgtcaaaac agccggaaatcaagtccgtttatctcaaactttagcattttgggaataaatgatatttgctatgctggttaaattagattttagttaaatttcctgctgaa gctctagtacgataagcaacttgacctaagtgtaaagttgagacttccttcaggtttatatagcttgtgcgccgcttgggtacctcG

